# Different olfactory neuron classes use distinct temporal and molecular programs to complete synaptic development

**DOI:** 10.1101/2022.03.31.486637

**Authors:** Michael A. Aimino, Alison T. DePew, Lucas Restrepo, Timothy J. Mosca

**Author notes:** University of Massachusetts Chan Medical School, Department of Molecular, Cell, and Cancer Biology, Worcester MA, 01605.

## Abstract

Developing neurons must meet core molecular, cellular, and temporal requirements to ensure the correct formation of synapses, resulting in functional circuits. However, because of the vast diversity in neuronal class and function, it is unclear whether or not all neurons use the same organizational mechanisms to form synaptic connections and achieve functional and morphological maturation. Moreover, it remains unknown if neurons united in a common goal and comprising the same sensory circuit develop on similar timescales and using identical molecular approaches to ensure the formation of the correct number of synapses. To begin to answer these questions, we took advantage of the Drosophila antennal lobe, a model olfactory circuit with remarkable genetic access and synapse-level resolution. Using tissue-specific genetic labeling of active zones, we performed a quantitative analysis of synapse formation in multiple classes of neurons throughout development and adulthood. We found that olfactory receptor neurons (ORNs), projection neurons (PNs), and local interneurons (LNs) each have unique time-courses of synaptic development, addition, and refinement, demonstrating that each class follows a distinct developmental program. This raised the possibility that these classes may also have distinct molecular requirements for synapse formation. We genetically altered neuronal activity in each neuronal subtype and observed differing effects on synapse number based on the neuronal class examined. Silencing neuronal activity in ORNs, PNs, and LNs impaired synaptic development but only in ORNs did enhancing neuronal activity influence synapse formation. ORNs and LNs demonstrated similar impairment of synaptic development with enhanced activity of a master kinase, GSK-3β, suggesting that neuronal activity and GSK-3β kinase activity function in a common pathway. ORNs also, however, demonstrated impaired synaptic development with GSK-3β loss-of-function, suggesting additional activity-independent roles in development. Ultimately, our results suggest that the requirements for synaptic development are not uniform across all neuronal classes with considerable diversity existing in both their developmental timeframes and molecular requirements. These findings provide novel insights into the mechanisms of synaptic development and lay the foundation for future work determining their underlying etiologies.

## INTRODUCTION

Neuronal development is a complex, multi-step process that must occur over a specific timeframe and under precise conditions to produce a functioning nervous system. One crucial stage of this process is the formation of synapses (Wilson, 2013; Dalva et al., 2007; Lin and Goodman, 1994; Mosca and Luo, 2014; de Ramon Francàs et al., 2017; Siddiqui and Craig, 2011). Synaptic connections between neurons enable communication between cells as well as the processing of information to generate behavioral outputs (Farhy-Tselnicker and Allen, 2018). Despite having the same core function of synaptic communication, there is a vast diversity of neuron types in the nervous system (Masland, 2004; Wilton et al., 2019; Zeng and Sanes, 2017) and it is unclear if all neurons follow the same time-course of and rely on the same molecular pathways and mechanisms for completing synapse formation. Because different regions of the nervous system, such as the peripheral nervous system or the central nervous system (CNS), develop over different timeframes and under different conditions (Baines and Bate, 1998; Farhy-Tselnicker and Allen, 2018; Li et al., 2010; Liu and Chakkalakal, 2018), it is likely that variations in synaptic development exist based on neuron type. Historically, much of our understanding of synapse formation has come from the neuromuscular junction (NMJ); the NMJ in particular has been essential in providing early insight into receptor structure, the calcium hypothesis of neurotransmitter release, and spontaneous neurotransmission as well as the basic elements of synaptogenic signals like Agrin (Kummer et al., 2006; Li et al., 2018; Nitkin et al., 1987; Sanes and Lichtman, 1999; Shi et al., 2012; Wu et al., 2010). Many of the molecular mechanisms that regulate NMJ synapse formation are also found in the CNS, but a thorough understanding is currently lacking and it remains unknown whether peripheral synaptic mechanisms are fully conserved with those that govern central synapse development and organization (Collins and DiAntonio, 2007; Depew and Mosca, 2021; DePew et al., 2019; Goda and Davis, 2003; Keshishian et al., 1996). It is further unclear if all CNS neuron types follow the same rules for synapse formation given the considerable complexity of neuronal class, morphology, synaptic organization, and functional properties (including release probability, safety factor, firing frequency, etc) in the brain. Therefore, it is critical to determine the basic temporal and molecular characteristics of synapse development within central circuits and to understand how different potential mechanisms of development influence the diversity of synaptic organization in mature neurons. Knowing the general rules of normal development will inform our perspective on how defects during development can lead to neurodevelopmental and neurological disorders associated with synapse dysfunction such as autism, schizophrenia, or epilepsy (Bennett, 2011; Bonansco and Fuenzalida, 2016; Grant, 2012; Mullins et al., 2016). Further, there is considerable evidence that different neurological and neurodegenerative disorders, such as Parkinson’s Disease or Amyotrophic Lateral Sclerosis, specifically affect distinct populations of neurons over others (Dabool et al., 2019; Ilieva et al., 2009). The underlying basis for this selectivity remains unknown but understanding the functional, molecular, and developmental differences between different neuron types may provide unique insight into their vulnerability, or resistance, to such disease mechanisms.

The Drosophila antennal lobe (AL) represents a powerful model system for understanding the circuit-level, synaptic, and molecular aspects of how individual neuronal classes in a network undergo synaptic development (Grabe et al., 2016; Hummel and Rodrigues, 2008; Jefferis and Hummel, 2006; Mosca and Luo, 2014; Seki et al., 2017; Wilson, 2013). In Drosophila, olfaction is a guiding sensory modality, enabling flies to find food or mates, avoid danger, and communicate (Grabe and Sachse, 2018; Hildebrand and Shepherd, 1997). The AL is the first order olfactory structure in the fly brain (Couto et al., 2005; Fishilevich and Vosshall, 2005; Stocker et al., 1990) and is the essential gatekeeper of all olfactory information to higher brain centers. The AL is functionally analogous to the mammalian olfactory bulb (Sakano, 2020; Wilson, 2013) and represents a simple sensory circuit in which each neuron class is genetically accessible and current imaging technologies enable high resolution analysis with synaptic resolution (Li et al., 2019; Mosca and Luo, 2014; Wilson, 2013). Moreover, due to the high throughput nature of the system, different developmental stages are easily studied across multiple animals to examine stereotypy and variability during development. Taken together, current abilities enable a quantitative, temporal comparison of synapse development in AL neurons by visualizing the relative number of active zones, the sites of neurotransmitter release, in a single neuronal class and among different neuronal classes in the same sensory circuit. The multiple types of neurons found within the AL each have their own specific function, but also possess the shared goal of interpreting olfactory information to achieve a proper behavioral output. Olfactory receptor neurons (ORNs) comprise the primary sensory neuron population and receive input in the form of olfactory stimuli, detected from the outside environment, through the antennae and maxillary palps (Vosshall et al., 2000). Projection neurons (PNs) receive input from ORNs via dendritic terminals in the antennal lobe and project information to higher order olfactory centers like the mushroom body and the lateral horn via axon terminals (Modi et al., 2020) to result in a behavioral output (Jefferis et al., 2001). PNs also form dendrodendritic synapses with other PNs and synapse onto a third class of cells, the local interneurons (LNs). LNs form synapses with ORNs and other LNs throughout the ALs to regulate gain control of the olfactory signals, completing and refining the circuit (Tanaka et al., 2009; Yaksi and Wilson, 2010). The three main neuronal classes of the AL have clearly defined roles, but all interact to achieve the goal of odorant sensation. ORNs, PNs, and LNs thus comprise a complex, but tractable, sensory circuit suited to high resolution synaptic and developmental analyses (Figure 1A). We sought to compare the temporal and molecular programs ORNs, PNs, and LNs employ to accomplish synaptic development.

**Figure 1.**
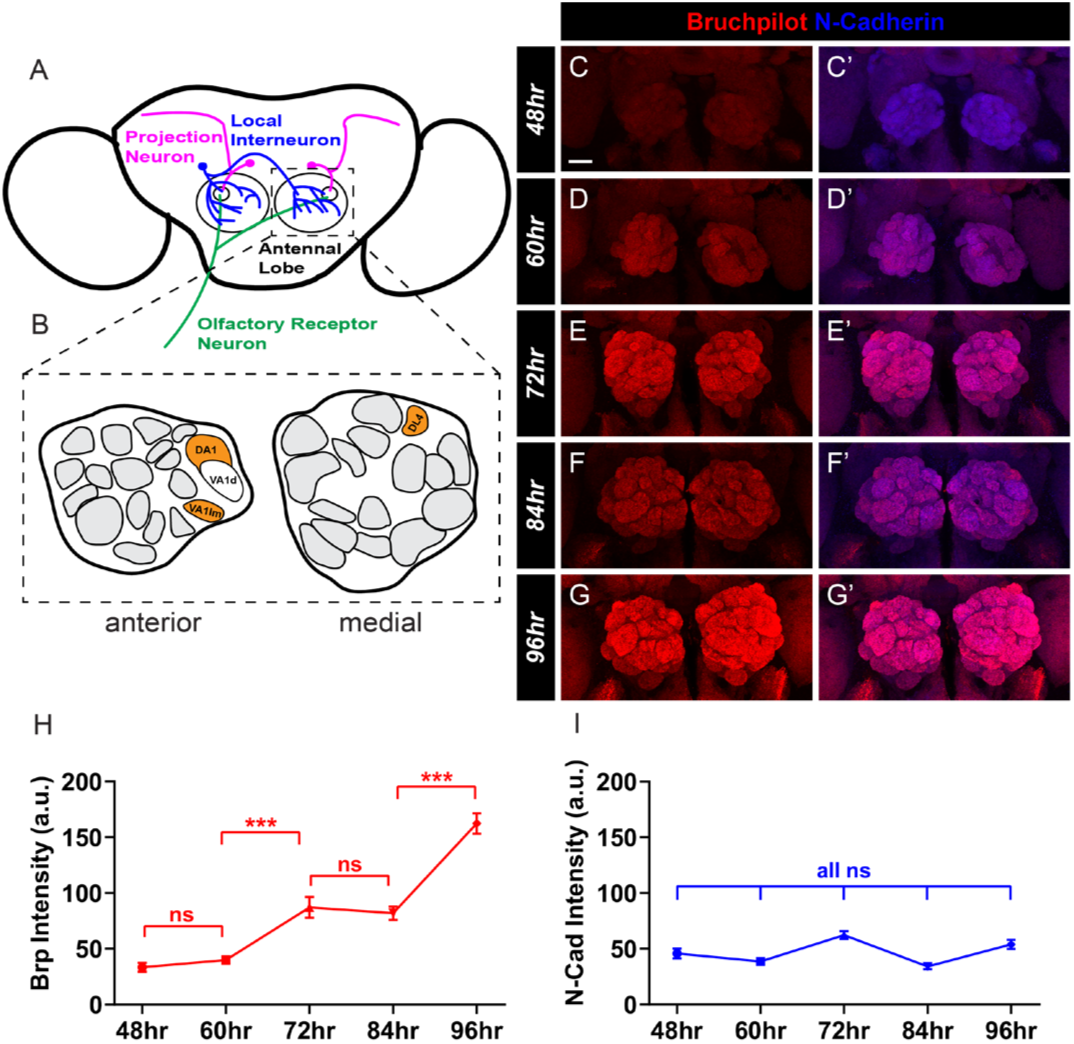
Synapse addition across the antennal lobe during pupal development occurs during distinct timeframes. A-B, Schematic of the Drosophila antennal lobes. Three major neuron classes project to the antennal lobes to relay and process olfactory information: Olfactory Receptor Neurons (green), Projection Neurons (magenta), and Local Interneurons (blue) (A). Each antennal lobe consists of about 50 glomeruli (B) with the three glomeruli examined throughout this study (DA1, VA1lm, and DL4) highlighted (orange). C-G’, Representative confocal image stacks of pupal antennal lobes stained with antibodies to Bruchpilot (red) and N-Cadherin (blue) at 48 (C), 60 (D), 72 (E), 84, (F) and 96 (G) hours after puparium formation. H-I, Quantification of average staining intensity for Brp (H) and N-Cadherin (I) from 48 to 96 hours APF. Brp intensity increases significantly from 60 to 72 hours APF as well as from 84 to 96 hours APF. N-Cadherin intensity does not change throughout this timeframe. For each time-point, n ≥ 9 antennal lobes from 8 brains. ***, p < 0.001. n.s. = not significant. Scale bar = 20 μm.

In this study, we mapped synaptic development in identified, genetically accessible classes of ORNs, PNs, and LNs across both sexes from early pupal development to mature adulthood in the Drosophila AL, creating the first, detailed, time-course of synaptic development in individual olfactory neuron populations. We found that ORNs, PNs, and LNs each have a distinct pattern of synaptic development, including periods of synaptic addition, removal or pruning stages, and synapse loss in late adulthood. We also determined that different subclasses of ORNs use different temporal programs of synaptic development based on their preferred odorant type, such as odors for food or pheromones, or sexual dimorphism. Having established distinct timelines of synapse addition and pruning that culminate in mature synaptic organization for each class of AL neurons, we examined how activity influenced mature synapse number after development in ORNs, PNs, and LNs. Intriguingly, beyond the temporal differences in synaptic development, we also determined that each class of AL neurons responded differently to perturbations in neuronal activity. When synaptic transmission was blocked cell-autonomously, ORNs, PNs, and LNs all showed a reduction in synapse number. Increasing electrical excitability, however, only impacted ORN synapse number, while PNs and LNs were unaffected. In the Drosophila olfactory system, neuronal activity can influence activity of GSK-3β, a master kinase, though it is unclear how it influences synapse formation (Chiang et al., 2009). We found that reducing GSK-3β kinase activity in ORNs impairs synapse addition, but similar GSK-3β impairments in PNs and LNs have no effect on synapse formation. Enhancing GSK-3β activity in both ORNs and LNs, however, phenocopied the effects of reduced neuronal excitability, suggesting that electrical activity may function through GSK-3β to regulate synapse formation. This suggests that though normal activity is required in all three AL neuron classes for mature synaptic development, each class may use different downstream molecular pathways under different circumstances to achieve proper development. These findings demonstrate that, even within the same sensory circuit, synaptic development is not a uniform process for all types of neurons and that the temporal and molecular programs that govern development can vary. Our results reveal novel insights into the mechanisms of synaptic development and demonstrate developmental individuality among different neuron classes.

## RESULTS

### Antennal lobe synapses are added over two distinct phases of pupal development

In the antennal lobe (AL), ORNs, PNs, and LNs (Figure 1A) comprise three major neuron types that contribute to the detection and subsequent transfer of olfactory information to higher order brain structures (Jefferis et al., 2001; Tanaka et al., 2009; Vosshall et al., 2000). Different classes of each neuron type project to the roughly 50 glomeruli that make up the AL, which are subdivided based on the type of olfactory information they receive (Figure 1B), and form synapses with each other to generate functional circuits (Axel et al., 2004; Grabe and Sachse, 2018; Hallem and Carlson, 2006; Jefferis et al., 2007). To begin to understand how synapse number changes over time in the antennal lobe at large, we measured the levels of Bruchpilot, an active zone protein and presynaptic marker (Fouquet et al., 2009; Wagh et al., 2006), during pupal development. During the roughly four-day metamorphic process of pupal development, larvae break down their cellular architecture, including their current nervous system, and re-form as an adult fly (Thummel, 2001). Neuronal wiring, axon pathfinding, and partner matching are largely complete by 48 hours after puparium formation (APF), indicating that synapses are unlikely to form before the component neurons of the AL have reached their final destination (Komiyama and Luo, 2006). We used antibody staining to measure Bruchpilot intensity (Laissue et al., 1999) from 48 hours APF to 96 hours APF (Figure 1C-G’). After 96h APF, pupation is complete and the adult fly ecloses. Quantification of Bruchpilot levels can serve as a measurement of the amount of active zone protein present – in cases of synapse development, the levels of Bruchpilot would necessarily increase concomitant with synapse addition as sites of neurotransmitter release grow more numerous. Changes in Bruchpilot are consistent with synapse formation and are regulated by genetic and molecular pathways that influence synaptic development (Fouquet et al., 2009; Mosca and Luo, 2014; Mosca et al., 2017; Wagh et al., 2006). We found that average Bruchpilot intensity in the AL increases significantly from 60-72 hours APF as well as from 84-96 hours APF (Figure 1H). Bruchpilot intensity remains largely unchanged, however, between 72 and 84 hours APF, suggesting that there are multiple phases of general synapse addition (Figure 1H). In contrast, the average intensity of N-cadherin, which functions a general neuropil marker in the fly brain (Hummel and Zipursky, 2004), did not change during the course of pupal development (Figure 1I). This increase in Bruchpilot levels during development suggests that the total number of synaptic connections within the entire antennal lobe circuit increases during specific timeframes.

### Pheromone-sensing olfactory receptor neurons develop synapses during pupation then remove synapses in late adulthood

Our results suggest that the multiple classes of antennal lobe neurons, when considered together, add synapses during specific time frames of pupal development (Figure 1). However, our data represents the aggregate of all neuronal classes and does not distinguish between each of the major classes of antennal lobe neurons (ORNs, PNs, and LNs). To understand how the cell-type specific synaptic development of each class of neurons progresses, we expressed Bruchpilot-Short (Brp-Short), an active zone label associated with presynaptic contacts (Mosca and Luo, 2014), in each class of neurons using the UAS-GAL4 binary expression system (Brand and Perrimon, 1993). We performed a high-resolution time course analysis of synaptic development throughout pupation and adulthood using confocal microscopy followed by three-dimensional rendering (see Methods). Brp-Short is a fluorescently tagged truncated variant of the active zone protein Bruchpilot (Fouquet et al., 2009; Mosca and Luo, 2014) that interacts with full-length Bruchpilot without interfering with its function or adding ectopic synapses (Fouquet et al., 2009; Mosca and Luo, 2014; Mosca et al., 2017; Wagh et al., 2006). Brp-Short signal manifests as synaptically localized puncta, acting as a proxy label for active zones, and accurately reports synapse number from specific cell populations and fold changes in synapse number due to genetic perturbation, as supported by genetic and electron microscopy studies (Berger-Müller et al., 2013; Christiansen et al., 2011; Coates et al., 2017, 2020; Fouquet et al., 2009; Kremer et al., 2010; Mosca and Luo, 2014; Mosca et al., 2017). We also labeled each neuronal class examined with membrane-bound GFP to fluorescently label neurites (Lee and Luo, 1999), enabling the quantification of neurite volume for each neuron class within a given glomerulus (Mosca and Luo, 2014; Mosca et al., 2017). To establish a time course of synaptic development, we quantified Brp-Short puncta and neuronal membrane volume marked by mCD8-GFP every 12 hours from 48 to 92 hours APF and then every 3-5 days, at regular intervals, from 0 to 21 days after eclosion in ORN classes that innervate distinct glomeruli. We specifically assayed pupal synapse development at 92 hours APF to ensure that all pupae were collected before eclosion (at 25°C, adult eclosion typically occurs between 92-96h APF).

ORNs are the primary sensory neurons responsible for conveying odorant information from the environment to the antennal lobes (Wilson, 2013). We first sought to assess the temporal synaptic development of ORNs that project to the VA1lm glomerulus (Figure 2A) and are genetically accessed by Or47b-GAL4 (Vosshall et al., 2000). This glomerulus receives pheromone odorant information (Fishilevich and Vosshall, 2005; Vosshall et al., 2000), is involved in oviposition as well as courtship decisions, influencing preference for younger mates (Sethi et al. 2019; Chin et al., 2018; Zhuang et al., 2016), and is sexually dimorphic, with male glomeruli being ∼40% larger than female glomeruli (Fishilevich and Vosshall, 2005; Mosca and Luo, 2014). We quantified Brp-Short puncta and neurite volume at 48, 60, 72, 84, and 92 hours APF as well as at 0, 5, 10, 15, and 20 days after eclosion (Figure 2B-K’’) in both male and females separately to assay any sexual dimorphisms in synaptic development. In male pupal ORNs, there were no synapses detectable at 48 hours APF, but synapses were then steadily added over pupal development with the number of Brp-Short puncta doubling from 60 to 92 hours APF (Figure 2L). Neurite volume was also negligible at 48 hours APF, possibly because the neurons must become more mature before mCD8-GFP can stably localize to the membrane, but then increased 3-fold from 60 hours APF to 92 hours APF (Figure 2M). This glomerulus-specific time-course of synapse formation differs from the time-course of gross AL synapse formation (Figure 1) in that synapses were added steadily throughout pupal development rather than in two separated time periods. After eclosion, there was a 36% increase in the number of Brp-Short puncta from 0 to 5 days of age followed by a 13% decrease in puncta number from 5 to 10 days of age (Figure 2N). Following 10 days of age, synapse number remained constant throughout the analysis. Adult neurite volume increased by 29% between 0 and 5 days after eclosion and then an additional 21% by 15 days after eclosion (Figure 2O). Together, these data suggest that synapses are steadily added from 48 hours APF to 5 days after eclosion, demonstrating that individual classes of olfactory neurons may show distinct synapse addition phases than the aggregate antennal lobe. This is followed by a pruning phase where synapses are removed prior to the completion of synaptic development and establishment of a steady state synapse number. Neurites continually grow and add volume until 15 days after eclosion, after which point, they achieve a steady state volume. We additionally determined the time-course of female VA1lm ORN synaptic development (Figure 2 – Figure Supplement 1A-K’’) to determine if any of the developmental mechanisms displayed sexual dimorphism or if they occurred similarly to male ORNs. There were fewer Brp-Short puncta and lower neurite volume at all time points in female VA1lm ORNs compared to male VA1lm ORNs, consistent with established sexual dimorphisms in size and synapse number (Stockinger et al. 2005; Mosca and Luo, 2014). We found that females also had a 2-fold increase in puncta and 3-fold increase in neurite volume from 60 to 92 hours APF (Figure 2P-Q). Adult female ORNs exhibited a similar increase in Brp-Short puncta to the male ORNs from 0 to 5 days of age that was followed by a decrease from 5 to 10 days after eclosion (Figure 2R). Adult neurite volume, however, significantly increased from 0 to 5 days after eclosion and then did not significantly change afterwards (Figure 2S), suggesting that female ORNs achieve their mature adult volume earlier than male ORNs. This suggests similarities in the general addition of synapses throughout development despite sexual dimorphisms in absolute synapse number but a minor difference in the growth rate of neurites during development between males and females. Overall, these time-courses demonstrate a general pattern of synaptic development for a sexually-dimorphic, pheromone-specific glomerulus in which synapses continually form until 5 days after eclosion and then decrease before stabilizing to a mature quantity.

**Figure 2.**
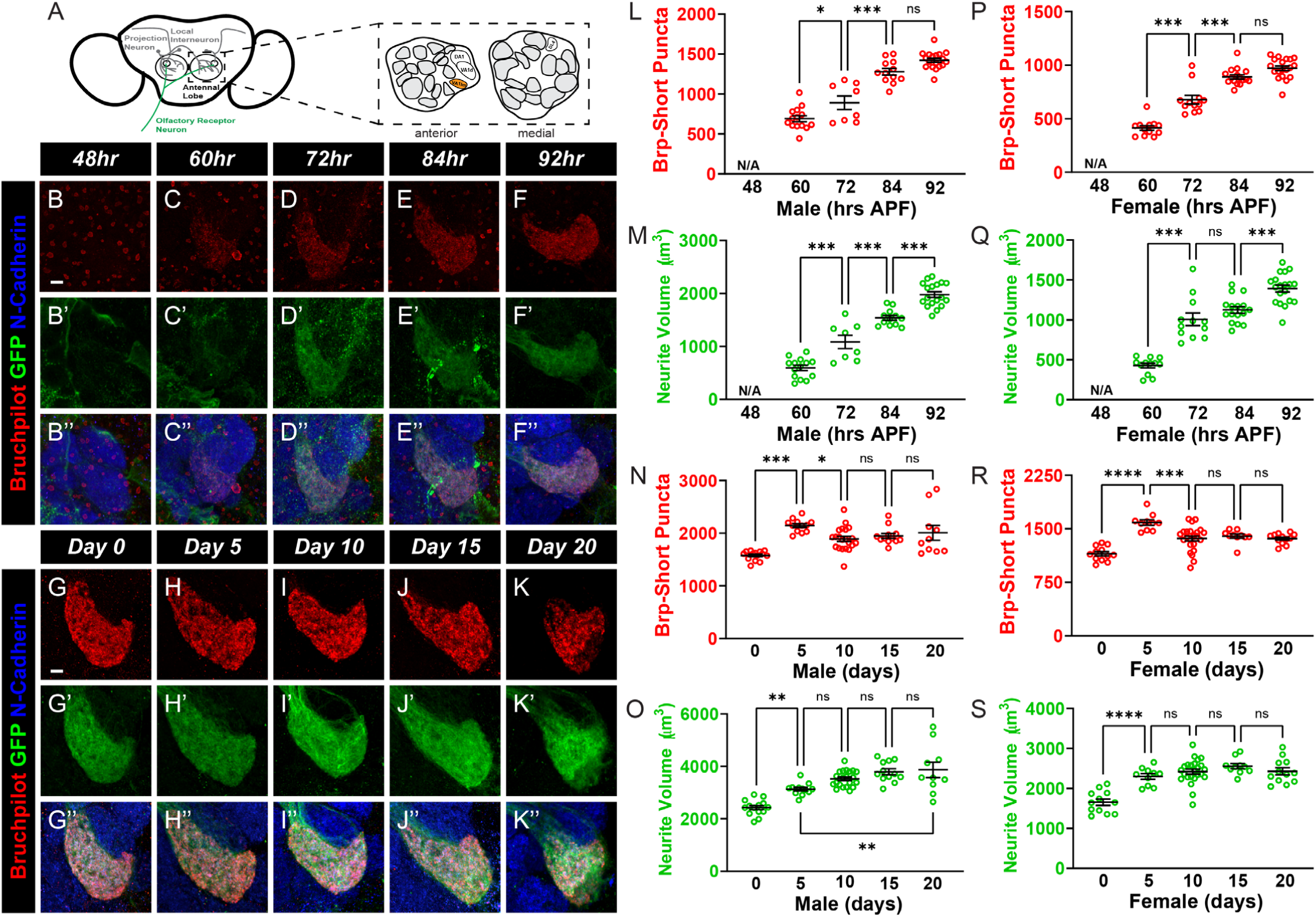
A developmental time course of synapse number and neurite volume in VA1lm ORNs. A. Schematic of the Drosophila antennal lobes showing ORNs (green) of the VA1lm glomerulus (orange). B-F’’, Representative confocal image stacks of male pupal VA1lm ORNs expressing Brp-Short-mStraw and membrane-tagged GFP and stained with antibodies against mStraw (red), GFP (green), and N-Cadherin (blue) at 48 (B), 60 (C), 72 (D), 84 (E), and 92 (F) hours APF. Note that synaptic puncta are not evident until 60 hours APF. G-K’’, Representative confocal image stacks of male adult VA1lm ORNs expressing Brp-Short-mStraw and membrane-tagged GFP and stained with antibodies as in B-F’’ at 0 (G), 5 (H), 10 (I), 15 (J), and 20 (K) days post eclosion. L-M, Quantification of Brp-Short-mStraw puncta (L) and membrane GFP volume (M) for pupal male ORNs. Both synaptic puncta and neurites are not measurable until 60h APF at which point they increase during the remainder of pupal development. N-O, Quantification of Brp-Short-mStraw puncta (N) and membrane GFP (O) volume for adult male ORNs. Synaptic puncta significantly increase from 0 to 5 days old and then decrease between 5 and 10 days post eclosion. Neurite volume increases from 0 to 15 days of age before achieving a steady state. P-Q, Quantification of Brp-Short-mStraw puncta (P) and membrane GFP volume (Q) for pupal female ORNs. Both synaptic puncta and neurites are not measurable until 60h APF at which point they increase during the remainder of pupal development. R-S, Quantification of Brp-Short-mStraw puncta (R) and membrane GFP (S) volume for adult female ORNs. Synaptic puncta significantly increase from 0 to 5 days old and then decrease between 5 and 10 days post eclosion. Neurite volume increases from 0 to 5 days of age before achieving a steady state. For each time-point, n ≥ 8 glomeruli from 4 brains. *, p < 0.05, **, p < 0.01, ***, p < 0.001, n.s. = not significant. Scale bar = 5 μm.

We next sought to determine if another sexually dimorphic class of ORNs, the Or67d-positive ORNs that project to the DA1 glomerulus (Figure 3A), develop similarly to VA1lm ORNs or if there are differences based on glomerulus (Datta et al., 2008). DA1 ORNs detect male pheromones, including 11-cis-Vaccenyl acetate, that are involved in courtship and aggression for both male and female flies (Datta et al., 2008; Kurtovic et al., 2007; Wilson, 2013). Similar to VA1lm, DA1 is a sexually-dimorphic glomerulus with increased glomerular size in males compared to females (Mosca and Luo, 2014; Stockinger et al., 2005). We used a GAL4 driver that labels DA1 ORNs via GAL4 expression under the control of the endogenous Or67d promoter (Stockinger et al., 2005) to examine the time course of synapse formation during adult stages; Or67d-GAL4 is not active during pupal development precluding our analysis of synapse development in these ORNs from 48-92h APF. We quantified Brp-Short puncta number and neurite volume for male DA1 ORNs every 3 days from 0-18 days of age (Figure 3B-H’’) and found that Brp-Short puncta increased by 50% between 0 and 6 days of age before achieving and maintaining a steady state synapse number until day 15. Surprisingly, we observed a 25% decline in synapse number between days 15 and 18, (Figure 3I). Neurite volume doubled from 0 to 6 days old and then did not significantly change for the remainder of adulthood (Figure 3J), thus maintaining a steady state. We found congruent results with female DA1 ORNs (Figure 3 – Figure Supplement 1A-H’’). Brp-Short puncta number in female ORNs significantly increased by 20% (Figure 3K) from 0 to 3 days of age, maintained a steady state from 3-12 days, and then decreased by 15% from 12 to 18 days of age. Neurite volume significantly increased by 33% from 0 to 3 days of age and then did not significantly change for the rest of adulthood (Figure 3L). Thus, the general female time-course of DA1 ORNs was nearly identical to the male time-course, with slight variability in the exact times of synapse change. Compared to VA1lm ORNs, DA1 ORN synapse development showed similar synapse addition until 5-6 days after eclosion but did not display a synapse reduction phase until 12-18 days old while VA1lm ORNs displayed synapse pruning from 5-10 days old. These data imply that pheromone-specific ORNs follow similar general patterns of synaptic development but have critical differences in the exact timing of developmental phases.

**Figure 3.**
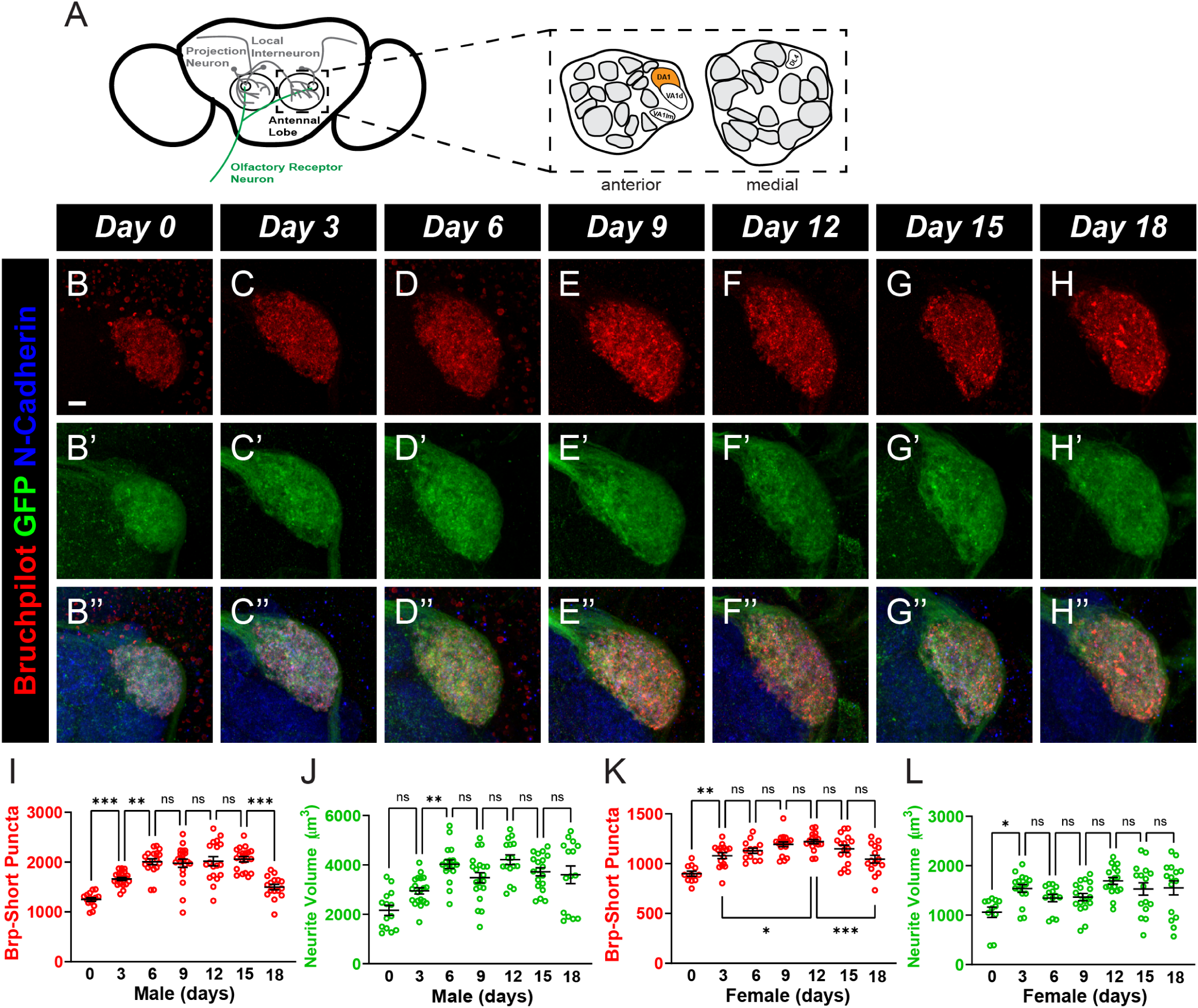
A developmental time course of synapse number and neurite volume in adult DA1 ORNs. A. Schematic of the antennal lobes showing ORNs (green) of the DA1 glomerulus (orange). B-H’’, Representative confocal image stacks of male adult DA1 ORNs expressing Brp-Short-mStraw and membrane-tagged GFP and stained with antibodies against mStraw (red), GFP (green), and N-Cadherin (blue) at 0 (B), 3 (C), 6 (D), 9 (E), 12 (F), 15 (G), and 18 (H) days of age. I-J, Quantification of Brp-Short-mStraw puncta (I) and membrane GFP volume (J) for adult male DA1 ORNs. Both Brp-Short puncta number and neurite volume steadily increase from 0 to 6 days of age; after 6 days post eclosion, neurite volume stabilizes while Brp-Short puncta maintain a steady state before decreasing between 15 and 18 days of age. K-L, Quantification of Brp-Short-mStraw puncta (K) and membrane GFP volume (L) for adult female DA1 ORNs. Both Brp-Short puncta number and neurite volume steadily increase from 0 to 3 days of age, after which neurite volume stabilizes. Brp-Short puncta maintain a steady state before decreasing between 12 and 18 days of age. For each time-point, n ≥ 11 from 6 brains glomeruli. **, p < 0.01, ***, p < 0.001, n.s. = not significant. Scale bar = 5 μm.

### Food-sensing olfactory receptors neurons develop synapses during pupation and early adulthood then stabilize

Drosophila rely on olfaction to find and interact with mates as well as identify suitable food sources. Food sensing glomeruli lack that sexual dimorphism seen in size and synapse number of pheromone-sensing glomeruli (Couto et al., 2005; Mosca and Luo, 2014; Stockinger et al., 2005), likely because all flies have a shared, unabating need to identify food sources. Our assays on sexually dimorphic glomeruli thus far followed the synaptic development of pheromone-sensing ORNs but we further sought to understand if the same developmental features apply in glomeruli that sense food odorants or if these glomeruli followed a distinct developmental program. We first established a time-course for male ORNs that project to the DL4 glomerulus (Figure 3 – Figure Supplement 2A), a class of neurons that conveys information about volatile chemical odorants related to food (Endo et al., 2007; Martin et al., 2013; Mosca and Luo, 2014), throughout pupal (48 to 92 hours APF) development (Figure 3 – Figure Supplement 2B-F’’) and into adulthood from 0 to 18 days old (Figure 3 – Figure Supplement 2G-M’’). We found that pupal Brp-Short puncta increased 1.5-fold from 48 to 72 hours APF and then did not significantly change (Figure 3 – Figure Supplement 2N) throughout the rest of pupation. Pupal neurite volume also exhibited a nearly 1.5-fold change from 48 to 72 hours APF before reaching a steady-state neurite volume that was maintained until adulthood (Figure 3 – Figure Supplement 2O), suggesting much of the development of pupal DL4 ORNs occurs well in advance of eclosion. Adult male Brp-Short puncta in DL4 ORNs significantly increased by 1.2-fold from 0 to 3 days after eclosion before achieving their mature synapse number and maintaining that number of puncta throughout the remainder of observed adulthood (Figure 3 – Figure Supplement 2P), further demonstrating that synapse formation continues into adulthood. Adult neurite volume did not significantly change from 0 to 6 days after eclosion, maintaining a similar level to that measure during pupal stages – intriguingly, however, neurite volume declined from 6-15 days by 39% (Figure 3 – Figure Supplement 2Q). We additionally generated a time-course of development for female DL4 ORNs (Figure 3 – Figure Supplement 3A-M’’) to examine any potential sexual dimorphism. Female DL4 ORNs showed a 1.5-fold increase in both Brp-Short puncta (Figure 3 – Figure Supplement 2R) and neurite volume (Figure 3 – Figure Supplement 2S) from 48 to 60 hours APF with no significant changes for the remainder of pupal development. Adult female DL4 Brp-Short puncta significantly decreased between 6 and 12 days after eclosion but did not significantly change between any other time-points (Figure 3 – Figure Supplement 2T). Female DL4 ORN adult neurite volume also did not significantly change from 0 to 6 days of age (Figure 3 – Figure Supplement 2U) but significantly declined between 6 and 9 days (Figure 3 – Figure Supplement 2U). While neurite volume steadily decreased as the fly aged in both sexes, this is likely an effect of the AM29-GAL4 driver weakening over time (Endo et al., 2007). Overall, however, the absolute puncta numbers and neurite volumes at each stage were comparable between males and females, implying that synaptic development for food sensing ORNs is not sexually dimorphic and that both sexes use similar developmental programs for food-sensing ORNs. Thus, food-sensing ORNs utilize a different time-course of synaptic development than pheromone-sensing glomeruli like DA1 and VA1lm – one that emphasizes early development and lacks sexual dimorphism. Taken together, this indicates that ORNs of sexually dimorphic glomeruli follow similar rules for synaptic development which are distinct from the rules followed by non-sexually dimorphic ORNs and that within classes, each ORN subtype has subtle, individual variations in development.

### Projection Neurons continually develop synapses during pupation and early adulthood

Projection neurons (PNs) comprise the second order of neurons of the olfactory system, relaying odorant information from ORNs that project onto PN dendrites in the antennal lobe to higher order brain structures such as the mushroom bodies and the lateral horn (Modi et al., 2020). ORN to PN communication is vital for making behavioral decisions, including foraging, oviposition, and courtship, based on the odor information available to the fly (Jefferis et al., 2001). Despite the importance of these neurons, we lack a detailed understanding of how they develop synapses over time. We addressed this by constructing a time-course of synapse formation for male DA1 PNs (Figure 4A) using genetic access via the Mz19-GAL4 driver (Jefferis et al., 2004); Mz19-GAL4 drives expression in DA1, VA1d, and DC3 PNs, but we restricted our analysis to DA1 PNs to enable a direct comparison to the ORNs (and LNs, see below) of the same glomerulus (Figure 3, Figure 3 – Figure Supplement 1). This allowed us to assess if different classes of neurons belonging to the same microcircuit, such as those observed within an olfactory glomerulus, would share developmental programs or utilize distinct modes of synapse addition. As with ORNs, we quantified Brp-Short puncta number and neurite volume from 48 to 92 hours APF and from 0 to 18 days after eclosion (Figure 4B-M’’). Unlike ORNs, however, Brp-Short puncta were clearly visible in PNs at 48 hours APF and steadily increased in number over 2.5-fold by 92 hours APF (Figure 4N). Pupal neurite volume also continually increased during pupation, doubling from 48 hours to 92 hours APF (Figure 4O). This data show that PN synapses develop throughout the course of pupation. During adulthood, Brp-Short puncta continued to be added after eclosion with puncta number increasing 2-fold from 0 to 6 days of age, followed by no significant changes for the rest of adulthood (Figure 4P). Neurite volume increased nearly 2-fold from 0 to 3 days after eclosion and then remained unchanged throughout the rest of adulthood (Figure 4Q). Therefore, DA1 PNs develop synapses throughout pupation and even early into adulthood before stabilizing at their mature quantity. Unlike ORNs, however, they lack a pruning phase and reach their mature state more quickly.

**Figure 4.**
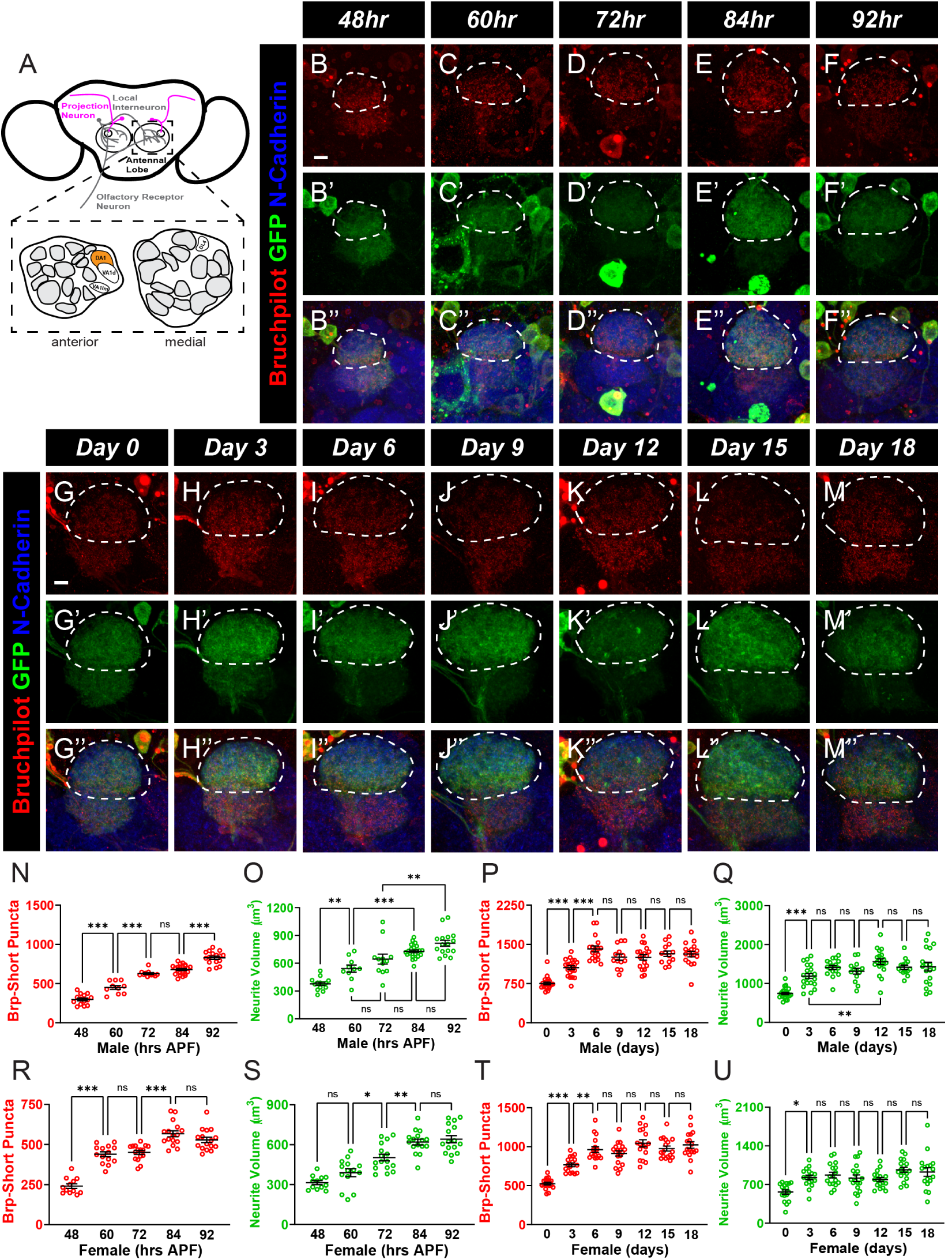
A developmental time course of synapse number and neurite volume in DA1 projection neurons. A. Schematic of the antenna lobes showing PNs (magenta) of the DA1 glomerulus (orange). B-F’’, Representative confocal image stacks of male pupal DA1 PNs (dashed white lines) expressing Brp-Short-mStraw and membrane-tagged GFP and stained with antibodies against mStraw (red), GFP (green), and N-Cadherin (blue) at 48 (B), 60 (C), 72 (D), 84 (E), and 92 (F) hours APF. G-M’’, Representative confocal image stacks of male adult DA1 PNs (dashed white lines) expressing Brp-Short-mStraw and membrane-tagged GFP and stained with antibodies as in A-E at 0 (G), 3 (H), 6 (I), 9 (J), 12 (K), 15 (L), and 18 (M) days of age. N-O, Quantification of Brp-Short-mStraw puncta (N) and membrane GFP volume (O) for pupal male DA1 PNs. Brp-Short puncta and accompanying neurites are visible at 48h APF and increase steadily during pupal development. P-Q, Quantification of synaptic puncta (P) and neurite volume (Q) for adult male DA1 PNs. Brp-Short puncta and neurite volume increase until 6d after eclosion after which both stabilize for the remainder of adulthood. R-S, Quantification of Brp-Short-mStraw puncta (R) and membrane GFP volume (S) for female pupal DA1 PNs. Synaptic puncta are added between 48 and 60 hours APF as well as between 72 and 84 hours APF. Neurite volume steadily increases until about 84-92 hours APF. T-U, Quantification of synaptic puncta (T) and neurite volume (U) for female adult DA1 PNs. Synapses addition occurs from 0 to 6 days of age and then stabilizes. Neurite volume slightly increases from 0 to 3 days and the remains constant. For each time-point, n ≥ 10 glomeruli from 5 brains. *, p < 0.1, **, p < 0.01, ***, p < 0.001, n.s. = not significant. Scale bar = 5 μm.

We additionally generated a time-course of development for female DA1 PNs to examine any potential sexual dimorphisms in PN synaptic development (Figure 4 – Figure Supplement 1A-M’’). In female PNs, both the number of Brp-Short puncta (Figure 4R) and neurite volume increased 2-fold from 48 to 92 hours APF (Figure 4S), showing that males and females both develop synapses during the entirety of pupation. Adult female PNs exhibited a nearly 2-fold increase in Brp-Short puncta from 0 to 6 days of age, and then did not change for the remainder of adulthood (Figure 4T). Female neurite volume increased 1.5-fold from 0 to 3 days after eclosion and then did not significantly change (Figure 4U, similar to males). These data demonstrate that male and female PNs have nearly identical time-courses of synaptic development. Furthermore, their exact timeframes of synapse addition and stabilization are distinct from those of ORNs. Therefore, PNs that receive information predominantly from pheromone-sensing ORNs have a unique pattern of synapse formation, demonstrating that different neuronal classes within the same microcircuit have distinct patterns of synaptic development. This may underlie their specific function in the antennal lobe olfactory circuit (see Discussion).

### Local interneurons rapidly undergo synapse formation late in pupation to reach maturity

In the antennal lobe, the most diverse class of cells are local interneurons (Chou et al., 2010; Ng et al., 2002; Tanaka et al., 2008, 2012). Compared to ORNs and PNs, less is known about their precise physiological functions. LNs have been implicated in a number of processing steps in olfactory information including gain control, regulating interglomerular communication, and refining olfactory signal (Chou et al., 2010; Hong and Wilson, 2015; Tanaka et al., 2009; Yaksi and Wilson, 2010). Within the antennal lobe, different types of LNs synapse onto both ORN and PN targets. The multiglomerular LNs comprise the most numerous class and innervate multiple glomeruli throughout the antennal lobe. However, little is known about how synapse addition and active zone development occurs in these essential cells. Given their prominence among the olfactory LNs, we wanted to understand how their synapses developed. Therefore, we established a time course of synapse addition and neurite growth during development for the multiglomerular LNs accessed genetically by NP3056-GAL4 (Chou et al., 2010) for males (Figure 5A) as well as females (Figure 5 – Figure Supplement 1A). To enable a direct comparison with Or67d-expressing ORNs and Mz19-accessed PNs, we focused our examination specifically to the projections of those multiglomerular LNs within the DA1 glomerulus (Figure 5B-M’’). This allowed us to compare synapse formation between each neuron class for the same precise region of the antennal lobe, eliminating cellular geography as a potential variable influencing synaptic development.

**Figure 5.**
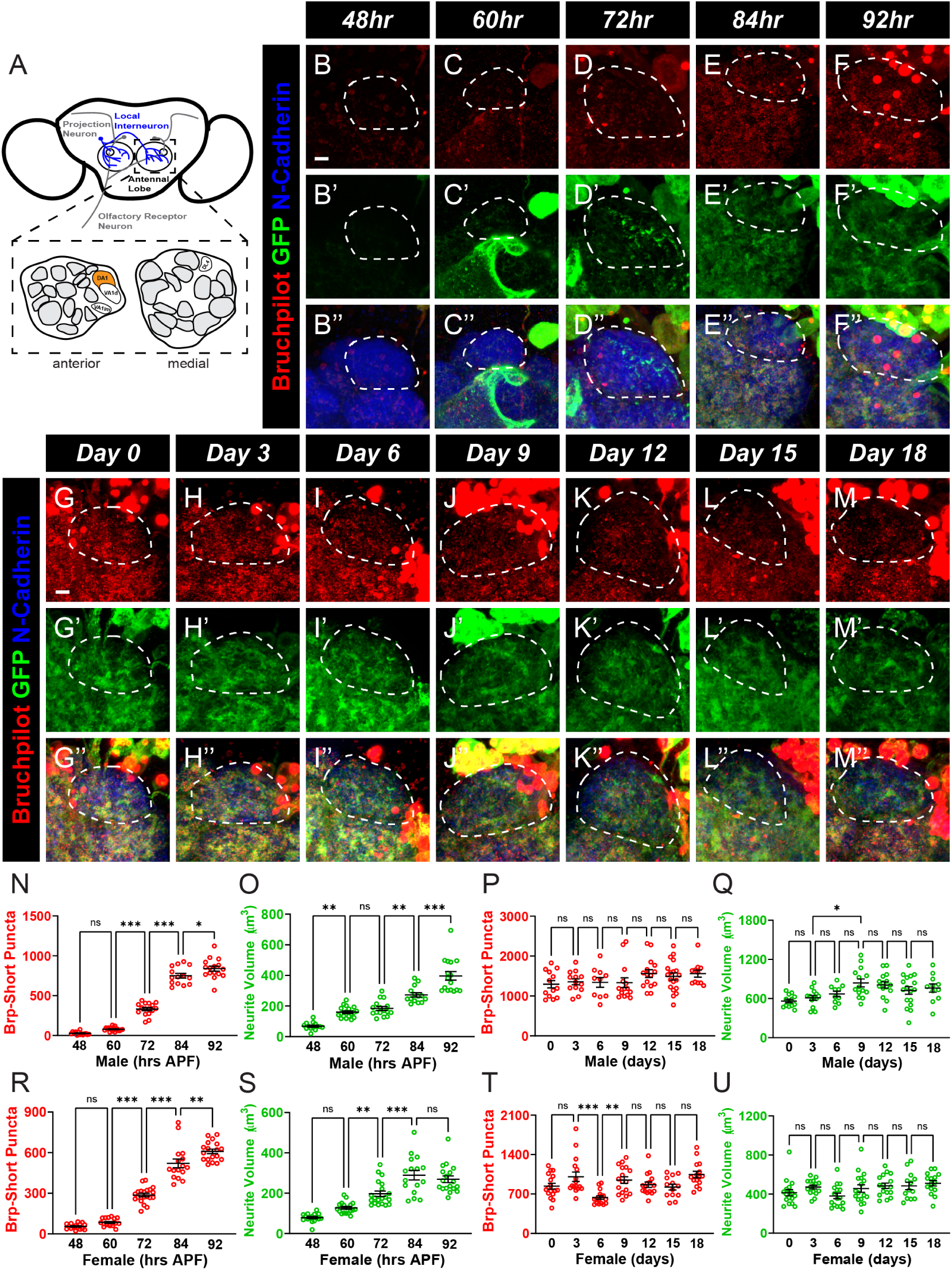
A developmental time course of synapse number and neurite volume in DA1 local interneurons. A. Schematic of the antennal lobes showing LNs (blue) of the DA1 glomerulus (orange). B-F’’, Representative confocal image stacks of male pupal DA1 LNs (dashed white lines) expressing Brp-Short-mStraw and membrane-tagged GFP and stained with antibodies against mStraw (red), GFP (green), and N-Cadherin (blue) at 48 (B), 60 (C), 72 (D), 84 (E), and 92 (F) hours APF. G-M’’, Representative confocal image stacks of male adult DA1 LNs (dashed white lines) expressing Brp-Short-mStraw and membrane-tagged GFP and stained with antibodies as in B-F’’ at 0 (G), 3 (H), 6 (I), 9 (J), 12 (K), 15 (L), and 18 (M) days of age. N-O, Quantification of Brp-Short-mStraw puncta (N) and membrane GFP volume (O) for pupal male DA1 LNs. Synaptic puncta and neurites begin to significantly form between 60 and 72 hours APF and rapidly increase for the remainder of pupal development. P-Q, Quantification of synaptic puncta (P) and neurite volume (Q) for adult male DA1 LNs. There is no significant difference in the number of synaptic puncta or neurite volume across all adult times examined while neurite volume increases until 9 days of age. R-S, Quantification of Brp-Short-mStraw puncta (R) and membrane GFP volume (S) for female pupal DA1 LNs. Addition of quantifiable synaptic puncta and neurite volume begins between 48 and 60 hours APF and then occurs rapidly until 84-92 hours APF. T-U, Quantification of synaptic puncta (T) and neurite volume (U) for female adult DA1 LNs. Synapses remain largely constant throughout adulthood, except for a decrease and subsequent increase from 3 to 9 days of age. Neurite volume is constant throughout all of adulthood. For each time-point, n ≥ 10 glomeruli from 6 brains. *, p < 0.05, **, p < 0.01, ***, p < 0.001, n.s. = not significant. Scale bar = 5 μm.

From 48 to 60 hours APF, we observed very few LN synapses in DA1 and neurite elaboration within the glomerulus was minimal (Figure 5B-C’’). Between 60- and 92-hour APF, however, there was significant addition of active zones and neurite growth (Figure 5D-F’’). In the final 30 hours of pupation, active zone number increased nearly 10-fold (Figure 5N). LN neurite volume within DA1 also increased nearly 2.5-fold during the same time period (Figure 5O). We also determined the time course of synapse addition and neurite growth for multiglomerular LNs during the first 18 days of adulthood. At day 0, immediately after eclosion, we observed a higher number of active zones than at 92h APF, indicating that even during the short time between 92h APF and eclosion, there was a significant amount of synapse addition. The same was true of neurite outgrowth. Surprisingly, however, we found that synapse addition remained constant from 0-to 18-day old adult flies (Figure 5P). Neurite outgrowth increased nearly 2-fold from 0 to 9 days after eclosion and then remained stable (Figure 5Q). Female DA1 LNs appeared nearly identical to male LNs (Figure 5 – Figure Supplement 1B-M’’). From 60h to 92h APF, female Brp-Short puncta increased 7-fold (Figure 5R). In the same timeframe, neurite volume increased 2-fold (Figure 5S). Female adult LNs differed from males in that Brp-Short puncta decreased 37% from 3 days of age to 6 days of age and then re-increased 49% at 9 days of age (Figure 5T). Neurite volume, however, did not change after eclosion (Figure 5U). Overall, this suggests that LNs complete most synapse addition during pupal development. Males and females both showed the same general pattern of synapse formation, with the expected sex-specific dichotomy in synapse number and neurite volume (as observed previously in ORNs and PNs – Stockinger et al., 2005 and Figure 3-4). Female LNs, however, showed more variation throughout the adult time course than males, who retained their steady state of synapse number once achieved. This unique timeline of development for multiglomerular LNs in the DA1 glomerulus highlights an additional developmental program that differs from the programs previously observed in ORNs and PNs

### Decreasing neuronal activity lowers synapse number in antennal lobe neurons

In a glomerular microcircuit, ORNs, PNs, and LNs have interconnected projections and fulfill distinct individual roles to support the larger goal of the antennal lobe in processing olfactory information. Our data suggests that each class of antennal lobe neurons, despite a shared goal and common microcircuit, utilizes different temporal programs to complete synaptic and neurite development. As even closely related neurons of the same microcircuit use different developmental programs, this raised the possibility that each class of neurons may further utilize different molecular and cellular mechanisms to accomplish synaptic formation. One well-characterized modulator of synapse formation and plasticity is neuronal activity (Cang and Feldheim, 2013; Simi and Studer, 2018). Neurons with increased activity can outcompete other, less active neurons for synaptic partners as well as increase the number of synaptic connections made to other neurons (Miller, 1996). In Drosophila, there is a rich history of studying activity-dependent synaptic development at peripheral neuromuscular synapses (Keshishian et al. 1996); there, changes in activity influence synaptic development in terms of bouton number, ectopic projection, active zone organization, and physiological output (Akin and Zipursky, 2020; Harris and Littleton, 2015; Hazan and Ziv, 2020; Jarecki and Keshishian, 1995; Schmid et al., 2008). The role of activity-dependent synapse organization in the CNS is less well understood, raising questions as to whether all synapses follow similar activity-dependent requirements for development or, as our temporal data might suggest, there is also heterogeneity within molecular and cellular programs that different central neurons use to complete development. We first sought to determine if antennal lobe neuron synapse development in ORNs, PNs, or LNs could be regulated by activity-dependent mechanisms. To accomplish this, we either decreased or increased neuronal activity chronically throughout development in ORNs, PNs, or LNs and quantified Brp-Short puncta number and neurite volume in those neurons. Because our developmental analyses showed that ORNs, PNs, and LNs all achieved stable synapse number by 10 days post eclosion (Figures 2-5), we specifically quantified synapse organization in 10-day old adult flies. This allowed us to observe how changes in neuronal activity during development influence mature synapse density and to compare our findings between each neuron class. To decrease neuronal activity, we expressed an active tetanus toxin (TeTxLc) in either ORNs, PNs, and LNs that innervate the DA1 glomerulus and compared Brp-Short puncta numbers to neurons that expressed an inactive variant of the toxin as a control (Sweeney et al., 1995). Tetanus toxin lowers neuronal activity by cleaving Synaptobrevin at the synapse, impairing synaptic vesicle fusion and blocking neurotransmission (Sweeney et al., 1995). For male and female ORNs, there was a 29% and 21% decrease, respectively, in Brp-Short puncta number in neurons where active TeTxLc was expressed compared to the inactive control (Figure 6A-B’, E-F; Figure 6 – Figure Supplement 1A-B’). However, neurite volume was unaffected by reduced activity (Figure 6C-D’, E-F; Figure 6 – Figure Supplement 1C-D’). PNs also exhibited a significant decrease (by 60% for males and 36% for females) in Brp-Short puncta number following blocked synaptic transmission (Figure 6G-H’, K-L; Figure 6 – Figure Supplement 1E-F’). Unlike ORNs, however, neurite volume was also significantly decreased in PNs with decreased neuronal activity (Figure 6I-J’, K-L; Figure 6 – Figure Supplement 1G-H). Finally, we observed that TeTxLc expression in LNs caused a reduction in Brp-Short puncta (43% in males, 28% in females; Figure 6M-N’, Q-R; Figure 6 – Figure Supplement 1I-J’) but did not influence neurite volume (Figure 6O-P’, Q-R; Figure 6 – Figure Supplement 1K-L’). These results indicate that synapse addition is dependent on a baseline level of neuron activity across all antennal lobe neuron classes, but that neurite growth is activity-dependent in PNs and activity-independent in ORN and LN classes. This suggests that different activity-related mechanisms exist to control specific aspects of synapse organization in different classes of cells.

**Figure 6.**
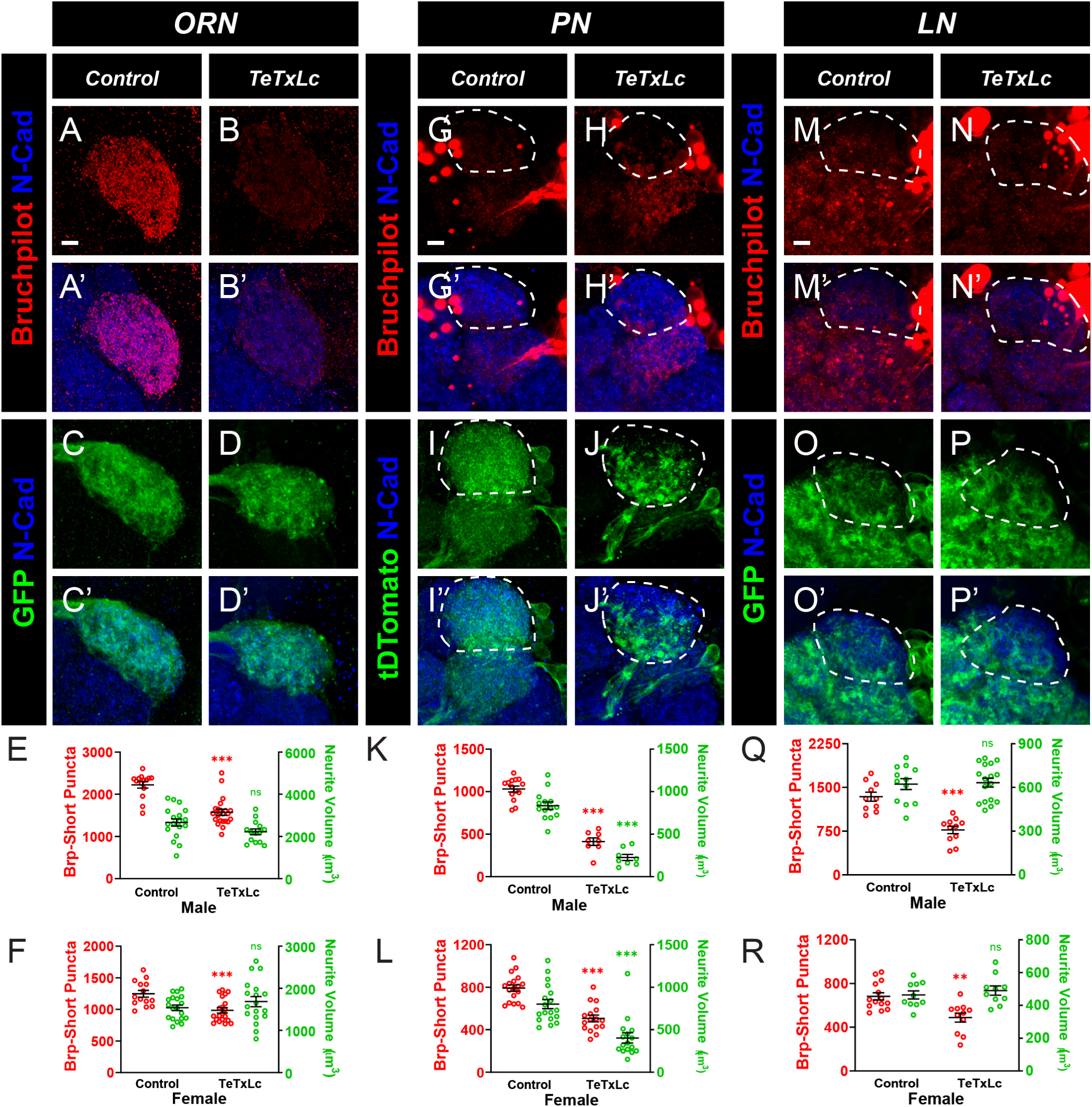
Normal neuronal activity is required for normal synaptic development in ORNs, PNs, and LNs of the antennal lobe. A-D’, Representative confocal image stacks of ORNs of the DA1 glomerulus expressing either Brp-Short-mStraw (A-B) or a membrane-tagged GFP (C-D) and either an inactive (A, C) or active (B, D) variant of tetanus toxin light-chain (TeTxLc). Brains were stained with antibodies against mStraw (red) or GFP (green) and N-Cadherin (blue). E, Quantification of Brp-Short puncta number and neurite volume between male DA1 ORNs expressing either active or inactive tetanus toxin. Expression of active tetanus toxin reduces the number of synaptic puncta without altering neurite volume. F, Quantification of Brp-Short puncta number and neurite volume between female DA1 ORNs expressing either active or inactive tetanus toxin. Expression of active tetanus toxin reduces the number of synaptic puncta without altering neurite volume. G-J’, Representative image stacks of 10-day old DA1 PNs (dashed white lines) expressing Brp-Short-GFP (G-H) or membrane-tagged tdTomato (I-J) and either inactive (G, I) or active (H, J) tetanus toxin. Brains were stained with antibodies against tdTomato (green) or GFP (red) and N-Cadherin (blue). K, Quantification of Brp-Short puncta number and neurite volume between male DA1 PNs expressing tetanus toxin dead or active. Decreasing neuronal activity in PNs results in fewer synaptic puncta and reduced neurite volume. L, Quantification of Brp-Short puncta number and neurite volume in female DA1 PNs expressing tetanus toxin dead or active. Decreasing neuronal activity in PNs results in fewer synaptic puncta and reduced neurite volume. M-P’, Representative image stacks of 10-day old DA1 LNs (dashed white lines) expressing Brp-Short-mStraw (M-N) or membrane-tagged GFP (O-P) as well as inactive (M, O) or active (N, P) tetanus toxin. Brains were stained with antibodies as in A-D’. Q, Quantification of Brp-Short puncta number and neurite volume between male DA1 LNs expressing tetanus toxin dead or active. Decreasing synaptic transmission in LNs decreased synaptic puncta but did not affect neurite volume. R, Quantification of Brp-Short puncta number and neurite volume between female DA1 LNs expressing tetanus toxin dead or active. Decreasing synaptic transmission in LNs decreased synaptic puncta but did not affect neurite volume. For each experimental group, n ≥ 8 glomeruli from 4 brains. **, p < 0.01, ***, p < 0.001, n.s. = not significant. Scale bar = 5 μm.

### Increasing neuronal activity increases ORN, but not PN or LN synapse formation

Neuronal activity regulates synapse formation and the strength of connections within neural circuits (Cang and Feldheim, 2013; Pan and Monje, 2020; Simi and Studer, 2018; Vonhoff and Keshishian, 2017). We established that reduced neural activity results in fewer synapses in all classes of neurons observed and sought next to determine if the reciprocal relationship was true for antennal lobe neurons: does increased electrical activity result in additional synapses? To increase neuronal activity in DA1 ORNs, PNs, and LNs, we expressed the bacterial sodium channel NaChBac, which increases electrical activity by enhancing the inward Na+ current into the cells, resulting in additional action potentials and neuronal firing (Nitabach et al., 2006; Ren et al., 2001). We observed that NaChBac expression in male or female DA1 ORNs (Figure 7A-D’; Figure 7 – Figure Supplement 1A-D’) led to a 49% or 38% increase, respectively, in Brp-Short puncta (Figure 7E-F) but did not significantly alter neurite volume (Figure 7E-F). This suggests a direct relationship between cell-autonomous neuronal activity and ORN synapse formation. Intriguingly, we found that this relationship did not exist in PNs: NaChBac expression in DA1 PNs had no effect on either Brp-Short puncta number (Figure 7G-H’, K-L; Figure 7 – Figure Supplement 1E-F’) or neurite volume (Figure 7I-J’, K-L; Figure 7 – Figure Supplement 1G-H’) in males or females. This suggests that PN synapse formation is resistant to increases in electrical activity, but sensitive to reduced neuronal firing, indicating partial activity-dependence. Finally, NaChBac expression in LNs exhibited sexually dimorphic changes in synapse number. Increased LN activity in males had no effect on Brp-Short puncta number (Figure 7M-N’, Q) or neurite volume (Figure 7O-P’, Q) but significantly reduced Brp-Short puncta (Figure 7R; Figure 7 – Figure Supplement 1I-J’) in females by 19% without altering neurite volume (Figure 7R; Figure 7 – Figure Supplement 1K-L’). This indicates a number of aspects about the relationship between LN synapse formation and neural activity – 1) that there is a sexual dimorphism between males and females in the activity-dependence of synapse formation; 2) in females, there is likely to be an optimal electrical activity that promotes synapse formation whereby any alteration (increase or decrease) results in impaired synapse formation; and 3) in males, there is likely to be some level of independence from increased activity in regulating synapse formation but sensitivity to reduced activity in regulating synapse formation (similar to PNs but in contrast to ORNs). Collectively, these results demonstrate that each class of neuron is influenced differently by increased neuronal activity and that the rules that govern synapse formation are not the same for each class of neuron.

**Figure 7.**
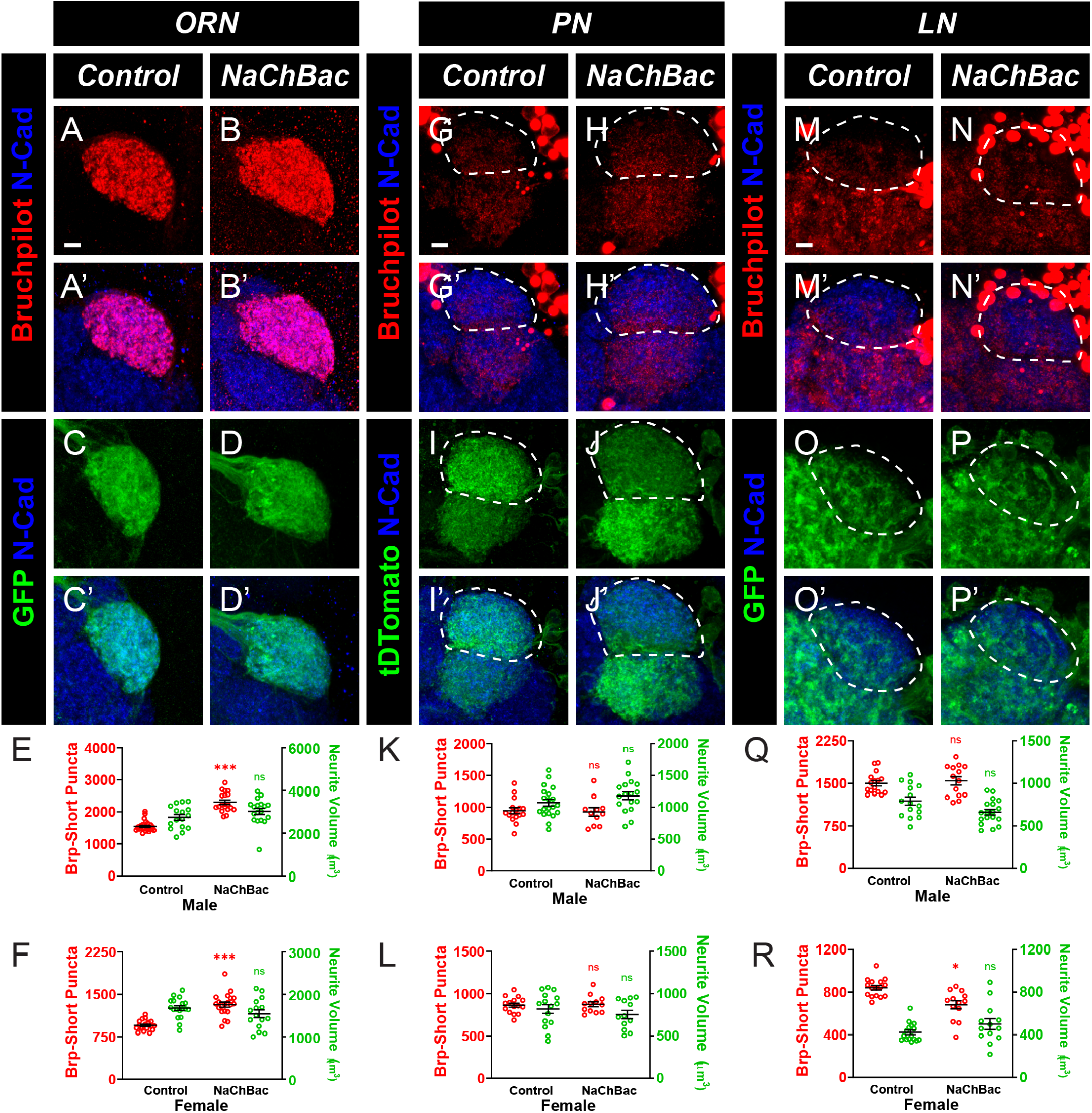
Increased neuronal activity in ORNs, but not PNs or LNs, results in an increase in synapse number. A-D’, Representative confocal image stacks of 10-day old DA1 ORNs expressing either Brp-Short-mStraw (A-B) or membrane-tagged GFP (C-D) in control flies (A, C) or in flies expressing the NaChBac (B, D) construct to increase neuronal activity. Brains were stained with antibodies against mStraw (red) or GFP (green) and N-Cadherin (blue). E, Quantification of Brp-Short puncta and neurite volume in control and NaChBac expressing male DA1 ORNs. Increasing neuronal activity increased the number of synaptic puncta without affecting neurite volume. F, Quantification of Brp-Short puncta and neurite volume in control and NaChBac expressing female DA1 ORNs. Increasing neuronal activity increased the number of synaptic puncta without affecting neurite volume. G-J’, Representative image stacks of 10-day old DA1 PNs (dashed white lines) expressing Brp-Short-GFP (G-H) or membrane-tagged tdTomato (I-J) in control (G, I) or NaChBac expressing (H, J) flies. Brains were stained with antibodies against tdTomato (green) or GFP (red) as well as N-Cadherin (blue). K, Quantification of Brp-Short puncta and neurite volume for control male DA1 PNs or those expressing NaChBac. Increasing neuronal activity in PNs did not affect synaptic puncta number or neurite volume. L, Quantification of Brp-Short puncta and neurite volume for control female DA1 PNs or those expressing NaChBac. Increasing neuronal activity in PNs did not affect synaptic puncta number or neurite volume. M-P’, Representative image stacks of 10-day old DA1 LNs (dashed white lines) expressing Brp-Short-mStraw (M-N) or membrane-tagged GFP (O-P) in control (M, O) or NaChBac expressing (N, P) flies with antibody staining as in A-D’. Q, Quantification of Brp-Short puncta and neurite volume for male DA1 LNs. Increasing excitability did not affect puncta number or neurite volume. R, Quantification of Brp-Short puncta and neurite volume for female DA1 LNs. Increasing excitability slightly decreased puncta number without affecting neurite volume. For each experimental group, n ≥ 11 glomeruli from 6 brains. *, p < 0.1, ***, p < 0.001, n.s. = not significant. Scale bar = 5 μm.

### Altering kinase activity affects synapse number differently in ORNs compared to PNs and LNs

Diverse signal transduction and molecular pathways are responsible for translating changes in neuronal activity to cellular modifications in neurons, including altered neurite volume, synapse number, and gene expression (Chiang et al., 2009; Faust et al., 2021; Heinz and Bloodgood, 2020; Lee and Fields, 2021). Our data suggested that perturbing electrical activity had both shared and differential effects on synapse formation across distinct neuronal class (ORN vs. PN vs. LN): reduced electrical activity reduced synapse number in all classes, but increased electrical activity affected synapse number in some (ORN) but not all (PN and LN) neuronal classes. We next sought to identify a molecular determinant that could underlie at least some of the changes we observed in activity-related synapse number. Shaggy (sgg), the Drosophila homologue of the mammalian kinase GSK-3β, functions with neuronal activity to mediate neuronal stability in the CNS (Chiang et al., 2009) and at the NMJ to regulate bouton formation and microtubule organization (Franco et al. 2004; Ataman et al. 2008; Miech et al. 2008). In olfactory neurons, there is an inverse relationship between neuronal activity and sgg kinase levels, which influences neuronal stability (Chiang et al., 2009). Our data indicates that ORN, PN, and LN synapse number are all reduced when neuronal activity is decreased (Figure 6). Thus, we hypothesized that if Shaggy was the relevant downstream kinase to mediate these changes in synapse number (and given the inverse relationship between neuronal activity and kinase levels), overexpression of a constitutively active sgg (sgg-CA) would result in a similar reduction in synapse number. To test this, we expressed a constitutively active variant of sgg (Sgg-CA; Bourouis, 2002) in all three classes of cells to increase kinase activity along with Brp-Short or mCD8-GFP to quantify synaptic puncta number and neurite volume. Consistent with this hypothesis, Sgg-CA expression in ORNs or LNs that innervate the DA1 glomerulus both similarly resulted in a cell autonomous decrease in Brp-Short puncta number in male (Figure 8A-B’, E, G-H’, K) and female (Figure 8F, L; Figure 8 – Figure Supplement 2A-A’, C-C’, K-K’,M-M’) brains. In male and female ORNs, neurite volume was unaffected (Figure 8C-D’, E-F; Figure 8 – Figure Supplement 2D-D’, F-F’) while in LNs, there was a 28% decrease in neurite volume in males while female LN neurite volume was unaffected (Figure 8I-J’, K-L; Figure 8 – Figure Supplement 2N-N’, P-P’). We were unable to assess the effects of Sgg-CA expression on Brp puncta number in PNs as expression via Mz19-GAL4 was lethal. This is likely due to constitutive developmental expression causing lethality, as previously observed (Berdnik et al. 2006). This ORN and LN data, however, supports the hypothesis that electrical activity may signal through the GSK-3β homologue Shaggy to regulate olfactory synapse number across all observed classes of antennal lobe neurons. However, as male LN neurite volume is affected by Sgg activity but not neuronal activity (and vice versa for female LN neurite volume), there are also likely Sgg-dependent, activity-independent and Sgg-independent, activity-dependent phenomena that regulate neuronal growth.

**Figure 8.**
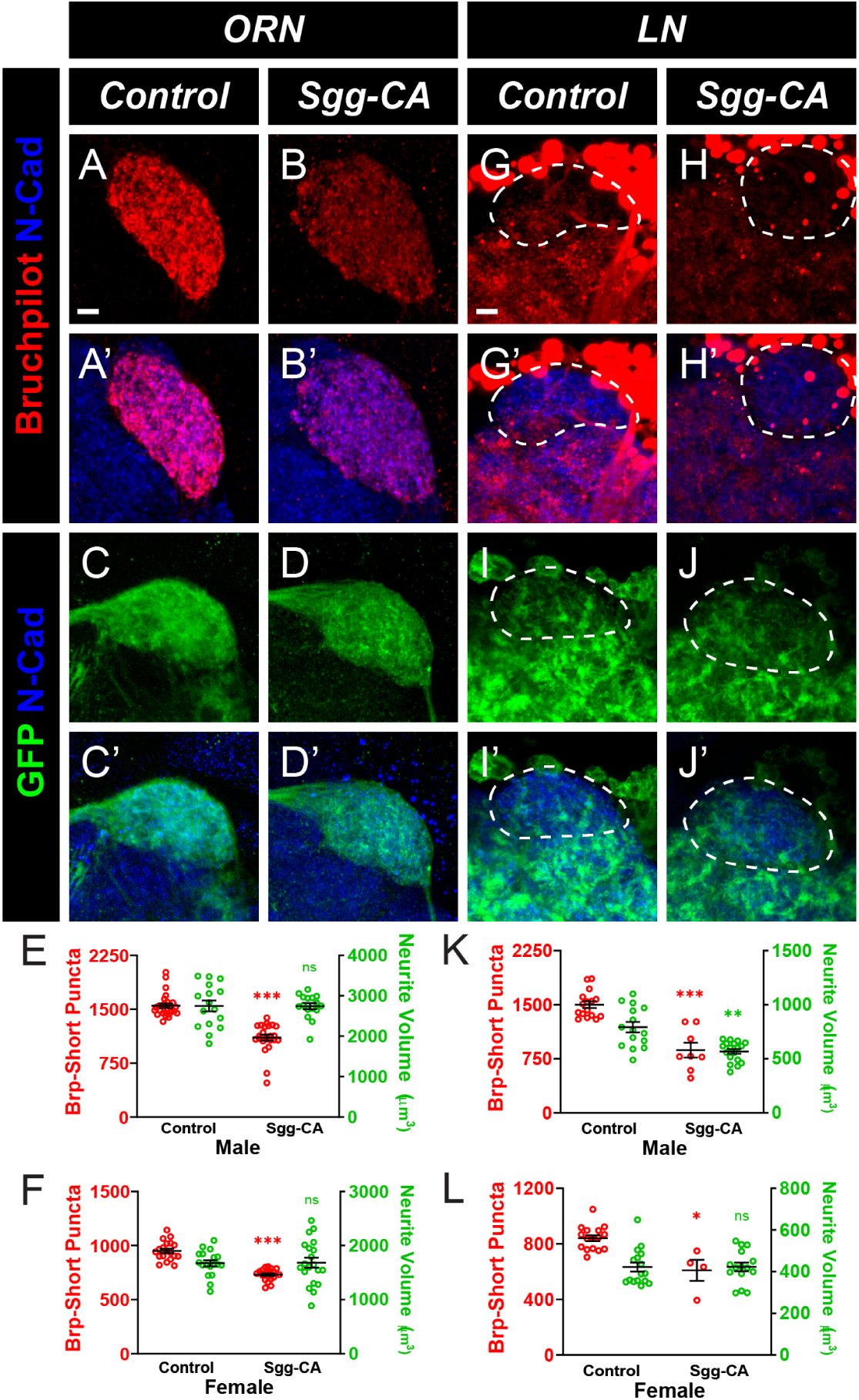
Increased kinase activity decreases ORN and LN synapse number. A-D’, Representative confocal image stacks of 10 day old ORNs of the DA1 glomerulus expressing Brp-Short-mStraw (A-B’) or membrane-tagged GFP (C-D’) in control flies (A, C) or in flies expressing a constitutively active (B, D) variant of Shaggy (GSK3β) to increase overall kinase activity. Brains were stained with antibodies against mStraw (red) or GFP (green) and N-Cadherin (blue). E, Quantification of Brp-Short synaptic puncta and membrane GFP volume in control and Sgg-CA expressing male DA1 ORN. Increasing kinase activity in ORNs caused a significant decrease in synaptic puncta but did not affect neurite volume. F, Quantification of Brp-Short synaptic puncta and membrane GFP volume in female DA1 ORNs. Increasing kinase activity in ORNs caused a significant decrease in synaptic puncta but did not affect neurite volume. G-J’, Representative confocal image stacks of 10-day old DA1 LNs (dashed white lines) expressing Brp-Short-mStraw (G-H) or membrane-tagged GFP (I-J) in control flies (G, I) or in flies expressing constitutively active (H, J) Shaggy and stained with antibodies as in A-D’. K, Quantification of Brp-Short puncta and neurite volume for control and Sgg-CA expressing male DA1 LNs. Increased activity decreased both puncta number and neurite volume in male DA1 LNs. L, Quantification of Brp-Short puncta and neurite volume for female DA1 LNs. Increased activity decreased puncta number without affecting neurite volume in female LNs. For each experimental group, n ≥ 4 glomeruli from 2 brains. *, p < 0.1, **, p < 0.01, ***, p < 0.001, n.s. = not significant. Scale bar = 5 μm.

We similarly assessed the relationship between neuronal activity and kinase levels by expressing a dominant negative sgg transgene (sgg-DN) to decrease kinase levels (Franco et al., 2004). Because of the inverse relationship between the two activity levels, we hypothesized that decreasing Shaggy would phenocopy increasing neuronal activity. Intriguingly, however, decreasing kinase activity in ORNs decreased Brp-Short puncta in male (Figure 8 – Figure Supplement 1A-B’, E) and female (Figure 8 – Figure Supplement 1F; Figure 8 – Figure Supplement 2A-B’) brains by 27% and 30%, respectively. However, decreasing sgg levels in PNs and LNs did not affect Brp-Short puncta in males (Figure 8 – Figure Supplement 1G-H’, K, M-N’, Q) and females (Figure 8 – Figure Supplement 1L, R; Figure 8 – Figure Supplement 2G-H’, K-L’), matching the effects of increased excitability in these neuron classes. Decreased kinase activity increased neurite volume 26% in male PNs but did not affect ORNs or LNs (Figure 8 – Figure Supplement 1C-D’, E, I-J’, K, O-P’, Q) nor any class of neuron in females (Figure 8 – Figure Supplement 1F, L, R; Figure 8 – Figure Supplement 2D-E’, I-J’, N-O’). This suggests additional heterogeneity in the molecular cascades controlling neuronal and synapse development as well as sexual dimorphism. Overall, our data shows that development is not a uniform process between neuronal classes, even in neurons belonging to the same microcircuit with the same general processing goal, and that each class of neuron forms synapses using its own unique temporal, activity-dependent, and molecular programs. Thus, considerable temporal and molecular heterogeneity exists in the development of central synaptic organization.

## DISCUSSION

Synapse formation and maintenance are essential processes that begin during nervous system development and continue over the course of an animal’s lifetime. Understanding how synapse organization changes during developmental and adult times is critical to understanding how different classes of cells employ temporal, molecular, and cellular programs to achieve proper development. As neuropsychiatric, neurodevelopmental, and even neurodegenerative disorders affect circuits at different times during the life cycle of an animal (Bennett, 2011; Bonansco and Fuenzalida, 2016; Dabool et al., 2019; Grant, 2012; Ilieva et al., 2009; Mullins et al., 2016), a clear picture of how different neurons, even neurons within the same circuit, develop is key to understanding how those disorders can influence circuit biology. Here, we have determined the first comprehensive timelines of active zone formation for three Drosophila olfactory neuron classes within the same microcircuit of the antennal lobe: olfactory receptor neurons, projection neurons, and local interneurons. Moreover, we also determined the developmental synaptic time courses for different subtypes of olfactory receptors that sense either food or pheromone odors. We find that each distinct class of olfactory neurons, despite having the same overarching goal – to process olfactory information from the environment and transfer it to higher order brain centers to inform behavioral decisions, employs non-identical temporal programs to accomplish synaptic development. We also demonstrate that development is still occurring during adult stages, up to as late as 6 days after eclosion. Further, our data suggests that all three antennal lobe neuron classes share an activity-dependent component that influences synaptic development whereby reduced activity results in increased levels of GSK-3β, resulting in fewer synapses.

Only ORNs increase synapse number in response to increased activity while synapse number in LNs and PNs remain unchanged following increased activity, suggesting divergence in activity-dependent mechanisms based on antennal lobe neuron class. Taken together, this indicates that synaptic development is not a uniform process for all neurons, with each neuron having its own time-course of synapse formation depending on its class or subtype and that each class uses activity-dependent and activity-independent mechanisms differently to complete synaptic development. These results raise the importance of understanding the precise events in synaptic development for each class of neurons and the limitations of generalizing findings from one connection or neuronal class to another. By obtaining more detailed knowledge of how each neuron class develops in relation to others, including those within its own circuits, we gain a better understanding of each individual neuron class and can begin to determine the underlying genetic and molecular mechanisms that underlie these differences. This is essential to future developmental, molecular, and even degenerative studies to understand which and when events go awry during developmental disorders, why certain classes of neurons are more susceptible to damage or disease than others, and how different cells are organized to coordinate brain development.

Our findings suggest that developmental timing is not identical when comparing different classes or even subtypes of antennal lobe neurons. Each group of neurons has subtle, but clear, variations in how synapses are added or removed over time depending on its class and the type of olfactory information it relays. What is the evolutionary or ethological benefit of this? Why employ multiple temporal modes of development for the component neurons of a circuit that function together to promote a single sensory function? And why employ further developmental programs with different subtypes of the same neuronal class? ORNs that innervate the non-sexually dimorphic DL4 glomerulus are responsible for detecting food-based odorants (Couto et al., 2005; Endo et al., 2007). DL4 ORNs have a period of synapse formation from mid-pupation to 6 days post-eclosion followed by a plateau, thus maintaining a steady state number of synapses over the entire time observed (Figure 3 – Figure Supplements 2 and 3). In contrast to this, ORNs of the sexually dimorphic DA1 and VA1lm glomeruli, that are responsible for detecting pheromone-based odorants, instead show a period of rapid synaptic addition in early adulthood, followed by a decrease in synapses later in adulthood (Figures 2-3 with associated supplements). In the case of the VA1lm ORNs, this decrease appears to be a pruning stage that is followed by maintenance of a steady state number of synapses. For DA1 ORNs, however, this occurs later in adulthood, perhaps as a period of senescence (Koch et al., 2021). It is tempting to speculate that these temporal patterns of synapse addition and removal may be related to each the specific function of each class of ORNs. ORNs that detect food odorants and innervate non-sexually dimorphic glomeruli may retain the maximum synapse number throughout adulthood to ensure that the sensory abilities necessary to find food always exist. The need to locate and consume nutrients is a need that persists at all stages of the adult fly. ORNs that innervate sexually dimorphic glomeruli, like DA1 and VA1lm, may instead lose connections as the animal ages because older flies exhibit reduced fecundity (Flatt, 2020) and the energetic cost of maintaining high synaptic numbers in a mate-seeking circuit may no longer be favored. Further, when PNs and LNs are considered with food-sensing or pheromone-sensing ORNs, why are there additional differences in developmental time courses? All ORN classes examined here and PNs increase synapses during the first 5-6 days of adult life before either plateauing at a steady state or pruning, then plateauing at a steady state (Figures 2-4). LNs, however, develop their mature synaptic complement very quickly during pupation and are largely maintained from eclosion throughout adult life (Figure 5). Female flies make the decision to lay their eggs in areas of resource richness (Aranha and Vasconcelos, 2018) to ensure that their progeny are born in areas that promote their survival. If olfactory acuity is connected to the number of synapses, and they spend their initial days of adult life in an area where there is ample food supply and other flies from other egg-laying females nearby, they may not have to rely on olfaction as much to find food and mates due to simple proximity to both during those initial days. However, when food resources become scarcer in the days following birth due to usage, and adult flies begin to travel further apart, the greater olfactory acuity afforded by an increase in synapse number is necessary to find new food sources and to locate mates over a larger geographical space. In such a situation, LNs may be immediately needed to control gain between ORNs and PNs and appropriate communication. ORNs and PNs, however, can attain their mature synapse number more slowly as peak acuity for transfer of the odorant information into the brain and conveyance of that information to higher brain centers isn’t needed until later in adult life. Further, as LNs regulate gain control of the AL circuit through connections with PNs and other LNs throughout adulthood, it is likely necessary for their synaptic connections to remain numerous throughout the fly’s life without decline. Finally, the temporal differences may further be influenced by prior wiring events. There is a distinct order to ORN, PN, and LN wiring during pupation (Chou et al., 2010; Liou et al., 2018; Sakuma et al., 2014; Wilson, 2013) and though targeting events are largely complete by 48h APF, neurite growth continues throughout pupal stages. With respect to synapse addition, PNs and ORNs had similar periods of synapse addition and plateau, but PN stages typically occurred slightly later than in ORNs. As LNs are the major targets of these connections (Schlegel et al., 2021), their later timing may be a result of the LNs being the last of the AL neurons to complete synaptic development. The PNs cannot begin forming pre-synapses until the LN projections have entered the glomerulus and have begun forming synapses of their own. Therefore, ORNs can begin forming synapses with the PNs immediately after they enter the glomerulus, but PN synapse formation may not be able to commence until LNs begin forming connections. In this consideration, the synaptic development of one class of neuron is partially dependent on the development of the other neuron classes; this will be an interesting avenue for future study. Overall, these detailed time-courses demonstrate that even within the same circuit, different classes and subtypes of neurons have notable differences in their temporal programs of synaptic development. This highlights a previously unappreciated diversity in the development of central neurons and suggests that function, timing, and role may influence even similar neuron classes to use variant temporal programs during development to achieve mature circuit function.

Beyond temporal differences in the developmental programs used by different olfactory neuron classes, we also identified both similarities and differences in the activity-dependent and activity-independent molecular influences on synapse development. In ORNs, PNs, and LNs, we found that reduced neuronal activity results in fewer synapses, suggesting that a shared, baseline level of neuronal activity is generally necessary for complete synaptic development (Figure 6). Previous work in the antennal lobe (Chiang et al., 2009) showed that decreased neuronal activity leads to increased activity of shaggy, the Drosophila GSK3β homologue and leads to less axonal stability. Consistent with this, we find that overexpression of Sgg phenocopies the effects of reduced neuronal activity in ORNs and LNs (Figure 8, Figure 8 – figure supplement 2; PN overexpression of Sgg is pupal lethal, precluding our analysis). In females, we did not observe any alterations to axonal stability with Sgg-CA overexpression, as neurite volume in ORNs and LNs remains unchanged. However, male LN neurite volume was affected by Sgg-CA expression. This suggests that this activity-dependent requirement for a baseline activity that functions through Sgg levels may be a common mechanism across olfactory neurons to control the development of active zone number. There is, however, a sexual dimorphism as to whether neurite volume is regulated by Sgg activity. Reduced electrical activity that results in increased kinase activity likely leads to more phosphorylation of downstream targets, resulting in removal of synapses. Whether this is a completely cell autonomous phenomenon or is circuit-based, remains unclear. It will be important to test in the future whether different classes of neurons alter their synaptic development or growth when the activity of another neuronal class is impaired, possibly due to competition (Miller, 1996). Antennal lobe neurons showed different responses to increased electrical activity, however. Expression of NaChBac, a bacterial Na+ channel that increases neuronal firing, in ORNs resulted in more ORN synapses while either PN or LN expression of NaChBac had no effect on synapse number in those neurons (Figure 7, Figure 7 – figure supplement 1). This suggests that PNs and LNs are more resistant to activity increases compared to ORNs and that this may be an element of activity-independent regulation of synapse number. Alternatively, this may also indicate an upper limit on synapse number or to the extent by which neuronal activity influences synapse number. In ORNs, the downstream mechanism of how increased activity influences synapse number remains unclear. If a direct relationship existed between activity and GSK-3β activity, we would expect that reducing kinase levels via expression of Sgg-DN would phenocopy the effects of NaChBac expression. Instead, Sgg-DN expression cell autonomously decreases synapse number in ORNs (Figure 8 – figure supplements 1 and 2). This highlights two major points: 1) there is an optimal level of Sgg activity in ORNs that influences synapse number and perturbations away from that optimal level impair synapse formation, and 2) there is an additional, yet undiscovered, mechanism that connects increased ORN activity with increased synapse number. In PNs and LNs, increased activity does not result in any changes to synapse number. Consistent with neuronal activity functioning through Sgg in these neurons, Sgg-DN expression in PNs or LNs also has no effect on synapse number. This represents a different mechanism for PNs and LNs than that which governs ORN synapse formation. This alternative mechanism for ORN synapse formation further highlights that the process of synaptic development is not uniform across all neuron types. Future work will therefore tease apart the activity-dependent and activity-independent mechanisms that influence synapse development in each class of neurons to understand how neuronal activity influences downstream cellular events, such as regulating kinase activity, to promote synapse formation and maintenance in distinct classes of neurons. Overall, our findings demonstrate the first synapse-level, quantitative analysis of presynaptic active zone development in each neuron class of a sensory circuit. These data increase our understanding of the rules that govern synaptic development and highlight critical caveats that should be considered for all future analyses of synaptic development and function. Developmental processes need to be uniquely understood for each class of neurons studied as our data indicates that there is no “one size fits all” rule for different classes of CNS neurons, even within the same circuit. Moreover, as synaptic development continues to occur during the adult stage, behavioral analyses need to be completed with appropriately age-matched controls and not spanning a range of ages, as this would compare neuronal function between neurons with different complements of presynaptic active zones and at different stages in their development. Taken together, these findings provide an essential bedrock from which future questions on the causes and consequences of incorrect development can be built and answered.

## ACKNOWLEDGMENTS

We would like to thank Dr. Michael Parisi, Dr. Kristen Davis, Dr. Juan Carlos Duhart, Dr. Stephen Tymanskyj, Jesse Humenik, Dr. Le Ma, and S. Zosimus for stimulating discussions and comments on the manuscript. We also thank the Bloomington Stock Center for flies and the Developmental Studies Hybridoma Bank (University of Iowa, created by the NICHD) for antibodies. This work was supported by the US National Institute of Health grant R00-DC013059 and the Commonwealth Universal Research Enhancement (CURE) program of the Pennsylvania Department of Health grant 4100077067 (to TJM). Aspects of this work and general work in the TJM Lab are supported by grants from the Alfred P. Sloan Foundation, Whitehall Foundation, Jefferson Dean’s Transformational Science Award, Jefferson Synaptic Biology Center, and Thomas Jefferson University start-up funds.

## METHODS

### Drosophila stocks and transgenic lines

All control lines and genetic fly stocks were maintained on cornmeal::dextrose medium (Archon Scientific, Durham, NC) at 21°C while crosses were raised on similar medium at 25°C (unless noted in the text) in incubators (Darwin Chambers, St. Louis, MO) at 60% relative humidity with a 12/12 light/ dark cycle. Transgenes were maintained over balancers with fluorescent markers and visible phenotypic traits to allow for selection of pupae or adults of the desired genotype. To drive expression in specific classes of antennal lobe neurons, we used the following GAL4 lines: AM29-GAL4 (Endo et al., 2007), Or47b-GAL4 (Vosshall et al., 2000), Or67d-GAL4 (Stockinger et al., 2005), Mz19-GAL4 (Jefferis et al., 2004), and NP3056-GAL4 (Chou et al., 2010). The following UAS transgenes were used as synaptic labels or to express molecular constructs for genetic perturbation experiments: UAS-Brp-Short-mStraw (Fouquet et al., 2009; Mosca and Luo, 2014), UAS-Brp-Short-GFP (Schmid et al., 2008), UAS-mCD8-GFP (Lee and Luo, 1999), UAS-3xHA-mtdTomato (Potter et al., 2010), UAS-NaChBac (Nitabach et al., 2006), UAS-Sgg-DN (Franco, 2004), UAS-Sgg-CA (Bourouis, 2002), UAS-TNT (Sweeney et al., 1995), and UAS-TNT-Imp (Sweeney et al., 1995).

### Immunocytochemistry

For time-course experiments, male and female pupal brains were dissected at 48, 60, 72, 84, and 92 hours after puparium formation (APF) and adult brains were dissected at 0, 3, 6, 9, 12, 15, and 18 days post eclosion. To collect pupae at 0-hour APF, white pre-pupae were selected from vials based on genotype using fluorescent balancer markers and aged to the desired time-point. For adults, flies were cleared from vials one day before collection and on the following day, newly eclosed adults were chosen based on genotype using identifiable balancers and phenotypic markers. Flies were then aged to the desired time-point before dissection and immunostaining. For experiments involving genetic manipulation of either neuronal or kinase activity, adult brains were dissected at 10 days post eclosion. Brains were fixed in 4% paraformaldehyde for 20 minutes before being washed in phosphate buffer (1x PB) with 0.3% Triton (PBT). Brains were then blocked for an hour in PBT containing 5% normal goat serum (NGS) before being incubated in primary antibodies diluted in PBT with 5% NGS for two days at 4°C. Following staining, primary antibodies were discarded and the brains washed 3 × 20’ with PBT and incubated in secondary antibodies diluted in PBT with 5% NGS for an additional two days at 4°C. The secondary antibodies were then discarded, the brains washed 3 × 20’ in PBT, and then incubated overnight in SlowFade(tm) (ThermoFisher Scientific, Waltham, MA) gold antifade mounting media and allowed to sink. Brains were then mounted in SlowFade mounting media using a bridge-mount method with No. 1 cover glass shards and stored at 20°C before being imaged (Wu and Luo, 2006). The following primary antibodies were used: mouse anti-Nc82 (Laissue et al., 1999) rabbit anti-DsRed (TaKaRa Bio, cat. no. 632496, 1:250; Mosca and Luo, 2014), chicken anti-GFP (Aves, cat. no. GFP-1020, 1:1000; Mosca and Luo, 2014), and rat anti-N-cadherin (Hummel and Zipursky, 2004). Alexa568-(ThermoFisher Scientific, Waltham, MA) and Alexa647-conjugated (Jackson ImmunoResearch, West Grove, PA) secondary antibodies were used at 1:250 while FITC-conjugated (Jackson ImmunoResearch, West Grove, PA) secondary antibodies were used at 1:200. In some cases, non-specific background recognized by the dsRed antibodies (in the form of large red spots appear around the antennal lobes and outside of the tissue observed). These are part of the background, are not caused by any of the transgenic constructs used (Mosca and Luo, 2014) and did not influence any quantification or scoring methods (see below).

### Imaging, analysis, and image processing

All images were obtained using a Zeiss LSM880 Laser Scanning Confocal Microscope (Carl Zeiss, Oberlochen, Germany) using a 40X 1.4 NA Plan-Apochromat lens or a 63X 1.4 NA Plan-Apochromat f/ELYRA lens at an optical zoom of 3x. Images were centered on the glomerulus of interest and the z-boundaries were set based on the appearance of the synaptic labels, Brp-Short-mStraw and mCD8-GFP. Images were analyzed three dimensionally using the Imaris Software 9.3.1 (Oxford Instruments, Abingdon, UK) on a custom-built image processing computer (Digital Storm, Fremont, CA) following previously established methods (Mosca and Luo, 2014; Mosca et al., 2017). Brp-Short puncta were quantified using the “Spots” function with a spot size of 0.6 µm. Neurite volume was quantified using the “Surfaces” function with a local contrast of 3 µm and smoothing of 0.2 µm for AM29, Or47b, and Or67d ORNs or a local contrast of 0.5 and a smoothing of 0.2 µm for Mz19 PNs and NP3056 LNs. The resultant masks were then visually inspected to ensure their conformation to immunostaining. Images were processed using ImageJ (NIH, Bethesda, MD) and Adobe Photoshop 2020 (Adobe Systems, San Jose, CA). Figures were produced using Adobe Illustrator CC 2019 (Adobe Systems, San Jose, CA).

### Statistical Analysis

All data was analyzed using Prism 8 (GraphPad Software, Inc., La Jolla, CA). This software was also used to generate graphical representations of data. Unpaired Student’s t-tests were used to determine significance between two groups while one-way ANOVA with Tukey’s multiple comparisons tests were used to determine significance between groups of three or more. A p-value of 0.05 was set as the threshold for significance in all studies. For experiments involving disruption of neuronal activity or kinase activity (Figures 6-8, Figures 6-8 Figure Supplements), multiple concurrent experiments were performed with experimental genotypes as well as wild-type flies and appropriate controls for each genotype. This approach necessitated the need for one-way ANOVA with Tukey’s multiple comparisons tests. For each figure, informative genotypes have been presented along with controls appropriate for each genotype.

## REFERENCES

1. Akin, O., and Zipursky, S.L. (2020). Activity regulates brain development in the fly. Curr. Opin. Genet. Dev. 65, 8–13.

2. Aranha, M.M., and Vasconcelos, M.L. (2018). Deciphering Drosophila female innate behaviors. Curr. Opin. Neurobiol. 52, 139–148.

3. Axel, R., Wong, A.M., Suh, G.S.B., Hergarden, A.C., Anderson, D.J., Wang, J.W., Benzer, S., and Simon, A.F. (2004). A single population of olfactory sensory neurons mediates an innate avoidance behaviour in Drosophila. Nature 431, 854–859.

4. Baines, R.A., and Bate, M. (1998). Electrophysiological development of central neurons in the Drosophila embryo. J. Neurosci. 18, 4673–4683.

5. Bennett, M.R. (2011). Schizophrenia: Susceptibility genes, dendritic-spine pathology and gray matter loss. Prog. Neurobiol. 95, 275–300.

6. Berger-Müller, S., Sugie, A., Takahashi, F., Tavosanis, G., Hakeda-Suzuki, S., and Suzuki, T. (2013). Assessing the role of cell-surface molecules in central synaptogenesis in the Drosophila visual system. PLoS One 8, 1–14.

7. Bonansco, C., and Fuenzalida, M. (2016). Plasticity of hippocampal excitatory-inhibitory balance: Missing the synaptic control in the epileptic brain. Neural Plast. 2016.

8. Bourouis, M. (2002). Targeted increase in Shaggy activity levels blocks wingless signaling. Genesis 34, 99–102.

9. Brand, A.H., and Perrimon, N. (1993). Targeted gene expression as a means of altering cell fates and generating dominant phenotypes. Development 118, 401–415.

10. Cang, J., and Feldheim, D.A. (2013). Developmental mechanisms of topographic map formation and alignment. Annu. Rev. Neurosci. 36, 51–77.

11. Chiang, A., Priya, R., Ramaswami, M., VijayRaghavan, K., and Rodrigues, V. (2009). Neuronal activity and Wnt signaling act through Gsk3-β to regulate axonal integrity in mature Drosophila olfactory sensory neurons. Development 136, 1273–1282.

12. Chou, Y.H., Spletter, M.L., Yaksi, E., Leong, J.C.S., Wilson, R.I., and Luo, L. (2010). Diversity and wiring variability of olfactory local interneurons in the Drosophila antennal lobe. Nat. Neurosci. 13, 439–449.

13. Christiansen, F., Zube, C., Andlauer, T.F.M., Wichmann, C., Fouquet, W., Owald, D., Mertel, S., Leiss, F., Tavosanis, G., Luna, A.J.F., et al. (2011). Presynapses in Kenyon cell dendrites in the mushroom body calyx of Drosophila. J. Neurosci. 31, 9696–9707.

14. Coates, K.E., Majot, A.T., Zhang, X., Michael, C.T., Spitzer, S.L., Gaudry, Q., and Dacks, A.M. (2017). Identified serotonergic modulatory neurons have heterogeneous synaptic connectivity within the olfactory system of drosophila. J. Neurosci. 37, 7318– 7331.

15. Coates, K.E., Calle-Schuler, S.A., Helmick, L.M., Knotts, V.L., Martik, B.N., Salman, F., Warner, L.T., Valla, S. V, Bock, D.D., and Dacks, A.M. (2020). The Wiring Logic of an Identified Serotonergic Neuron That Spans Sensory Networks. J. Neurosci. 40, 6309–6327.

16. Collins, C.A., and DiAntonio, A. (2007). Synaptic development: insights from Drosophila. Curr. Opin. Neurobiol. 17, 35–42.

17. Couto, A., Alenius, M., and Dickson, B.J. (2005). Molecular, anatomical, and functional organization of the Drosophila olfactory system. Curr. Biol. 15, 1535–1547.

18. Dabool, L., Juravlev, L., Hakim-Mishnaevski, K., and Kurant, E. (2019). Modeling Parkinson’s disease in adult Drosophila. J. Neurosci. Methods 311, 89–94.

19. Dalva, M.B., McClelland, A.C., and Kayser, M.S. (2007). Cell adhesion molecules: Signalling functions at the synapse. Nat. Rev. Neurosci. 8, 206–220.

20. Datta, S.R., Vasconcelos, M.L., Ruta, V., Luo, S., Wong, A., Demir, E., Flores, J., Balonze, K., Dickson, B.J., and Axel, R. (2008). The Drosophila pheromone cVA activates a sexually dimorphic neural circuit. Nature 452, 473–477.

21. Depew, A.T., and Mosca, T.J. (2021). Conservation and innovation: Versatile roles for lrp4 in nervous system development. J. Dev. Biol. 9.

22. DePew, A.T., Aimino, M.A., and Mosca, T.J. (2019). The tenets of teneurin: Conserved mechanisms regulate diverse developmental processes in the Drosophila nervous system. Front. Neurosci. 13, 1–11.

23. Endo, K., Aoki, T., Yoda, Y., Kimura, K.I., and Hama, C. (2007). Notch signal organizes the Drosophila olfactory circuitry by diversifying the sensory neuronal lineages. Nat. Neurosci. 10, 153–160.

24. Farhy-Tselnicker, I., and Allen, N.J. (2018). Astrocytes, neurons, synapses: A tripartite view on cortical circuit development. Neural Dev. 13, 1–12.

25. Faust, T.E., Gunner, G., and Schafer, D.P. (2021). Mechanisms governing activity-dependent synaptic pruning in the developing mammalian CNS. Nat. Rev. Neurosci. 22, 657–673.

26. Fishilevich, E., and Vosshall, L.B. (2005). Genetic and functional subdivision of the Drosophila antennal lobe. Curr. Biol. 15, 1548– 1553.

27. Flatt, T. (2020). Life-history evolution and the genetics of fitness components in drosophila melanogaster. Genetics 214, 3–48.

28. Fouquet, W., Owald, D., Wichmann, C., Mertel, S., Depner, H., Dyba, M., Hallermann, S., Kittel, R.J., Eimer, S., and Sigrist, S.J. (2009). Maturation of active zone assembly by Drosophila Bruchpilot. J. Cell Biol. 186, 129–145.

29. Franco, B. (2004). Shaggy, the Homolog of Glycogen Synthase Kinase 3, Controls Neuromuscular Junction Growth in Drosophila. J. Neurosci. 24, 6573–6577.

30. Franco, B., Bogdanik, L., Bobinnec, Y., Debec, A., Bockaert, J., Parmentier, M.-L., and Grau, Y. (2004). Shaggy, the homolog of glycogen synthase kinase 3, controls neuromuscular junction growth in Drosophila. J. Neurosci. 24, 6573–6577.

31. Goda, Y., and Davis, G.W. (2003). Mechanisms of synapse assembly and disassembly. Neuron 40, 243–264.

32. Grabe, V., and Sachse, S. (2018). Fundamental principles of the olfactory code. BioSystems 164, 94–101.

33. Grabe, V., Baschwitz, A., Dweck, H.K.M., Lavista-Llanos, S., Hansson, B.S., and Sachse, S. (2016). Elucidating the Neuronal Architecture of Olfactory Glomeruli in the Drosophila Antennal Lobe. Cell Rep. 16, 3401–3413.

34. Grant, S.G.N. (2012). Synaptopathies: Diseases of the synaptome. Curr. Opin. Neurobiol. 22, 522–529.

35. Hallem, E.A., and Carlson, J.R. (2006). Coding of Odors by a Receptor Repertoire. Cell 125, 143–160.

36. Harris, K.P., and Littleton, J.T. (2015). Transmission, development, and plasticity of synapses. Genetics 201, 345–375.

37. Hazan, L., and Ziv, N.E. (2020). Activity Dependent and Independent Determinants of Synaptic Size Diversity. J. Neurosci. 40, 2828– 2848.

38. Heinz, D.A., and Bloodgood, B.L. (2020). Mechanisms that communicate features of neuronal activity to the genome. Curr. Opin. Neurobiol. 63, 131–136.

39. Hildebrand, J.G., and Shepherd, G.M. (1997). Mechanisms of olfactory discrimination: Converging evidence for common principles across phyla. Annu. Rev. Neurosci. 20, 595–631.

40. Hong, E.J., and Wilson, R.I. (2015). Simultaneous encoding of odors by channels with diverse sensitivity to inhibition. Neuron 85, 573–589.

41. Hummel, T., and Rodrigues, V. (2008). Development of the Drosophila olfactory system. Adv. Exp. Med. Biol. 628, 82–101.

42. Hummel, T., and Zipursky, S.L. (2004). Afferent induction of olfactory glomeruli requires N-cadherin. Neuron 42, 77–88.

43. Ilieva, H., Polymenidou, M., and Cleveland, D.W. (2009). Non-cell autonomous toxicity in neurodegenerative disorders: ALS and beyond. J. Cell Biol. 187, 761–772.

44. Jarecki, J., and Keshishian, H. (1995). Role of neural activity during synaptogenesis in Drosophila. J. Neurosci. 15, 8177–8190.

45. Jefferis, G.S.X.E., and Hummel, T. (2006). Wiring specificity in the olfactory system. Semin. Cell Dev. Biol. 17, 50–65.

46. Jefferis, G.S.X.E., Marin, E.C., Stocker, R.F., and Luo, L. (2001). Target neuron prespecification in the olfactory map of Drosophila. Nature 414, 204–208.

47. Jefferis, G.S.X.E., Vyas, R.M., Berdnik, D., Ramaekers, A., Stocker, R.F., Tanaka, N.K., Ito, K., and Luo, L. (2004). Developmental origin of wiring specificity in the olfactory system of Drosophila. Development 131, 117–130.

48. Jefferis, G.S.X.E., Potter, C.J., Chan, A.M., Marin, E.C., Rohlfing, T., Maurer, C.R., and Luo, L. (2007). Comprehensive Maps of Drosophila Higher Olfactory Centers: Spatially Segregated Fruit and Pheromone Representation. Cell 128, 1187–1203.

49. Keshishian, H., Broadie, K., Chiba, A., and Bate, M. (1996). The Drosophila neuromuscular junction: A model system for studying synaptic development and function. Annu. Rev. Neurosci. 19, 545–575.

50. Koch, S.C., Nelson, A., and Hartenstein, V. (2021). Structural aspects of the aging invertebrate brain. Cell Tissue Res. 383, 931–947.

51. Komiyama, T., and Luo, L. (2006). Development of wiring specificity in the olfactory system. Curr. Opin. Neurobiol. 16, 67–73.

52. Kremer, M.C., Christiansen, F., Leiss, F., Paehler, M., Knapek, S., Andlauer, T.F.M., Förstner, F., Kloppenburg, P., Sigrist, S.J., and Tavosanis, G. (2010). Structural long-term changes at mushroom body input synapses. Curr. Biol. 20, 1938–1944.

53. Kummer, T.T., Misgeld, T., and Sanes, J.R. (2006). Assembly of the postsynaptic membrane at the neuromuscular junction: Paradigm lost. Curr. Opin. Neurobiol. 16, 74–82.

54. Kurtovic, A., Widmer, A., and Dickson, B.J. (2007). A single class of olfactory neurons mediates behavioural responses to a Drosophila sex pheromone. Nature 446, 542–546.

55. Laissue, P.P., Reiter, C., Hiesinger, P.R., Halter, S., Fischbach, K.F., and Stocker, R.F. (1999). Three-Dimensional Reconstruction of the Antennal Lobe in. J. Comp. Neurol. 552, 543–552.

56. Lee, P.R., and Fields, R.D. (2021). Activity-Dependent Gene Expression in Neurons. Neuroscientist 27, 355–366.

57. Lee, T., and Luo, L. (1999). Mosaic analysis with a repressible neurotechnique cell marker for studies of gene function in neuronal morphogenesis. Neuron 22, 451–461.

58. Li, A., Rao, X., Zhou, Y., and Restrepo, D. (2019). Complex neural representation of odor information in the olfactory bulb. Acta Physiol. e13333.

59. Li, L., Xiong, W.C., and Mei, L. (2018). Neuromuscular Junction Formation, Aging, and Disorders. Annu. Rev. Physiol. 80, 159–188.

60. Li, M., Cui, Z., Niu, Y., Liu, B., Fan, W., Yu, D., and Deng, J. (2010). Synaptogenesis in the developing mouse visual cortex. Brain Res. Bull. 81, 107–113.

61. Lin, D.M., and Goodman, C.S. (1994). Ectopic and increased expression of fasciclin II alters motoneuron growth cone guidance. Neuron 13, 507–523.

62. Liou, N.F., Lin, S.H., Chen, Y.J., Tsai, K.T., Yang, C.J., Lin, T.Y., Wu, T.H., Lin, H.J., Chen, Y.T., Gohl, D.M., et al. (2018). Diverse populations of local interneurons integrate into the Drosophila adult olfactory circuit. Nat. Commun. 9.

63. Liu, W., and Chakkalakal, J. V. (2018). The Composition, Development, and Regeneration of Neuromuscular Junctions (Elsevier Inc.).

64. Martin, F., Boto, T., Gomez-Diaz, C., and Alcorta, E. (2013). Elements of olfactory reception in adult Drosophila melanogaster. Anat. Rec. 296, 1477–1488.

65. Masland, R.H. (2004). Neuronal cell types. Curr. Biol. 14, R497– R500.

66. Miller, K.D. (1996). Synaptic economics: Competition and cooperation in synaptic plasticity. Neuron 17, 371–374.

67. Modi, M.N., Shuai, Y., and Turner, G.C. (2020). The Drosophila Mushroom Body: From Architecture to Algorithm in a Learning Circuit. Annu. Rev. Neurosci. 43, 465–484.

68. Mosca, T.J., and Luo, L. (2014). Synaptic organization of the Drosophila antennal lobe and its regulation by the Teneurins. Elife 3, e03726.

69. Mosca, T.J., Luginbuhl, D.J., Wang, I.E., and Luo, L. (2017). Presynaptic LRP4 promotes synapse number and function of excitatory CNS neurons. Elife 6, 115907.

70. Mullins, C., Fishell, G., and Tsien, R.W. (2016). Unifying Views of Autism Spectrum Disorders: A Consideration of Autoregulatory Feedback Loops. Neuron 89, 1131–1156.

71. Ng, M., Roorda, R.D., Lima, S.Q., Zemelman, B. V., Morcillo, P., and Miesenböck, G. (2002). Transmission of olfactory information between three populations of neurons in the antennal lobe of the fly. Neuron 36, 463–474.

72. Nitabach, M.N., Wu, Y., Sheeba, V., Lemon, W.C., Strumbos, J., Zelensky, P.K., White, B.H., and Holmes, T.C. (2006). Electrical hyperexcitation of lateral ventral pacemaker neurons desynchronizes downstream circadian oscillators in the fly circadian circuit and induces multiple behavioral periods. J. Neurosci. 26, 479–489.

73. Nitkin, R., Smith, M., Magill, C., Fallon, J., Yao, Y.-M., Wallace, B., and McMahan, U. (1987). Protein from Torpedo Electric Organ. J. Cell Biol. 105, 2471–2478.

74. Pan, Y., and Monje, M. (2020). Activity shapes neural circuit form and function: A historical perspective. J. Neurosci. 40, 944–954.

75. Potter, C.J., Tasic, B., Russler, E. V., Liang, L., and Luo, L. (2010). The Q system: A repressible binary system for transgene expression, lineage tracing, and mosaic analysis. Cell 141, 536– 548.

76. de Ramon Francàs, G., Zuñiga, N.R., and Stoeckli, E.T. (2017). The spinal cord shows the way – How axons navigate intermediate targets. Dev. Biol. 432, 43–52.

77. Ren, D., Navarro, B., Xu, H., Yue, L., Shi, Q., and Clapham, D.E. (2001). A Prokaryotic Voltage-Gated Sodium Channel. 294, 2372– 2376.

78. Sakano, H. (2020). Developmental regulation of olfactory circuit formation in mice. Dev. Growth Differ. 62, 199–213.

79. Sakuma, C., Anzo, M., Miura, M., and Chihara, T. (2014). Development of olfactory projection neuron dendritesthat contribute to wiring specificity of the Drosophila olfactory circuit. Genes Genet. Syst. 89, 17–26.

80. Sanes, J.R., and Lichtman, J.W. (1999). Development of the vertebrate neuromuscular junction. Annu. Rev. Neurosci. 22, 389– 442.

81. Schlegel, P., Bates, A.S., Stürner, T., Jagannathan, S.R., Drummond, N., Hsu, J., Serratosa Capdevila, L., Javier, A., Marin, E.C., Barth-Maron, A., et al. (2021). Information flow, cell types and stereotypy in a full olfactory connectome. Elife 10, 2020.01.21.911859.

82. Schmid, A., Hallermann, S., Kittel, R.J., Khorramshahi, O., Frölich, A.M.J., Quentin, C., Rasse, T.M., Mertel, S., Heckmann, M., and Sigrist, S.J. (2008). Activity-dependent site-specific changes of glutamate receptor composition in vivo. Nat. Neurosci. 11, 659– 666.

83. Seki, Y., Dweck, H.K.M., Rybak, J., Wicher, D., Sachse, S., and Hansson, B.S. (2017). Olfactory coding from the periphery to higher brain centers in the Drosophila brain. BMC Biol. 15, 56.

84. Shi, L., Fu, A.K.Y., and Ip, N.Y. (2012). Molecular mechanisms underlying maturation and maintenance of the vertebrate neuromuscular junction. Trends Neurosci. 35, 441–453.

85. Siddiqui, T.J., and Craig, A.M. (2011). Synaptic organizing complexes. Curr. Opin. Neurobiol. 21, 132–143.

86. Simi, A., and Studer, M. (2018). Developmental genetic programs and activity-dependent mechanisms instruct neocortical area mapping. Curr. Opin. Neurobiol. 53, 96–102.

87. Stocker, R.F., Lienhard, M.C., Borst, A., and Fischbach, K. (1990). Neuronal architecture of the antennal lobe in Drosophila melanogaster. Cell Tissue Res. 9–34.

88. Stockinger, P., Kvitsiani, D., Rotkopf, S., Tirián, L., and Dickson, B.J. (2005). Neural circuitry that governs Drosophila male courtship behavior. Cell 121, 795–807.

89. Sweeney, S.T., Broadie, K., Keane, J., Niemann, H., and O’Kane, C.J. (1995). Targeted expression of tetanus toxin light chain in Drosophila specifically eliminates synaptic transmission and causes behavioral defects. Neuron 14, 341–351.

90. Tanaka, N.K., Tanimoto, H., and Ito, K. (2008). Neuronal assemblies of the Drosophila mushroom body. J. Comp. Neurol. 508, 711–755.

91. Tanaka, N.K., Ito, K., and Stopfer, M. (2009). Odor-Evoked Neural Oscillations in Drosophila Are Mediated by Widely Branching Interneurons. J. Neurosci. 29, 8595–8603.

92. Tanaka, N.K., Endo, K., and Ito, K. (2012). Organization of antennal lobe-associated neurons in adult Drosophila melanogaster brain. J. Comp. Neurol. 520, 4067–4130.

93. Thummel, C.S. (2001). Molecular Mechanisms of Developmental Timing in C. elegans and Drosophila. Dev. Cell 1, 453–465.

94. Vonhoff, F., and Keshishian, H. (2017). Activity-dependent synaptic refinement: New insights from Drosophila. Front. Syst. Neurosci. 11, 1–9.

95. Vosshall, L.B., Wong, A.M., and Axel, R. (2000). An olfactory sensory map in the fly brain. 102, 147–159.

96. Wagh, D.A., Rasse, T.M., Asan, E., Hofbauer, A., Schwenkert, I., Dürrbeck, H., Buchner, S., Dabauvalle, M.C., Schmidt, M., Qin, G., et al. (2006). Bruchpilot, a protein with homology to ELKS/CAST, is required for structural integrity and function of synaptic active zones in Drosophila. Neuron 49, 833–844.

97. Wilson, R.I. (2013). Early Olfactory Processing in Drosophila : Mechanisms and Principles. Annu. Rev. Neurosci. 36, 217–241.

98. Wilton, D.K., Dissing-Olesen, L., and Stevens, B. (2019). Neuron-Glia Signaling in Synapse Elimination. Annu. Rev. Neurosci. 42, 107–127.

99. Wu, J.S., and Luo, L. (2006). A protocol for dissecting Drosophila melanogaster brains for live imaging or immunostaining. Nat. Protoc. 1, 2110–2115.

100. Wu, H., Xiong, W.C., and Mei, L. (2010). To build a synapse: Signaling pathways in neuromuscular junction assembly. Development 137, 1017–1033.

101. Yaksi, E., and Wilson, R.I. (2010). Electrical Coupling between Olfactory Glomeruli. Neuron 67, 1034–1047.

102. Zeng, H., and Sanes, J.R. (2017). Neuronal cell-type classification: Challenges, opportunities and the path forward. Nat. Rev. Neurosci. 18, 530–546.

