## Supplemental Tables for "Different olfactory neuron classes use distinct temporal and molecular programs to complete synaptic development"

*Genotypes*

Table 1. Genotypes for each Figure Panel

| Figure | Panel | Genotype |
| --- | --- | --- |
| **1** | **C-G** | *+; +; +; +* |
| **2, 2-S1** | **B-K** | *w; Or47b-GAL4 / UAS-Brp-Short-mStraw, UAS-mCD8-GFP; +; +* |
| **3, 3-S1** | **B-H** | *w; UAS-Brp-Short-mStraw, UAS-mCD8-GFP / +; Or67d-GAL4 / +; +* |
| **4, 4-S1** | **B-M** | *w; Mz19-GAL4 / UAS-Brp-Short-mStraw, UAS-mCD8-GFP; +; +* |
| **5, 5-S1** | **B-M** | *w; UAS-Brp-Short-mStraw, UAS-mCD8-GFP / +; NP3056-GAL4 / +; +* |
| **6** | **A** | *w; UAS-Brp-Short-mStraw / UAS-TNT-Imp; Or67d-GAL4 / +; +* |
|  | **B** | *w; UAS-Brp-Short-mStraw / UAS-TNT; Or67d-GAL4 / +; +* |
|  | **C** | *w; UAS-mCD8-GFP / UAS-TNT-Imp; Or67d-GAL4 / +; +* |
|  | **D** | *w; UAS-mCD8-GFP / UAS-TNT; Or67d-GAL4 / +; +* |
|  | **G, I** | *w; Mz19-GAL4, UAS-3xHA-mtdTomato / UAS-TNT-Imp; UAS-Brp-Short-GFP / +; +* |
|  | **H, J** | *w; Mz19-GAL4, UAS-3xHA-mtdTomato / UAS-TNT; UAS-Brp-Short-GFP / +; +* |
|  | **M** | *w; UAS-Brp-Short-mStraw / UAS-TNT-Imp; NP3056-GAL4 / +; +* |
|  | **N** | *w; UAS-Brp-Short-mStraw / UAS-TNT; NP3056-GAL4 / +; +* |
|  | **O** | *w; UAS-mCD8-GFP / UAS-TNT-Imp; NP3056-GAL4 / +; +* |
|  | **P** | *w; UAS-mCD8-GFP / UAS-TNT; NP3056-GAL4 / +; +* |
| **7** | **A** | *w; UAS-Brp-Short-mStraw / +; Or67d-GAL4 / +; +* |
|  | **B** | *w; UAS-Brp-Short-mStraw / UAS-NaChBac; Or67d-GAL4 / +; +* |
|  | **C** | *w; UAS-mCD8-GFP / +; Or67d-GAL4 / +; +* |
|  | **D** | *w; UAS-mCD8-GFP / UAS-NaChBac; Or67d-GAL4 / +; +* |
|  | **G, I** | *w; Mz19-GAL4, UAS-3xHA-mtdTomato / +; UAS-Brp-Short-GFP / +; +* |
|  | **H, J** | *w; Mz19-GAL4, UAS-3xHA-mtdTomato / UAS-NaChBac; UAS-Brp-Short-GFP / +; +* |
|  | **M** | *w; UAS-Brp-Short-mStraw / +; NP3056-GAL4 / +; +* |
|  | **N** | *w; UAS-Brp-Short-mStraw / UAS-NaChBac; NP3056-GAL4 / +; +* |
|  | **O** | *w; UAS-mCD8-GFP / +; NP3056-GAL4 / +; +* |
|  | **P** | *w; UAS-mCD8-GFP / UAS-NaChBac; NP3056-GAL4 / +; +* |
| **8** | **A** | *w; UAS-Brp-Short-mStraw / +; Or67d-GAL4 / +; +* |
|  | **B** | *w; UAS-Brp-Short-mStraw / UAS-Sgg-CA; Or67d-GAL4 / +; +* |
|  | **C** | *w; UAS-mCD8-GFP / +; Or67d-GAL4 / +; +* |
|  | **D** | *w; UAS-mCD8-GFP / UAS-Sgg-CA; Or67d-GAL4 / +; +* |
|  | **G** | *w; UAS-Brp-Short-mStraw / +; NP3056-GAL4 / +; +* |
|  | **H** | *w; UAS-Brp-Short-mStraw / UAS-Sgg-CA; NP3056-GAL4 / +; +* |
|  | **I** | *w; UAS-mCD8-GFP / +; NP3056-GAL4 / +; +* |
|  | **J** | *w; UAS-mCD8-GFP / UAS-Sgg-CA; NP3056-GAL4 / +; +* |
| **F3-FS2** | **B-M** | *w; AM29-GAL4 / UAS-Brp-Short-mStraw, UAS-mCD8-GFP; +; +* |
| **F3-FS3** | **B-M** | *w; AM29-GAL4 / UAS-Brp-Short-mStraw, UAS-mCD8-GFP; +; +* |
| **F6-FS1** | **A** | *w; UAS-Brp-Short-mStraw / UAS-TNT-Imp; Or67d-GAL4 / +; +* |
|  | **B** | *w; UAS-Brp-Short-mStraw / UAS-TNT; Or67d-GAL4 / +; +* |
|  | **C** | *w; UAS-mCD8-GFP / UAS-TNT-Imp; Or67d-GAL4 / +; +* |
|  | **D** | *w; UAS-mCD8-GFP / UAS-TNT; Or67d-GAL4 / +; +* |
|  | **E, G** | *w; Mz19-GAL4, UAS-3xHA-mtdTomato / UAS-TNT-Imp; UAS-Brp-Short-GFP / +; +* |
|  | **F, H** | *w; Mz19-GAL4, UAS-3xHA-mtdTomato / UAS-TNT; UAS-Brp-Short-GFP / +; +* |
|  | **I** | *w; UAS-Brp-Short-mStraw / UAS-TNT-Imp; NP3056-GAL4 / +; +* |
|  | **J** | *w; UAS-Brp-Short-mStraw / UAS-TNT; NP3056-GAL4 / +; +* |
|  | **K** | *w; UAS-mCD8-GFP / UAS-TNT-Imp; NP3056-GAL4 / +; +* |
|  | **L** | *w; UAS-mCD8-GFP / UAS-TNT; NP3056-GAL4 / +; +* |
| **F7-FS1** | **A** | *w; UAS-Brp-Short-mStraw / +; Or67d-GAL4 / +; +* |
|  | **B** | *w; UAS-Brp-Short-mStraw / UAS-NaChBac; Or67d-GAL4 / +; +* |
|  | **C** | *w; UAS-mCD8-GFP / +; Or67d-GAL4 / +; +* |
|  | **D** | *w; UAS-mCD8-GFP / UAS-NaChBac; Or67d-GAL4 / +; +* |
|  | **E, G** | *w; Mz19-GAL4, UAS-3xHA-mtdTomato / +; UAS-Brp-Short-GFP / +; +* |
|  | **F, H** | *w; Mz19-GAL4, UAS-3xHA-mtdTomato / UAS-NaChBac; UAS-Brp-Short-GFP / +; +* |
|  | **I** | *w; UAS-Brp-Short-mStraw / +; NP3056-GAL4 / +; +* |
|  | **J** | *w; UAS-Brp-Short-mStraw / UAS-NaChBac; NP3056-GAL4 / +; +* |
|  | **K** | *w; UAS-mCD8-GFP / +; NP3056-GAL4 / +; +* |
|  | **L** | *w; UAS-mCD8-GFP / UAS-NaChBac; NP3056-GAL4 / +; +* |
| **F8-FS1** | **A** | *w; UAS-Brp-Short-mStraw / +; Or67d-GAL4 / +; +* |
|  | **B** | *w; UAS-Brp-Short-mStraw / UAS-Sgg-DN; Or67d-GAL4 / +; +* |
|  | **C** | *w; UAS-mCD8-GFP / +; Or67d-GAL4 / +; +* |
|  | **D** | *w; UAS-mCD8-GFP / UAS-Sgg-DN; Or67d-GAL4 / +; +* |
|  | **G, I** | *w; Mz19-GAL4, UAS-3xHA-mtdTomato / +; UAS-Brp-Short-GFP / +; +* |
|  | **H, J** | *w; Mz19-GAL4, UAS-3xHA-mtdTomato / UAS-Sgg-DN; UAS-Brp-Short-GFP / +; +* |
|  | **M** | *w; UAS-Brp-Short-mStraw / +; NP3056-GAL4 / +; +* |
|  | **N** | *w; UAS-Brp-Short-mStraw / UAS-Sgg-DN; NP3056-GAL4 / +; +* |
|  | **O** | *w; UAS-mCD8-GFP / +; NP3056-GAL4 / +; +* |
|  | **P** | *w; UAS-mCD8-GFP / UAS-Sgg-DN; NP3056-GAL4 / +; +* |
| **F8-FS2** | **A** | *w; UAS-Brp-Short-mStraw / +; Or67d-GAL4 / +; +* |
|  | **B** | *w; UAS-Brp-Short-mStraw / UAS-Sgg-DN; Or67d-GAL4 / +; +* |
|  | **C** | *w; UAS-Brp-Short-mStraw / UAS-Sgg-CA; Or67d-GAL4 / +; +* |
|  | **D** | *w; UAS-mCD8-GFP / +; Or67d-GAL4 / +; +* |
|  | **E** | *w; UAS-mCD8-GFP / UAS-Sgg-DN; Or67d-GAL4 / +; +* |
|  | **F** | *w; UAS-mCD8-GFP / UAS-Sgg-CA; Or67d-GAL4 / +; +* |
|  | **G, I** | *w; Mz19-GAL4, UAS-3xHA-mtdTomato / +; UAS-Brp-Short-GFP / +; +* |
|  | **H, J** | *w; Mz19-GAL4, UAS-3xHA-mtdTomato / UAS-Sgg-DN; UAS-Brp-Short-GFP / +; +* |
|  | **K** | *w; UAS-Brp-Short-mStraw / +; NP3056-GAL4 / +; +* |
|  | **L** | *w; UAS-Brp-Short-mStraw / UAS-Sgg-DN; NP3056-GAL4 / +; +* |
|  | **M** | *w; UAS-Brp-Short-mStraw / UAS-Sgg-CA; NP3056-GAL4 / +; +* |
|  | **N** | *w; UAS-mCD8-GFP / +; NP3056-GAL4 / +; +* |
|  | **O** | *w; UAS-mCD8-GFP / UAS-Sgg-DN; NP3056-GAL4 / +; +* |
|  | **P** | *w; UAS-mCD8-GFP / UAS-Sgg-CA; NP3056-GAL4 / +; +* |

| VA1lm  ORN Male | Brp Mean +/- SEM (*n*)  GFP Mean +/- SEM (*n*) | | | | vs. 48h | vs. 60h | vs. 72h | vs. 84h | vs. 92h |
| --- | --- | --- | --- | --- | --- | --- | --- | --- | --- |
| 48 h | N/A  N/A | | | | N/A | N/A | N/A | N/A | N/A |
|  |  |  |  |  | N/A | N/A | N/A | N/A | N/A |
| 60 h | 692.1 | +/- | 37.92 | *(14)* | N/A | *---* | *.002* | *<.001.* | *<.001* |
|  | 595.7 | +/- | 50.1 | *(14)* | N/A | *---* | *<.001* | *<.001* | *<.001* |
| 72 h | 891.6 | +/- | 85.03 | *(8)* | N/A | *.002* | *---* | *<.001* | *<.001* |
|  | 1086 | +/- | 124.5 | *(8)* | N/A | *<.001* | *---* | *<.001* | *<.001* |
| 84 h | 1279 | +/- | 41.43 | *(12)* | N/A | *<.001* | *<.001* | *---* | *.063* |
|  | 1540 | +/- | 44.7 | *(12)* | N/A | *<.001* | *<.001* | *---* | *<.001* |
| 92 h | 1422 | +/- | 25.28 | *(18)* | N/A | *<.001* | *<.001* | *.063* | *---* |
|  | 1979 | +/- | 52.75 | *(18)* | N/A | *<.001* | *<.001* | *<.001* | *---* |
| VA1lm  ORN Male | Brp Mean +/- SEM (*n*)  GFP Mean +/- SEM (*n*) | | | | vs. 0d | vs. 5d | vs. 10d | vs. 15d | vs. 20d |
| 0 d | 1576 | +/- | 23.89 | *(14)* | *---* | *<.001* | *.002* | *.001* | *<.001* |
|  | 2425 | +/- | 78.98 | *(14)* | *---* | *.002* | *<.001* | *<.001* | *<.001* |
| 5 d | 2144 | +/- | 36.11 | *(12)* | *<.001* | *---* | *.027* | *.237* | *.631* |
|  | 3130 | +/- | 67.75 | *(12)* | *.002* | *---* | *.119* | *.005* | *.003* |
| 10 d | 1891 | +/- | 49.21 | *(21)* | *.002* | *.027* | *---* | *.958* | *.685* |
|  | 3527 | +/- | 66.58 | *(21)* | *<.001* | *.119* | *---* | *.476* | *.298* |
| 15 d | 1948 | +/- | 47.68 | *(12)* | *.001* | *.237* | *.958* | *---* | *.975* |
|  | 3794 | +/- | 111.6 | *(12)* | *<.001* | *.005* | *.476* | *---* | *.996* |
| 20 d | 2007 | +/- | 140.2 | *(10)* | *<.001* | *.631* | *.685* | *.975* | *---* |
|  | 3865 | +/- | 292.9 | *(10)* | *<.001* | *.003* | *.298* | *.996* | *---* |

**Figure 2 – table supplement 1. Statistical data and comparisons between each time-point for male VA1lm ORNs.**

Table of average Bruchpilot-Short-mStrawberry puncta counts (Brp Mean, red) and average membrane GFP neurite volumes (GFP Mean, green) for each time-point from male VA1lm ORN time-courses. Mean values are shown with calculated standard error of the mean (SEM) and number of glomeruli analyzed (*n*). Each time-point is compared to each other time-point using one-way ANOVA tests followed by Tukey’s tests to correct for multiple comparisons in determining *p*-values. *p*-values for Brp-Short-mStrawberry puncta are in red and in green for membrane GFP neurite volume.

| VA1lm ORN Female | Brp Mean +/- SEM (*n*)  GFP Mean +/- SEM (*n*) | | | | vs. 48h | vs. 60h | vs. 72h | vs. 84h | vs. 92h |
| --- | --- | --- | --- | --- | --- | --- | --- | --- | --- |
| 48 h | N/A  N/A | | | | N/A | N/A | N/A | N/A | N/A |
|  |  |  |  |  | N/A | N/A | N/A | N/A | N/A |
| 60 h | 413.7 | +/- | 22.39 | *(12)* | N/A | *---* | *<.001* | *<.001* | *<.001* |
|  | 240.0 | +/- | 29.86 | *(12)* | N/A | *---* | *<.001* | *<.001* | *<.001* |
| 72 h | 678.0 | +/- | 39.61 | *(12)* | N/A | *<.001* | *---* | *<.001* | *<.001* |
|  | 1007 | +/- | 78.05 | *(12)* | N/A | *<.001* | *---* | *.376* | *<.001* |
| 84 h | 890.9 | +/- | 19.89 | *(16)* | N/A | *<.001* | *<.001* | *---* | *.089* |
|  | 1124 | +/- | 41.04 | *(16)* | N/A | *<.001* | *.376* | *---* | *<.001* |
| 92 h | 970.8 | +/- | 21.71 | *(19)* | N/A | *<.001* | *<.001* | *.089* | *---* |
|  | 1392 | +/- | 43.31 | *(19)* | N/A | *<.001* | *<.001* | *<.001* | *---* |
| VA1lm ORN Female | Brp Mean +/- SEM (*n*)  GFP Mean +/- SEM (*n*) | | | | vs. 0d | vs. 5d | vs. 10d | vs. 15d | vs. 20d |
| 0 d | 1151 | +/- | 29.03 | *(12)* | *---* | *<.001* | *<.001* | *<.001* | *.001* |
|  | 1655 | +/- | 79.87 | *(12)* | *---* | *<.001* | *<.001* | *<.001* | *<.001* |
| 5 d | 1588 | +/- | 37.47 | *(10)* | *<.001* | *---* | *<.001* | *.013* | *.001* |
|  | 2302 | +/- | 70.04 | *(10)* | *<.001* | *---* | *.782* | *.257* | *.825* |
| 10 d | 1364 | +/- | 35.08 | *(24)* | *<.001* | *<.001* | *---* | *.962.* | *>.999* |
|  | 2422 | +/- | 63.00 | *(24)* | *<.001* | *.782* | *---* | *.701* | *>.999* |
| 15 d | 1396 | +/- | 28.84 | *(10)* | *<.001* | *.013* | *.962* | *---* | *.972* |
|  | 2557 | +/- | 64.14 | *(10)* | *<.001* | *.257* | *.701* | *---* | *.819* |
| 20 d | 1362 | +/- | 20.98 | *(12)* | *.001* | *.001* | *>.999* | *.972* | *---* |
|  | 2429 | +/- | 88.59 | *(12)* | *<.001* | *.825* | *>.999* | *.819* | *---* |

**Figure 2 – table supplement 2. Statistical data and comparisons between each time-point for female VA1lm ORNs.**

Table of average Bruchpilot-Short-mStrawberry puncta counts (Brp Mean, red) and average membrane GFP neurite volumes (GFP Mean, green) for each time-point from female VA1lm ORN time-courses. Mean values are shown with calculated standard error of the mean (SEM) and number of glomeruli analyzed (*n*). Each time-point is compared to each other time-point using one-way ANOVA tests followed by Tukey’s tests to correct for multiple comparisons in determining *p*-values. *p*-values for Brp-Short-mStrawberry puncta are in red and in green for membrane GFP neurite volume.

| DA1 ORN Male | Brp Mean +/- SEM (*n*)  GFP Mean +/- SEM (*n*) | | | | vs. 0d | vs. 3d | vs. 6d | vs. 9d | vs. 12d | vs. 15d | vs. 18d |
| --- | --- | --- | --- | --- | --- | --- | --- | --- | --- | --- | --- |
| 0 d | 1251 | +/- | 36.03 | *(15)* | *---* | *<.001* | *<.001* | *<.001* | *<.001* | *<.001* | *.148* |
|  | 2163 | +/- | 205.6 | *(14)* | *---* | *.13* | *<.001* | *<.001* | *<.001* | *<.001* | *<.001* |
| 3 d | 1662 | +/- | 32.17 | *(20)* | *<.001* | *---* | *.003* | *.006* | *.002* | *<.001* | *.555* |
|  | 2949 | +/- | 126.2 | *(20)* | *.13* | *---* | *.004* | *.449* | *<.001* | *.085* | *.296* |
| 6 d | 2007 | +/- | 59.2 | *(18)* | *<.001* | *.003* | *---* | *>.999* | *>.999* | *.998* | *<.001* |
|  | 4026 | +/- | 175 | *(18)* | *<.001* | *.004* | *---* | *.45* | *.996* | *.931* | *.79* |
| 9 d | 1978 | +/- | 82.22 | *(20)* | *<.001* | *.006* | *>.999* | *---* | *>.999* | *.969* | *<.001* |
|  | 3481 | +/- | 208.8 | *(20)* | *<.001* | *.449* | *.45* | *---* | *.159* | *.977* | *>.999* |
| 12 d | 2017 | +/- | 92.57 | *(18)* | *<.001* | *.002* | *>.999* | *>.999* | *---* | *>.999* | *<.001* |
|  | 4211 | +/- | 197.2 | *(16)* | *<.001* | *<.001* | *.996.* | *.159* | *---* | *.631* | *.437* |
| 15 d | 2058 | +/- | 53.87 | *(19)* | *<.001* | *<.001* | *.998* | *.969* | *>.999* | *---* | *<.001* |
|  | 3720 | +/- | 164.9 | *(19)* | *<.001* | *.085* | *.931* | *.977* | *.631* | *---* | *>.999* |
| 18 d | 1499 | +/- | 55.69 | *(16)* | *.148* | *.555* | *<.001* | *<.001* | *<.001* | *<.001* | *---* |
|  | 3600 | +/- | 358.3 | *(15)* | *<.001* | *.296* | *.79* | *>.999* | *.437* | *>.999* | *---* |

**Figure 3 – table supplement 1. Statistical data and comparisons between each time-point for male DA1 ORNs.**

Table of average Bruchpilot-Short-mStrawberry puncta counts (Brp Mean, red) and average membrane GFP neurite volumes (GFP Mean, green) for each time-point from male DA1 ORN time-courses. Mean values are shown with calculated standard error of the mean (SEM) and number of glomeruli analyzed (*n*). Each time-point is compared to each other time-point using one-way ANOVA tests followed by Tukey’s tests to correct for multiple comparisons in determining *p*-values. *p*-values for Brp-Short-mStrawberry puncta are in red and in green for membrane GFP neurite volume.

| DA1 ORN Female | Brp Mean +/- SEM (*n*)  GFP Mean +/- SEM (*n*) | | | | vs. 0d | vs. 3d | vs. 6d | vs. 9d | vs. 12d | vs. 15d | vs. 18d |
| --- | --- | --- | --- | --- | --- | --- | --- | --- | --- | --- | --- |
| 0 d | 898.5 | +/- | 26.31 | *(11)* | *---* | *.002* | *<.001* | *<.001* | *<.001* | *<.001* | *.033* |
|  | 1058 | +/- | 105 | *(11)* | *---* | *.03* | *.529* | *.365* | *<.001* | *.036* | *.027* |
| 3 d | 1080 | +/- | 29.12 | *(16)* | *.002* | *---* | *.912* | *.075* | *.013* | *.64* | *.979* |
|  | 1540 | +/- | 77.81 | *(16)* | *.03* | *---* | *.829* | *.843* | *.912* | *>.999* | *>.999* |
| 6 d | 1130 | +/- | 26.55 | *(13)* | *<.001* | *.912* | *---* | *.734* | *.345* | *>.999* | *.461* |
|  | 1347 | +/- | 75.61 | *(13)* | *.529* | *.829* | *---* | *>.999* | *.191* | *.862* | *.798* |
| 9 d | 1194 | +/- | 24.35 | *(18)* | *<.001* | *.075* | *.734* | *---* | *.993* | *.915* | *.007* |
|  | 1366 | +/- | 73.68 | *(18)* | *.365* | *.843* | *>.999* | *---* | *.165* | *.878* | *.812* |
| 12 d | 1221 | +/- | 21.8 | *(17)* | *<.001* | *.013* | *.345* | *.993* | *---* | *.561* | *<.001* |
|  | 1693 | +/- | 69.63 | *(17)* | *<.001* | *.912* | *.191* | *.165* | *---* | *.884* | *.943* |
| 15 d | 1148 | +/- | 36.94 | *(16)* | *<.001* | *.64* | *>.999* | *.915* | *.561* | *---* | *.174* |
|  | 1530 | +/- | 125.2 | *(16)* | *.036* | *>.999* | *.862* | *.878* | *.884* | *---* | *>.999* |
| 18 d | 1044 | +/- | 38.46 | *(15)* | *.033* | *.979* | *.461* | *.007* | *<.001* | *.174* | *---* |
|  | 1551 | +/- | 143.3 | *(15)* | *.027* | *>.999* | *.798* | *.812* | *.943* | *>.999* | *---* |

**Figure 3 – table supplement 2. Statistical data and comparisons between each time-point for female DA1 ORNs.**

Table of average Bruchpilot-Short-mStrawberry puncta counts (Brp Mean, red) and average membrane GFP neurite volumes (GFP Mean, green) for each time-point from female DA1 ORN time-courses. Mean values are shown with calculated standard error of the mean (SEM) and number of glomeruli analyzed (*n*). Each time-point is compared to each other time-point using one-way ANOVA tests followed by Tukey’s tests to correct for multiple comparisons in determining *p*-values. *p*-values for Brp-Short-mStrawberry puncta are in red and in green for membrane GFP neurite volume.

| DL4 ORN Male | Brp Mean +/- SEM (*n*)  GFP Mean +/- SEM (*n*) | | | | | | | vs. 48h | | vs. 60h | | vs. 72h | | | vs. 84h | | vs. 92h | |
| --- | --- | --- | --- | --- | --- | --- | --- | --- | --- | --- | --- | --- | --- | --- | --- | --- | --- | --- |
| 48 h | 98.58 | +/- | | | 5.109 | *(12)* | | *---* | | *.002* | | *<.001* | | | *<.001* | | *<.001* | |
|  | 217.9 | +/- | | | 12.14 | *(12)* | | *---* | | *.824* | | *<.001* | | | *<.001* | | *<.001* | |
| 60 h | 133.3 | +/- | | | 5.189 | *(16)* | | *.002* | | *---* | | *.1* | | | *.274* | | *.003* | |
|  | 238.7 | +/- | | | 9.555 | *(16)* | | *.824* | | *---* | | *.011* | | | *<.001* | | *<.001* | |
| 72 h | 154.1 | +/- | | | 6.807 | *(16)* | | *<.001* | | *.1* | | *---* | | | *.987* | | *.601* | |
|  | 299.3 | +/- | | | 17.91 | *(16)* | | *<.001* | | *.011* | | *---* | | | *.894* | | *.318* | |
| 84 h | 149.9 | +/- | | | 7.720 | *(16)* | | *<.001* | | *.274* | | *.987* | | | *---* | | *.315* | |
|  | 315.6 | +/- | | | 12.76 | *(16)* | | *<.001* | | *<.001* | | *.894* | | | *---* | | *.826* | |
| 92 h | 166.8 | +/- | | | 3.616 | *(13)* | | *<.001* | | *.003* | | *.601* | | | *.315* | | *---* | |
|  | 335.8 | +/- | | | 11.39 | *(13)* | | *<.001* | | *<.001* | | *.318* | | | *.826* | | *---* | |
| DL4 ORN Male | Brp Mean +/- SEM (*n*)  GFP Mean +/- SEM (*n*) | | | | | | | vs. 0d | vs. 3d | | vs. 6d | | vs. 9d | vs. 12d | | vs. 15d | | vs. 18d |
| 0 d | 226 | | +/- | 8.897 | | | *(14)* | *---* | *.03* | | *.293* | | *.999* | *.003* | | *.066* | | *.405* |
|  | 389.6 | | +/- | 19.08 | | | *(14)* | *---* | *.998* | | *.998* | | *.029* | *.135* | | *<.001* | | *<.001* |
| 3 d | 272.5 | | +/- | 7.655 | | | *(12)* | *.03* | *---* | | *.873* | | *.136* | *.997* | | *.999* | | *.738* |
|  | 374 | | +/- | 19.17 | | | *(12)* | *.998* | *---* | | *>.999* | | *.146* | *.454* | | *<.001* | | *<.001* |
| 6 d | 255.3 | | +/- | 7.732 | | | *(18)* | *.293* | *.873* | | *---* | | *.68* | *.448* | | *.985* | | *>.999* |
|  | 375.4 | | +/- | 16.88 | | | *(18)* | *.998* | *>.999* | | *---* | | *.074* | *.301* | | *<.001* | | *<.001* |
| 9 d | 233.2 | | +/- | 12.07 | | | *(12)* | *.999* | *.136* | | *.68* | | *---* | *.023* | | *.266* | | *.799* |
|  | 297.8 | | +/- | 26.61 | | | *(11)* | *.029* | *.146* | | *.074* | | *---* | *.988* | | *.202* | | *.116* |
| 12 d | 281.1 | | +/- | 12.21 | | | *(14)* | *.003* | *.997* | | *.448* | | *.023* | *---* | | *.906* | | *.28* |
|  | 319.6 | | +/- | 20.66 | | | *(14)* | *.135* | *.454* | | *.301* | | *.988* | *---* | | *.016* | | *.005* |
| 15 d | 265.3 | | +/- | 9.27 | | | *(16)* | *.066* | *.999* | | *.985* | | *.266* | *.906* | | *---* | | *.937* |
|  | 229.7 | | +/- | 18.03 | | | *(15)* | *<.001* | *<.001* | | *<.001* | | *.202* | *.016* | | *---* | | *>.999* |
| 18 d | 252.1 | | +/- | 8.575 | | | *(20)* | *.405* | *.738* | | *>.999* | | *.799* | *.28* | | *.937* | | *---* |
|  | 226.6 | | +/- | 12.81 | | | *(20)* | *<.001* | *<.001* | | *<.001* | | *.116* | *.005* | | *>.999* | | *---* |

**Figure 3 – table supplement 3. Statistical data and comparisons between each time-point for male DL4 ORNs.**

Table of average Bruchpilot-Short-mStrawberry puncta counts (Brp Mean, red) and average membrane GFP neurite volumes (GFP Mean, green) for each time-point from male DL4 ORN time-courses. Mean values are shown with calculated standard error of the mean (SEM) and number of glomeruli analyzed (*n*). Each time-point is compared to each other time-point using one-way ANOVA tests followed by Tukey’s tests to correct for multiple comparisons in determining *p*-values. *p*-values for Brp-Short-mStrawberry puncta are in red and in green for membrane GFP neurite volume.

| DL4 ORN Female | Brp Mean +/- SEM (*n*)  GFP Mean +/- SEM (*n*) | | | | | vs. 48h | | vs. 60h | | vs. 72h | | | vs. 84h | | vs. 92h | |
| --- | --- | --- | --- | --- | --- | --- | --- | --- | --- | --- | --- | --- | --- | --- | --- | --- |
| 48 h | 103.6 | +/- | | 5.577 | *(19)* | *---* | | *<.001* | | *<.001* | | | *<.001* | | *<.001* | |
|  | 197.4 | +/- | | 10.23 | *(19)* | *---* | | *<.001* | | *<.001* | | | *<.001* | | *<.001* | |
| 60 h | 153.2 | +/- | | 6.666 | *(16)* | *<.001* | | *---* | | *.053* | | | *.047* | | *.02* | |
|  | 288.8 | +/- | | 8.641 | *(16)* | *<.001* | | *---* | | *>.999* | | | *.499* | | *.523* | |
| 72 h | 179.4 | +/- | | 10.48 | *(14)* | *<.001* | | *.053* | | *---* | | | *>.999* | | *.997* | |
|  | 285.4 | +/- | | 13.22 | *(14)* | *<.001* | | *>.999* | | *---* | | | *.389* | | *.417* | |
| 84 h | 177.7 | +/- | | 3.503 | *(20)* | *<.001* | | *.047* | | *>.999* | | | *---* | | *.978* | |
|  | 311.1 | +/- | | 8.508 | *(20)* | *<.001* | | *.499* | | *.389* | | | *---* | | *>.999* | |
| 92 h | 182.9 | +/- | | 5.836 | *(14)* | *<.001* | | *.02* | | *.997* | | | *.978* | | *---* | |
|  | 312.5 | +/- | | 10.48 | *(14)* | *<.001* | | *.523* | | *.417* | | | *>.999* | | *---* | |
| DL4 ORN Female | Brp Mean +/- SEM (*n*)  GFP Mean +/- SEM (*n*) | | | | | vs. 0d | vs. 3d | | vs. 6d | | vs. 9d | vs. 12d | | vs. 15d | | vs. 18d |
| 0 d | 227 | | +/- | 11.81 | *(19)* | *---* | *.264* | | *<.001* | | *.185* | *.795* | | *.003* | | *.023* |
|  | 399.1 | | +/- | 32.37 | *(19)* | *---* | *>.999* | | *.994* | | *.195* | *.289* | | *.165* | | *.204* |
| 3 d | 268.3 | | +/- | 9.725 | *(16)* | *.264* | *---* | | *.541* | | *>.999* | *.981* | | *.602* | | *.954* |
|  | 383.4 | | +/- | 27.26 | *(16)* | *>.999* | *---* | | *.928* | | *.513* | *.619* | | *.401* | | *.497* |
| 6 d | 300.8 | | +/- | 10.12 | *(20)* | *<.001* | *.541* | | *---* | | *.499* | *.11* | | *>.999* | | *.99* |
|  | 420.3 | | +/- | 15.69 | *(20)* | *.994* | *.928* | | *---* | | *.035* | *.068* | | *.038* | | *.043* |
| 9 d | 269.1 | | +/- | 10.54 | *(20)* | *.185* | *>.999* | | *.499* | | *---* | *.969* | | *.574* | | *.952* |
|  | 321 | | +/- | 21.93 | *(20)* | *.195* | *.513* | | *.035* | | *---* | *>.999* | | *>.999* | | *>.999* |
| 12 d | 252.6 | | +/- | 9.024 | *(16)* | *.795* | *.981* | | *.11* | | *.969* | *---* | | *.165* | | *.543* |
|  | 323.3 | | +/- | 21.61 | *(16)* | *.289* | *.619* | | *.068* | | *>.999* | *---* | | *.999* | | *>.999* |
| 15 d | 302.6 | | +/- | 28.3 | *(13)* | *.003* | *.602* | | *>.999* | | *.574* | *.165* | | *---* | | *.989* |
|  | 303.7 | | +/- | 29.41 | *(11)* | *.165* | *.401* | | *.038* | | *>.999* | *.999* | | *---* | | *>.999* |
| 18 d | 287.3 | | +/- | 10.51 | *(15)* | *.023* | *.954* | | *.99* | | *.952* | *.543* | | *.989* | | *---* |
|  | 315.7 | | +/- | 20.02 | *(15)* | *.204* | *.497* | | *.043* | | *>.999* | *>.999* | | *>.999* | | *---* |

**Figure 3 – table supplement 4. Statistical data and comparisons between each time-point for female DL4 ORNs.**

Table of average Bruchpilot-Short-mStrawberry puncta counts (Brp Mean, red) and average membrane GFP neurite volumes (GFP Mean, green) for each time-point from female DL4 ORN time-courses. Mean values are shown with calculated standard error of the mean (SEM) and number of glomeruli analyzed (*n*). Each time-point is compared to each other time-point using one-way ANOVA tests followed by Tukey’s tests to correct for multiple comparisons in determining *p*-values. *p*-values for Brp-Short-mStrawberry puncta are in red and in green for membrane GFP neurite volume.

| DA1 PN Male | Brp Mean +/- SEM (*n*)  GFP Mean +/- SEM (*n*) | | | | | | | vs. 48h | | vs. 60h | | vs. 72h | | | vs. 84h | | vs. 92h | |
| --- | --- | --- | --- | --- | --- | --- | --- | --- | --- | --- | --- | --- | --- | --- | --- | --- | --- | --- |
| 48 h | 298.1 | +/- | | | 17.98 | *(14)* | | *---* | | *<.001* | | *<.001* | | | *<.001* | | *<.001* | |
|  | 377.1 | +/- | | | 18.04 | *(14)* | | *---* | | *.009* | | *<.001* | | | *<.001* | | *<.001* | |
| 60 h | 450.5 | +/- | | | 25.09 | *(10)* | | *<.001* | | *---* | | *<.001* | | | *<.001* | | *<.001* | |
|  | 542.6 | +/- | | | 38.45 | *(10)* | | *.009* | | *---* | | *.254* | | | *<.001* | | *<.001* | |
| 72 h | 627.8 | +/- | | | 11.23 | *(12)* | | *<.001* | | *<.001* | | *---* | | | *.239* | | *<.001* | |
|  | 644.8 | +/- | | | 53.83 | *(12)* | | *<.001* | | *.254* | | *---* | | | *.278* | | *.002* | |
| 84 h | 678.5 | +/- | | | 13.22 | *(22)* | | *<.001* | | *<.001* | | *.239* | | | *---* | | *<.001* | |
|  | 728.3 | +/- | | | 15.25 | *(22)* | | *<.001* | | *<.001* | | *.278* | | | *---* | | *.156* | |
| 92 h | 828.5 | +/- | | | 20.79 | *(16)* | | *<.001* | | *<.001* | | *<.001* | | | *<.001* | | *---* | |
|  | 816.4 | +/- | | | 32.51 | *(16)* | | *<.001* | | *<.001* | | *.002* | | | *.156* | | *---* | |
| DA1 PN Male | Brp Mean +/- SEM (*n*)  GFP Mean +/- SEM (*n*) | | | | | | | vs. 0d | vs. 3d | | vs. 6d | | vs. 9d | vs. 12d | | vs. 15d | | vs. 18d |
| 0 d | 751.1 | | +/- | 25.33 | | | *(22)* | *---* | *<.001* | | *<.001* | | *<.001* | *<.001* | | *<.001* | | *<.001* |
|  | 742.9 | | +/- | 29.6 | | | *(22)* | *---* | *<.001* | | *<.001* | | *<.001* | *<.001* | | *<.001* | | *<.001* |
| 3 d | 1055 | | +/- | 36.21 | | | *(20)* | *<.001* | *---* | | *<.001* | | *.066* | *.036* | | *.004* | | *.002* |
|  | 1192 | | +/- | 63.97 | | | *(20)* | *<.001* | *---* | | *.261* | | *.898* | *.003* | | *.296* | | *.154* |
| 6 d | 1412 | | +/- | 56.84 | | | *(17)* | *<.001* | *<.001* | | *---* | | *.3* | *.187* | | *.852* | | *.76* |
|  | 1411 | | +/- | 52.48 | | | *17)* | *<.001* | *.261* | | *---* | | *.972* | *.733* | | *>.999* | | *>.999* |
| 9 d | 1255 | | +/- | 59.11 | | | *(13)* | *<.001* | *.066* | | *.3* | | *---* | *>.999* | | *.978* | | *.981* |
|  | 1315 | | +/- | 76.72 | | | *(13)* | *<.001* | *.898* | | *.972* | | *---* | *.24* | | *.971* | | *.924* |
| 12 d | 1252 | | +/- | 51.81 | | | *(18)* | *<.001* | *.036* | | *.187* | | *>.999* | *---* | | *.96* | | *.962* |
|  | 1559 | | +/- | 74.61 | | | *(19)* | *<.001* | *.003* | | *.733* | | *.24* | *---* | | *.807* | | *.841* |
| 15 d | 1320 | | +/- | 52.11 | | | *(13)* | *<.001* | *.004* | | *.852* | | *.978* | *.96* | | *---* | | *>.999* |
|  | 1416 | | +/- | 55.53 | | | *(14)* | *<.001* | *.296* | | *>.999* | | *.971* | *.807* | | *---* | | *>.999* |
| 18 d | 1315 | | +/- | 53.53 | | | *(17)* | *<.001* | *.002* | | *.76* | | *.981* | *.962* | | *>.999* | | *---* |
|  | 1432 | | +/- | 110.7 | | | *(18)* | *<.001* | *.154* | | *>.999* | | *.924* | *.841* | | *>.999* | | *---* |

**Figure 4 – table supplement 1. Statistical data and comparisons between each time-point for male DA1 PNs.**

Table of average Bruchpilot-Short-mStrawberry puncta counts (Brp Mean, red) and average membrane GFP neurite volumes (GFP Mean, green) for each time-point from male DA1 PN time-courses. Mean values are shown with calculated standard error of the mean (SEM) and number of glomeruli analyzed (*n*). Each time-point is compared to each other time-point using one-way ANOVA tests followed by Tukey’s tests to correct for multiple comparisons in determining *p*-values. *p*-values for Brp-Short-mStrawberry puncta are in red and in green for membrane GFP neurite volume.

| DA1 PN Female | Brp Mean +/- SEM (*n*)  GFP Mean +/- SEM (*n*) | | | | | | | vs. 48h | | vs. 60h | | vs. 72h | | | vs. 84h | | vs. 92h | |
| --- | --- | --- | --- | --- | --- | --- | --- | --- | --- | --- | --- | --- | --- | --- | --- | --- | --- | --- |
| 48 h | 241 | +/- | | | 14.44 | *(12)* | | *---* | | *<.001* | | *<.001* | | | *<.001* | | *<.001* | |
|  | 314.4 | +/- | | | 14.95 | *(12)* | | *---* | | *.285* | | *<.001* | | | *<.001* | | *<.001* | |
| 60 h | 439.4 | +/- | | | 15.08 | *(14)* | | *<.001* | | *---* | | *.985* | | | *<.001* | | *.001* | |
|  | 388.5 | +/- | | | 29.24 | *(14)* | | *.285* | | *---* | | *.013* | | | *<.001* | | *<.001* | |
| 72 h | 450.9 | +/- | | | 12.25 | *(16)* | | *<.001* | | *.985* | | *---* | | | *<.001* | | *.005* | |
|  | 503 | +/- | | | 25.14 | *(16)* | | *<.001* | | *.013* | | *---* | | | *.008* | | *<.001* | |
| 84 h | 567.1 | +/- | | | 18.5 | *(16)* | | *<.001* | | *<.001* | | *<.001* | | | *---* | | *.393* | |
|  | 619.8 | +/- | | | 21.48 | *(16)* | | *<.001* | | *<.001* | | *.008* | | | *---* | | *.964* | |
| 92 h | 528.9 | +/- | | | 16.77 | *(16)* | | *<.001* | | *.001* | | *.005* | | | *.393* | | *---* | |
|  | 641.9 | +/- | | | 27.01 | *(16)* | | *<.001* | | *<.001* | | *<.001* | | | *.964* | | *---* | |
| DA1 PN Female | Brp Mean +/- SEM (*n*)  GFP Mean +/- SEM (*n*) | | | | | | | vs. 0d | vs. 3d | | vs. 6d | | vs. 9d | vs. 12d | | vs. 15d | | vs. 18d |
| 0 d | 524.6 | | +/- | 17.02 | | | *(17)* | *---* | *<.001* | | *<.001* | | *<.001* | *<.001* | | *<.001* | | *<.001* |
|  | 563.8 | | +/- | 39.14 | | | *(17)* | *---* | *.011* | | *.001* | | *.015* | *.048* | | *<.001* | | *<.001* |
| 3 d | 767 | | +/- | 23.94 | | | *(17)* | *<.001* | *---* | | *.002* | | *.044* | *<.001* | | *<.001* | | *<.001* |
|  | 825.8 | | +/- | 40.61 | | | *(17)* | *.011* | *---* | | *.997* | | *>.999* | *.999* | | *.496* | | *.828* |
| 6 d | 962 | | +/- | 36.94 | | | *(17)* | *<.001* | *.002* | | *---* | | *.951* | *.637* | | *>.999* | | *.875* |
|  | 870.8 | | +/- | 50.81 | | | *(17)* | *.001* | *.997* | | *---* | | *.987* | *.921* | | *.861* | | *.99* |
| 9 d | 913.4 | | +/- | 33.42 | | | *(18)* | *<.001* | *.044* | | *.951* | | *---* | *.107* | | *.85* | | *.267* |
|  | 813.9 | | +/- | 60.33 | | | *(18)* | *.015* | *>.999* | | *.987* | | *---* | *>.999* | | *.376* | | *.727* |
| 12 d | 1043 | | +/- | 46.36 | | | *(17)* | *<.001* | *<.001* | | *.637* | | *.107* | *---* | | *.81* | | *>.999* |
|  | 787.7 | | +/- | 37.04 | | | *(17)* | *.048* | *.999* | | *.921* | | *>.999* | *---* | | *.21* | | *.511* |
| 15 d | 976 | | +/- | 31.88 | | | *(17)* | *<.001* | *<.001* | | *>.999* | | *.85* | *.81* | | *---* | | *.962* |
|  | 965.7 | | +/- | 46.25 | | | *(17)* | *<.001* | *.496* | | *.861* | | *.376* | *.21* | | *---* | | *.998* |
| 18 d | 1023 | | +/- | 41.82 | | | *(17)* | *<.001* | *<.001* | | *.875* | | *.267* | *>.999* | | *.962* | | *---* |
|  | 925.9 | | +/- | 78.21 | | | *(17)* | *<.001* | *.828* | | *.99* | | *.727* | *.511* | | *.998* | | *---* |

**Figure 4 – table supplement 2. Statistical data and comparisons between each time-point for female DA1 PNs.**

Table of average Bruchpilot-Short-mStrawberry puncta counts (Brp Mean, red) and average membrane GFP neurite volumes (GFP Mean, green) for each time-point from female DA1 PN time-courses. Mean values are shown with calculated standard error of the mean (SEM) and number of glomeruli analyzed (*n*). Each time-point is compared to each other time-point using one-way ANOVA tests followed by Tukey’s tests to correct for multiple comparisons in determining *p*-values. *p*-values for Brp-Short-mStrawberry puncta are in red and in green for membrane GFP neurite volume.

| DA1 LN Male | Brp Mean +/- SEM (*n*)  GFP Mean +/- SEM (*n*) | | | | | | | vs. 48h | | vs. 60h | | vs. 72h | | | vs. 84h | | vs. 92h | |
| --- | --- | --- | --- | --- | --- | --- | --- | --- | --- | --- | --- | --- | --- | --- | --- | --- | --- | --- |
| 48 h | 27.43 | +/- | | | 5.355 | *(14)* | | *---* | | *.421* | | *<.001* | | | *<.001* | | *<.001* | |
|  | 68.57 | +/- | | | 6.696 | *(14)* | | *---* | | *.001* | | *<.001* | | | *<.001* | | *<.001* | |
| 60 h | 78.44 | +/- | | | 5.682 | *(18)* | | *.421* | | *---* | | *<.001* | | | *<.001* | | *<.001* | |
|  | 160.1 | +/- | | | 8.476 | *(18)* | | *.001* | | *---* | | *.809* | | | *<.001* | | *<.001* | |
| 72 h | 331.2 | +/- | | | 19.54 | *(16)* | | *<.001* | | *<.001* | | *---* | | | *<.001* | | *<.001* | |
|  | 184 | +/- | | | 12.77 | *(16)* | | *<.001* | | *.809* | | *---* | | | *.003* | | *<.001* | |
| 84 h | 750.3 | +/- | | | 29.62 | *(14)* | | *<.001* | | *<.001* | | *<.001* | | | *---* | | *.035* | |
|  | 272.9 | +/- | | | 14.33 | *(14)* | | *<.001* | | *<.001* | | *.003* | | | *---* | | *<.001* | |
| 92 h | 839.1 | +/- | | | 31.12 | *(16)* | | *<.001* | | *<.001* | | *<.001* | | | *.035* | | *---* | |
|  | 396.8 | +/- | | | 28.04 | *(16)* | | *<.001* | | *<.001* | | *<.001* | | | *<.001* | | *---* | |
| DA1 LN Male | Brp Mean +/- SEM (*n*)  GFP Mean +/- SEM (*n*) | | | | | | | vs. 0d | vs. 3d | | vs. 6d | | vs. 9d | vs. 12d | | vs. 15d | | vs. 18d |
| 0 d | 1295 | | +/- | 92.1 | | | *(13)* | *---* | *>.999* | | *>.999* | | *>.999* | *.474* | | *.755* | | *.568* |
|  | 562.3 | | +/- | 26.09 | | | *(13)* | *---* | *.993* | | *.809* | | *.004* | *.012* | | *.228* | | *.137* |
| 3 d | 1355 | | +/- | 73.15 | | | *(13)* | *>.999* | *---* | | *>.999* | | *>.999* | *.752* | | *.947* | | *.813* |
|  | 611.8 | | +/- | 36.92 | | | *(13)* | *.993* | *---* | | *.989* | | *.032* | *.087* | | *.658* | | *.449* |
| 6 d | 1340 | | +/- | 120.1 | | | *(10)* | *>.999* | *>.999* | | *---* | | *>.999* | *.755* | | *.94* | | *.809* |
|  | 670.6 | | +/- | 43.32 | | | *(10)* | *.809* | *.989* | | *---* | | *.308* | *.533* | | *.991* | | *.922* |
| 9 d | 1334 | | +/- | 122.9 | | | *(14)* | *>.999* | *>.999* | | *>.999* | | *---* | *.638* | | *.888* | | *.72* |
|  | 839.9 | | +/- | 60.76 | | | *(14)* | *.004* | *.032* | | *.306* | | *---* | *>.999* | | *.604* | | *.941* |
| 12 d | 1561 | | +/- | AAA | | | *(15)* | *.474* | *.752* | | *.755* | | *.638* | *---* | | *.998* | | *>.999* |
|  | 809.5 | | +/- | 48.16 | | | *(15)* | *.012* | *.087* | | *.533* | | *>.999* | *---* | | *.853* | | *.995* |
| 15 d | 1490 | | +/- | 87.89 | | | *(19)* | *.755* | *.947* | | *.94* | | *.888* | *.998* | | *---* | | *.999* |
|  | 724.3 | | +/- | 57.6 | | | *(17)* | *.226* | *.658* | | *.991* | | *.604* | *.853* | | *---* | | *.999* |
| 18 d | 1561 | | +/- | 88.87 | | | *(11)* | *.568* | *.813* | | *.809* | | *.72* | *>.999* | | *.999* | | *---* |
|  | 761.1 | | +/- | 62.58 | | | *(11)* | *.137* | *.449* | | *.922* | | *.941* | *.995* | | *.999* | | *---* |

**Figure 5 – table supplement 1. Statistical data and comparisons between each time-point for male DA1 LNs.**

Table of average Bruchpilot-Short-mStrawberry puncta counts (Brp Mean, red) and average membrane GFP neurite volumes (GFP Mean, green) for each time-point from male DA1 LN time-courses. Mean values are shown with calculated standard error of the mean (SEM) and number of glomeruli analyzed (*n*). Each time-point is compared to each other time-point using one-way ANOVA tests followed by Tukey’s tests to correct for multiple comparisons in determining *p*-values. *p*-values for Brp-Short-mStrawberry puncta are in red and in green for membrane GFP neurite volume.

| DA1 LN Female | Brp Mean +/- SEM (*n*)  GFP Mean +/- SEM (*n*) | | | | | | | vs. 48h | | vs. 60h | | vs. 72h | | | vs. 84h | | vs. 92h | |
| --- | --- | --- | --- | --- | --- | --- | --- | --- | --- | --- | --- | --- | --- | --- | --- | --- | --- | --- |
| 48 h | 55.19 | +/- | | | 5.182 | *(16)* | | *---* | | *.738* | | *<.001* | | | *<.001* | | *<.001* | |
|  | 78.74 | +/- | | | 5.444 | *(16)* | | *---* | | *.149* | | *<.001* | | | *<.001* | | *<.001* | |
| 60 h | 84.94 | +/- | | | 6.588 | *(18)* | | *.738* | | *---* | | *<.001* | | | *<.001* | | *<.001* | |
|  | 127.2 | +/- | | | 6.795 | *(18)* | | *.149* | | *---* | | *.006* | | | *<.001* | | *<.001* | |
| 72 h | 287.2 | +/- | | | 12.93 | *(20)* | | *<.001* | | *<.001* | | *---* | | | *<.001* | | *<.001* | |
|  | 197.3 | +/- | | | 13.48 | *(20)* | | *<.001* | | *.006* | | *---* | | | *<.001* | | *.004* | |
| 84 h | 520.6 | +/- | | | 32.18 | *(16)* | | *<.001* | | *<.001* | | *<.001* | | | *---* | | *.004* | |
|  | 289.2 | +/- | | | 23.64 | *(16)* | | *<.001* | | *<.001* | | *<.001* | | | *---* | | *.873* | |
| 92 h | 609.2 | +/- | | | 16.63 | *(18)* | | *<.001* | | *<.001* | | *<.001* | | | *.004* | | *---* | |
|  | 269.2 | +/- | | | 16.44 | *(18)* | | *<.001* | | *<.001* | | *.004* | | | *.873* | | *---* | |
| DA1 LN Female | Brp Mean +/- SEM (*n*)  GFP Mean +/- SEM (*n*) | | | | | | | vs. 0d | vs. 3d | | vs. 6d | | vs. 9d | vs. 12d | | vs. 15d | | vs. 18d |
| 0 d | 838.9 | | +/- | 50.77 | | | *(17)* | *---* | *.336* | | *.136* | | *.799* | *>.999* | | *>.999* | | *.132* |
|  | 414.2 | | +/- | 32.09 | | | *(17)* | *---* | *.828* | | *.986* | | *.95* | *.646* | | *.722* | | *.288* |
| 3 d | 1006 | | +/- | 82.93 | | | *(16)* | *.336* | *---* | | *<.001* | | *.988* | *.56* | | *.31* | | *.999* |
|  | 471.6 | | +/- | 18.91 | | | *(16)* | *.828* | *---* | | *.358* | | *>.999* | *>.999* | | *>.999* | | *.973* |
| 6 d | 635.8 | | +/- | 31.01 | | | *(16)* | *.136* | *<.001* | | *---* | | *.002* | *.068* | | *.285* | | *<.001* |
|  | 380.6 | | +/- | 28.56 | | | *(16)* | *.986* | *.358* | | *---* | | *.561* | *.205* | | *.277* | | *.056* |
| 9 d | 946.6 | | +/- | 58.22 | | | *(17)* | *.799* | *.988* | | *.002* | | *---* | *.94* | | *.744* | | *.87* |
|  | 456.6 | | +/- | 38.36 | | | *(17)* | *.95* | *>.999* | | *.561* | | *---* | *.994* | | *.997* | | *.872* |
| 12 d | 864.5 | | +/- | 47.47 | | | *(16)* | *>.999* | *.56* | | *.068* | | *.94* | *---* | | *.999* | | *.274* |
|  | 484.8 | | +/- | 27.74 | | | *(16)* | *.646* | *>.999* | | *.205* | | *.994* | *---* | | *>.999* | | *.997* |
| 15 d | 822.8 | | +/- | 52.15 | | | *(13)* | *>.999* | *.31* | | *.285* | | *.744* | *.999* | | *---* | | *.127* |
|  | 483.4 | | +/- | 38.18 | | | *(13)* | *.722* | *>.999* | | *.277* | | *.997* | *>.999* | | *---* | | *.997* |
| 18 d | 1046 | | +/- | 57.65 | | | *(15)* | *.132* | *.999* | | *<.001* | | *.87* | *.274* | | *.127* | | *---* |
|  | 510.8 | | +/- | 28.11 | | | *(15)* | *.288* | *.973* | | *.056* | | *.872* | *.997* | | *.997* | | *---* |

**Figure 5 – table supplement 2. Statistical data and comparisons between each time-point for female DA1 LNs.**

Table of average Bruchpilot-Short-mStrawberry puncta counts (Brp Mean, red) and average membrane GFP neurite volumes (GFP Mean, green) for each time-point from female DA1 LN time-courses. Mean values are shown with calculated standard error of the mean (SEM) and number of glomeruli analyzed (*n*). Each time-point is compared to each other time-point using one-way ANOVA tests followed by Tukey’s tests to correct for multiple comparisons in determining *p*-values. *p*-values for Brp-Short-mStrawberry puncta are in red and in green for membrane GFP neurite volume.
