## Supplemental Figures for "Different olfactory neuron classes use distinct temporal and molecular programs to complete synaptic development"

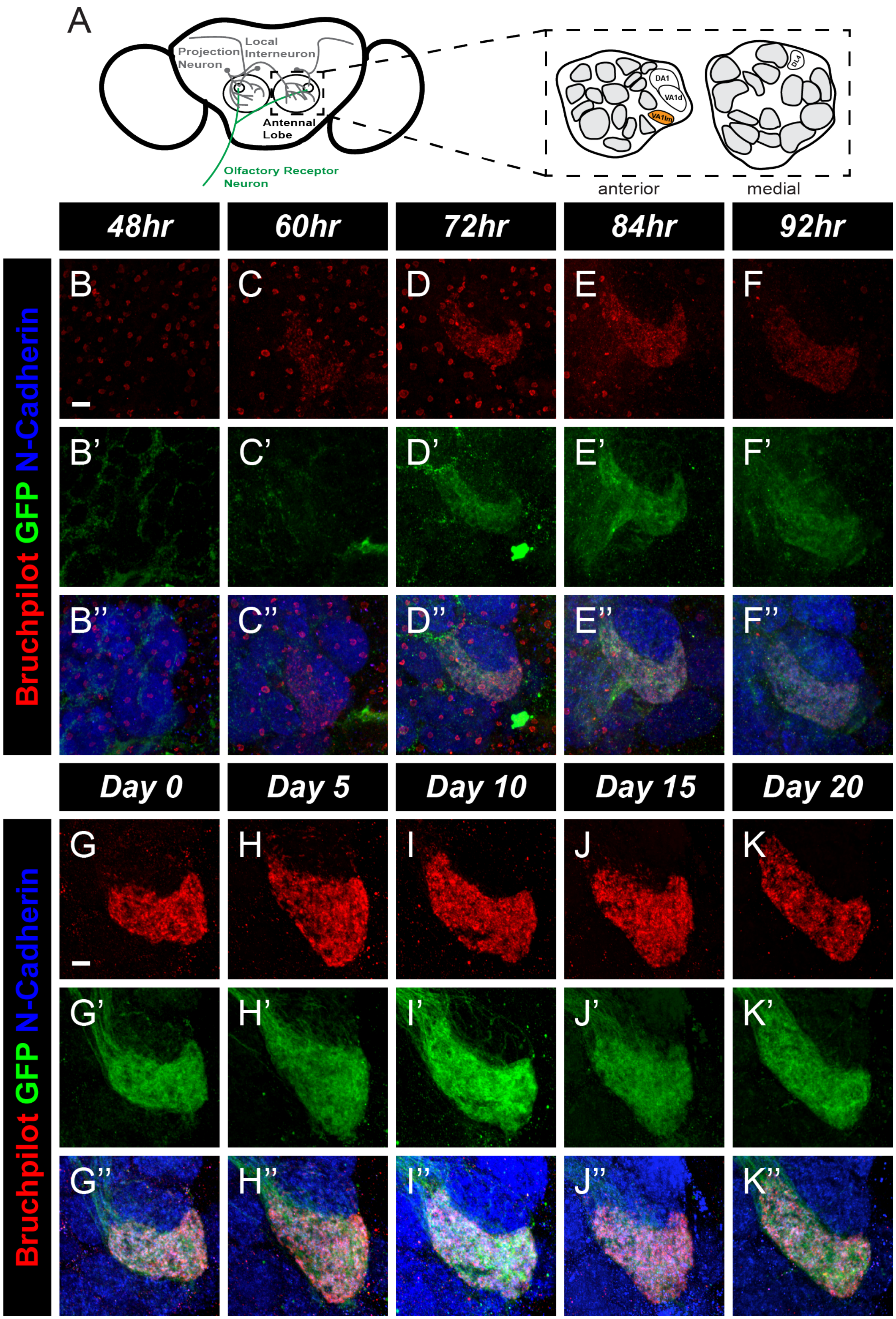


**Figure 2 – figure supplement 1. Synapse formation for female ORNs of the VA1lm glomerulus.** A. Schematic of the Drosophila antennal lobes showing ORNs (green) of the VA1lm glomerulus (orange). B-F'', Representative confocal image stacks of female pupal VA1lm ORNs expressing Brp-Short-mStraw and membrane-tagged GFP and stained with antibodies against mStraw (red), GFP (green), and N-Cadherin (blue) at 48 (B), 60 (C), 72 (D), 84 (E), and 92 (F) hours APF. Note that synaptic puncta do not appear until 60 hours APF. G-K'', Representative confocal image stacks of female adult VA1lm ORNs expressing Brp-Short-mStraw and membrane-tagged GFP and stained with antibodies as in B-F'' at 0 (G), 5 (H), 10 (I), 15 (J), and 20 (K) days post eclosion.

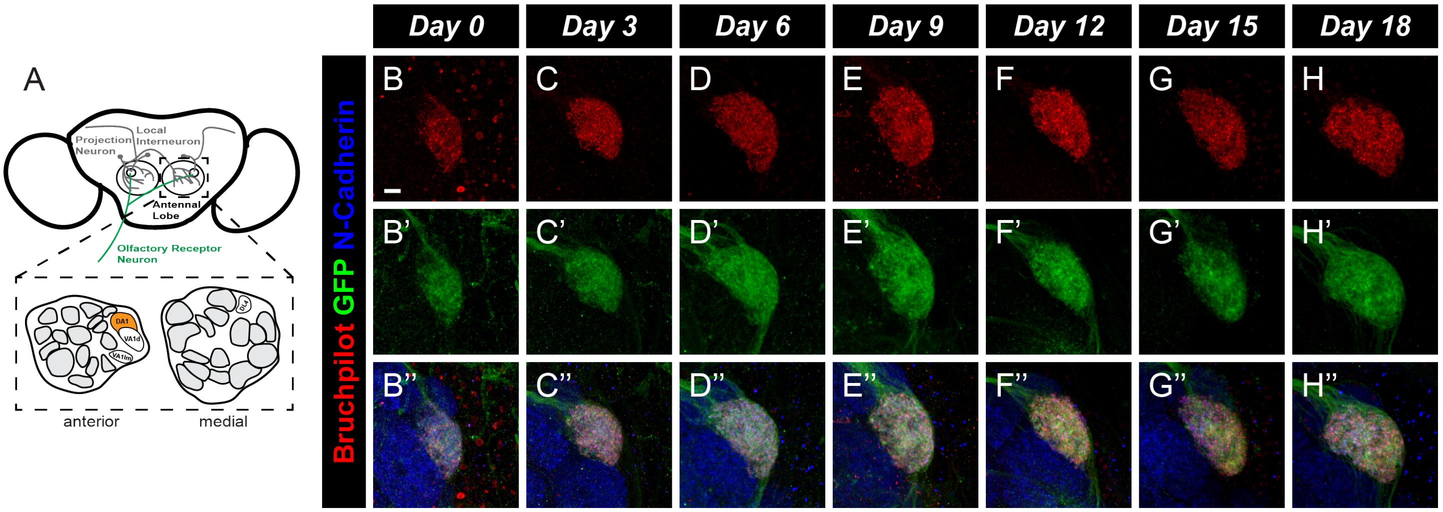


**Figure 3 – figure supplement 1. Synapse formation for the DA1 glomerulus in female ORNs.** A. Schematic of the antennal lobes showing ORNs (green) of the DA1 glomerulus (orange). B-H'', Representative confocal image stacks of female adult DA1 ORNs expressing Brp-Short-mStraw and membrane-tagged GFP and stained with antibodies against mStraw (red), GFP (green), and N-Cadherin (blue) at 0 (B), 3 (C), 6 (D), 9 (E), 12 (F), 15 (G), and 18 (H) days of age.

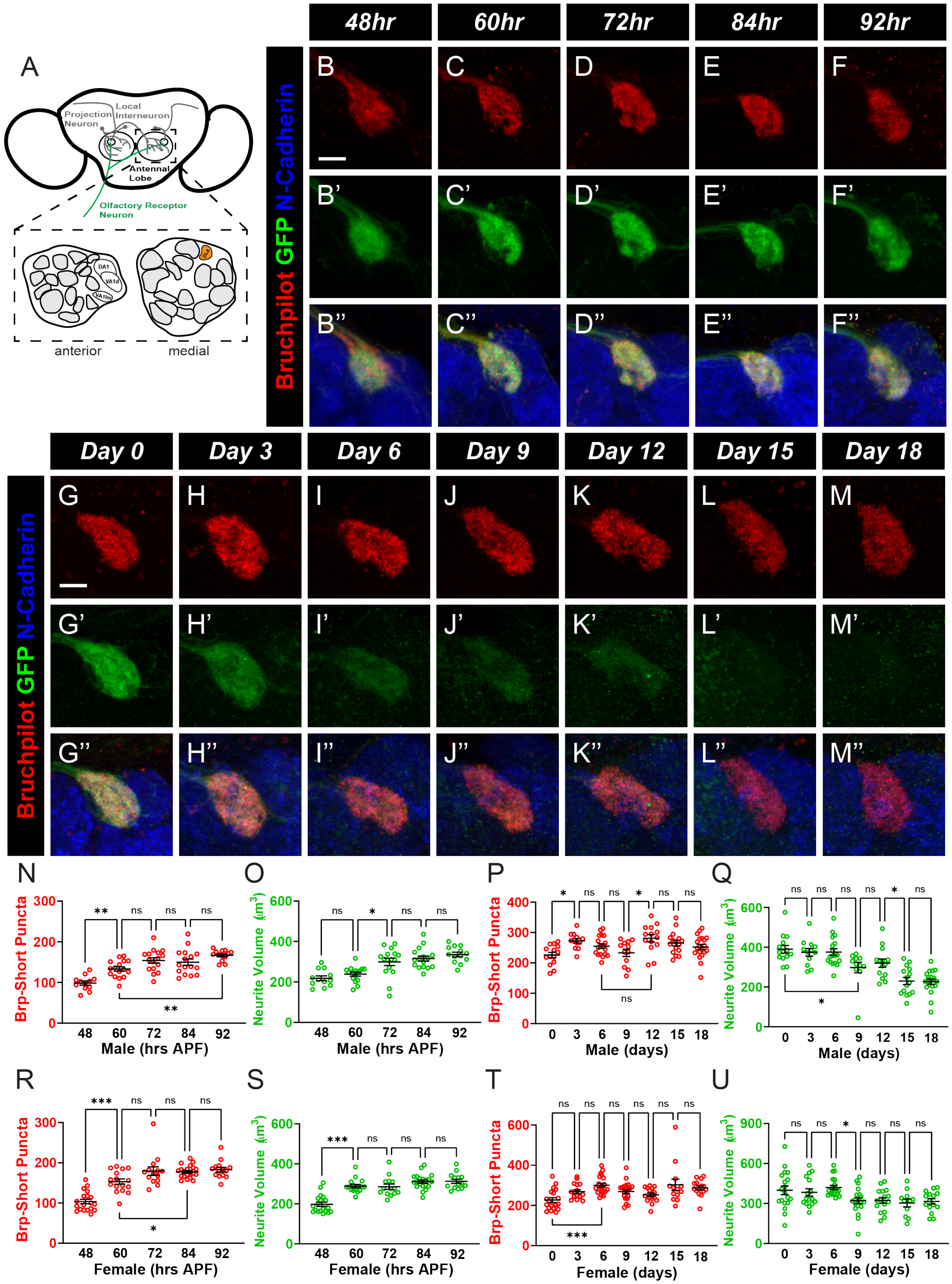

**Figure 3 – figure supplement 2. A developmental time course of synapse number and neurite volume in DL4 ORNs.** A. Schematic of the Drosophila antennal lobes showing ORNs (green) of the DL4 glomerulus (orange). B-F'', Representative confocal image stacks of male pupal DL4 ORNs expressing Brp-Short-mStraw and membrane-tagged GFP and stained with antibodies against mStraw (red), GFP (green), and N-Cadherin (blue) at 48 (B), 60 (C), 72 (D), 84 (E), and 92 (F) hours APF. G-M'', Representative confocal image stacks of male adult D4 ORNs expressing Brp-Short-mStraw and membrane-tagged GFP and stained with antibodies as in B-F'' at 0 (G), 3 (H), 6 (I), 9 (J), 12 (K), 15 (L), and 18 (M) days of age. N-O, Quantification of Brp-Short-mStraw puncta (N) and membrane GFP volume (O) for pupal male DL4 ORNs. Both synaptic puncta and neurite volume increase throughout pupal development. P-Q, Quantification of synaptic puncta (P) and neurite volume (Q) for adult male DL4 ORNs. Synaptic puncta increase from 0 to 3 days of age and then stabilize. Neurite volume steadily decreases over time. R-S, Quantification of Brp-Short-mStraw puncta (R) and membrane GFP volume (S) for pupal female DL4 ORNs. Both synaptic puncta and neurite volume increase from 48 to 60 hours APF. They then trend upward for the remainder of pupal development. T-U, Quantification of synaptic puncta (T) and neurite volume (U) for adult female DL4 ORNs. Synapses trend upward from 0 to 6 days old and then remain largely stable. Neurite volume remains steady until 12 days of age and then begins to decline. For each time-point, n ≥ 11 glomeruli from 6 brains. *, $p < 0.1$, **, $p < 0.01$, ***, $p < 0.001$, n.s. = not significant. Scale bar = 5 μm.

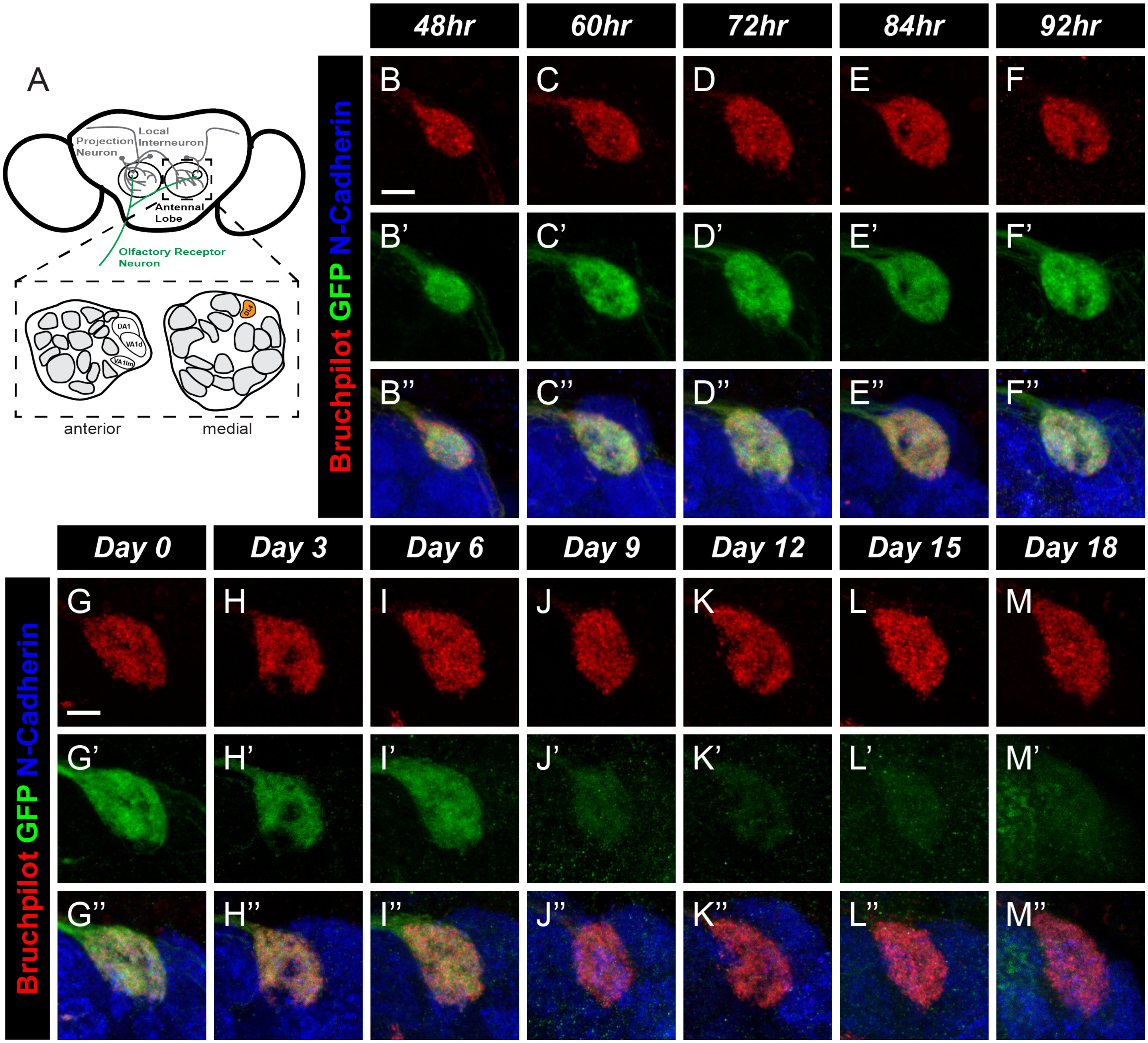


**Figure 3 – figure supplement 3. A developmental time course of synapse number and neurite volume in female DL4 ORNs.** A. Schematic of the Drosophila antennal lobes showing ORNs (green) of the DL4 glomerulus (orange). B-F'', Representative confocal image stacks of female pupal DL4 ORNs expressing Brp-Short-mStraw and membrane-tagged GFP and stained with antibodies against mStraw (red), GFP (green), and N-Cadherin (blue) at 48 (B), 60 (C), 72 (D), 84 (E), and 92 (F) hours APF. G-M'', Representative confocal image stacks of female adult DL4 ORNs expressing Brp-Short-mStraw and membrane-tagged GFP and stained with antibodies as in B-F'' at 0 (G), 3 (H), 6 (I), 9 (J), 12 (K), 15 (L), and 18 (M) days of age.

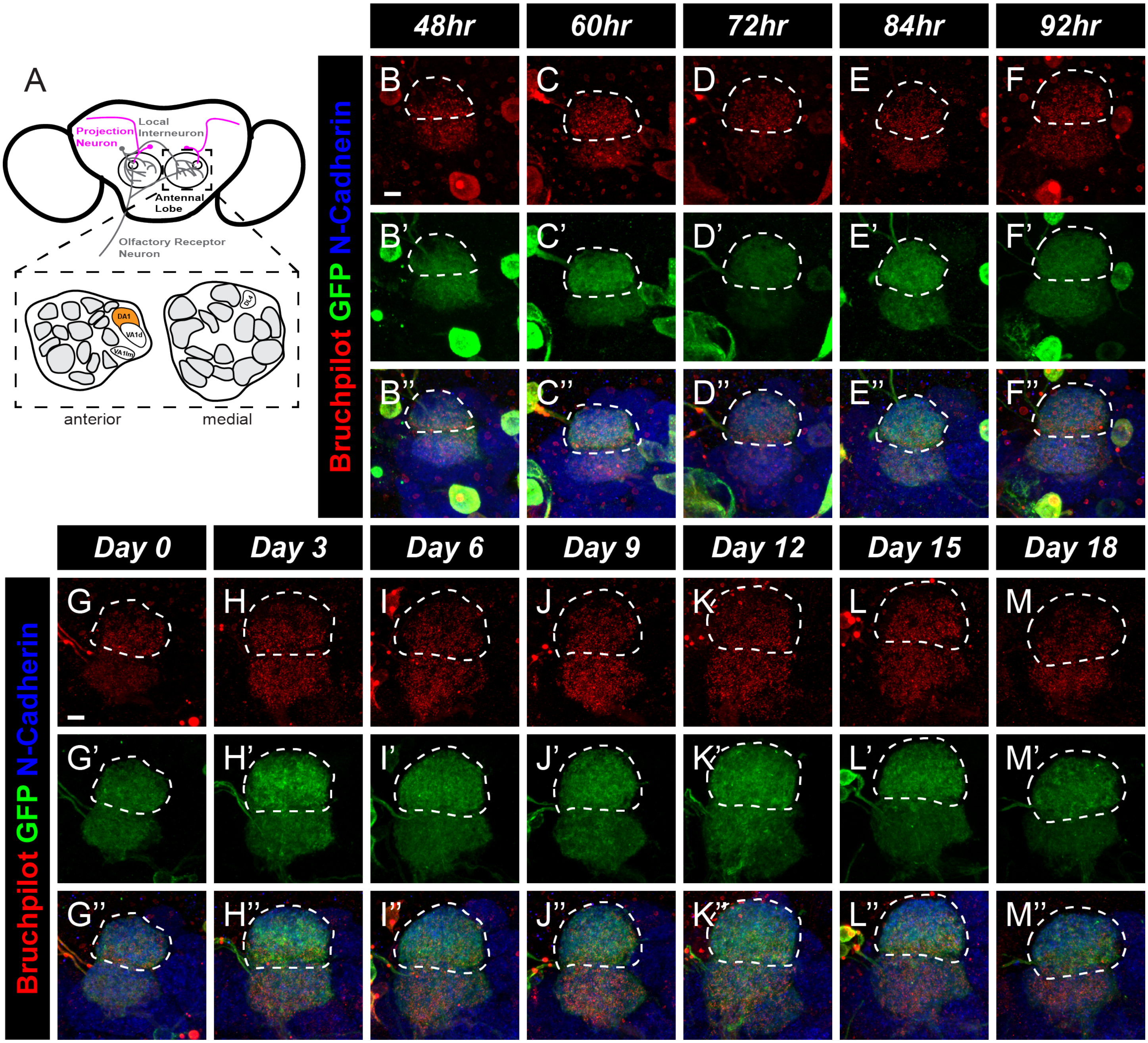


**Figure 4 – figure supplement 1. Synapse formation for female PNs of the DA1 glomerulus.** A. Schematic of the antenna lobes showing PNs (magenta) of the DA1 glomerulus (orange). B-F'', Representative confocal image stacks of female pupal DA1 PNs (dashed white lines) expressing Brp-Short-mStraw and membrane-tagged GFP and stained with antibodies against mStraw (red), GFP (green), and N-Cadherin (blue) at 48 (B), 60 (C), 72 (D), 84 (E), and 92 (F) hours APF. G-M'', Representative confocal image stacks of female adult DA1 PNs (dashed white lines) expressing Brp-Short-mStraw and membrane-tagged GFP and stained with antibodies as in A-E at 0 (G), 3 (H), 6 (I), 9 (J), 12 (K), 15 (L), and 18 (M) days of age.

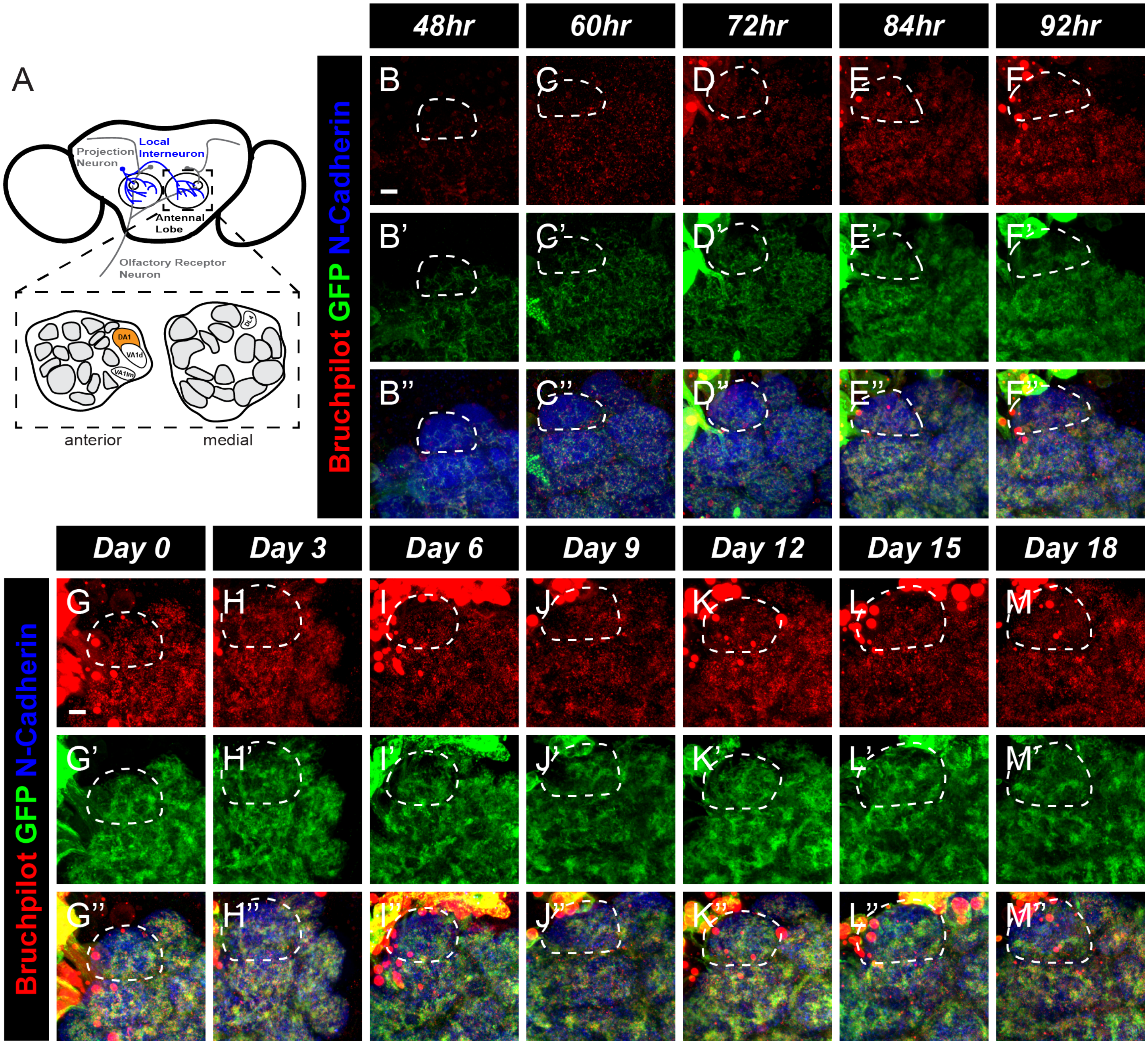


**Figure 5 – figure supplement 1. Synapse formation in the DA1 glomerulus for female LNs.** A. Schematic of the antennal lobes showing LNs (blue) of the DA1 glomerulus (orange). B-F”, Representative confocal image stacks of female pupal DA1 LNs (dashed white lines) expressing Brp-Short-mStraw and membrane-tagged GFP and stained with antibodies against mStraw (red), GFP (green), and N-Cadherin (blue) at 48 (B), 60 (C), 72 (D), 84 (E), and 92 (F) hours APF. G-M”, Representative confocal image stacks of female adult DA1 LNs (dashed white lines) expressing Brp-Short-mStraw and membrane-tagged GFP and stained with antibodies as in B-F” at 0 (G), 3 (H), 6 (I), 9 (J), 12 (K), 15 (L), and 18 (M) days of age.

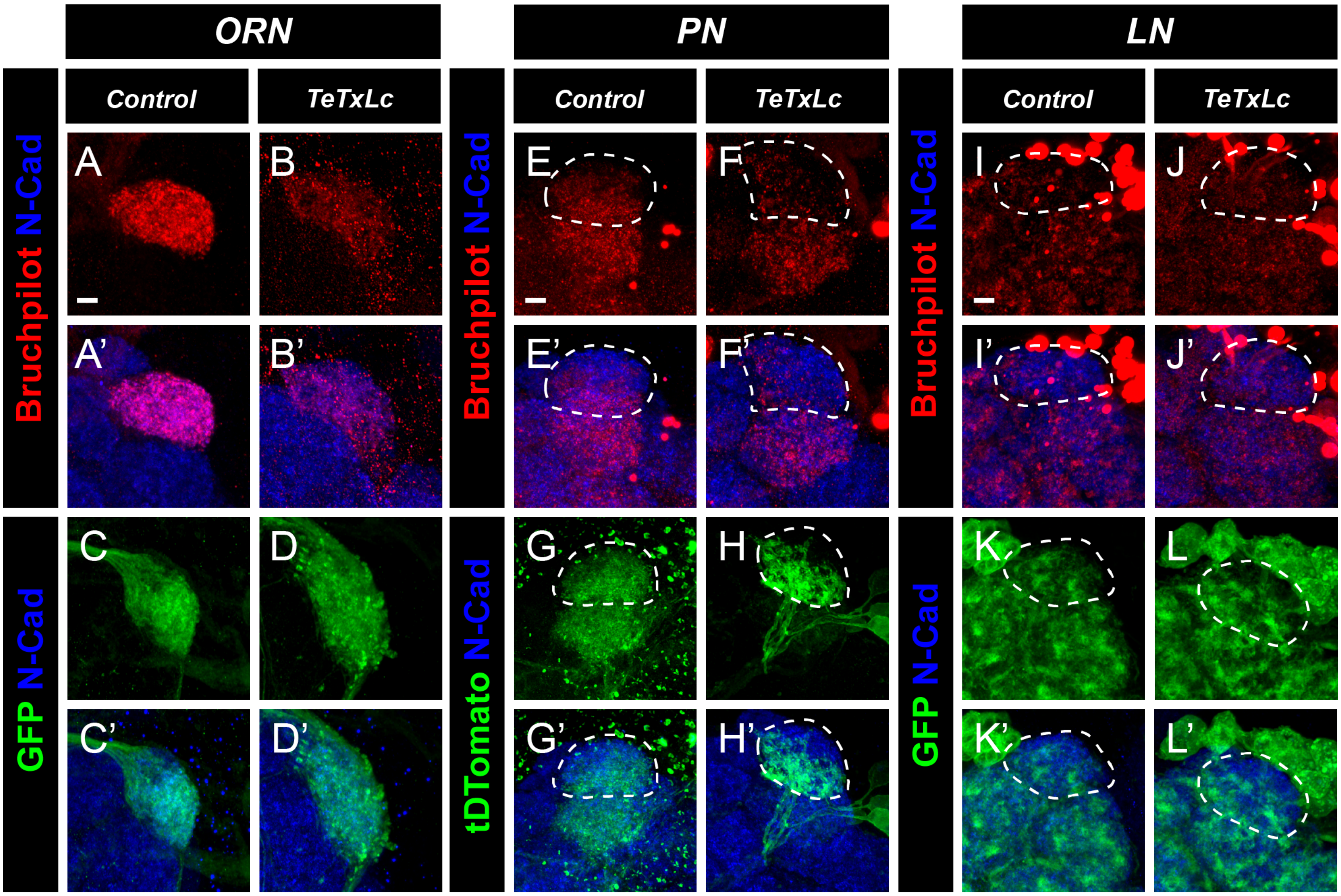


**Figure 6 – figure supplement 1. Decreasing neuronal activity in female antennal lobe neurons of the DA1 glomerulus.** A-D', Representative confocal image stacks of ORNs of the DA1 glomerulus expressing either Brp-Short-mStraw (A-B) or a membrane-tagged GFP (C-D) and either an inactive (A, C) or active (B, D) variant of tetanus toxin light-chain (TeTxLc. Brains were stained with antibodies against mStraw (red) or GFP (green) and N-Cadherin (blue). E-H', Representative image stacks of 10-day old DA1 PNs (dashed white lines) expressing Brp-Short-GFP (E-F) or membrane-tagged tdTomato (G-H) and either inactive (E, G) or active (F, H) tetanus toxin. Brains were stained with antibodies against tdTomato (green) or GFP (red) and N-Cadherin (blue). I-L', Representative image stacks of 10-day old DA1 LNs (dashed white lines) expressing Brp-Short-mStraw (I-J) or membrane-tagged GFP (K-L) as well as inactive (I, K) or active (J, L) tetanus toxin. Brains were stained with antibodies as in A-D'.

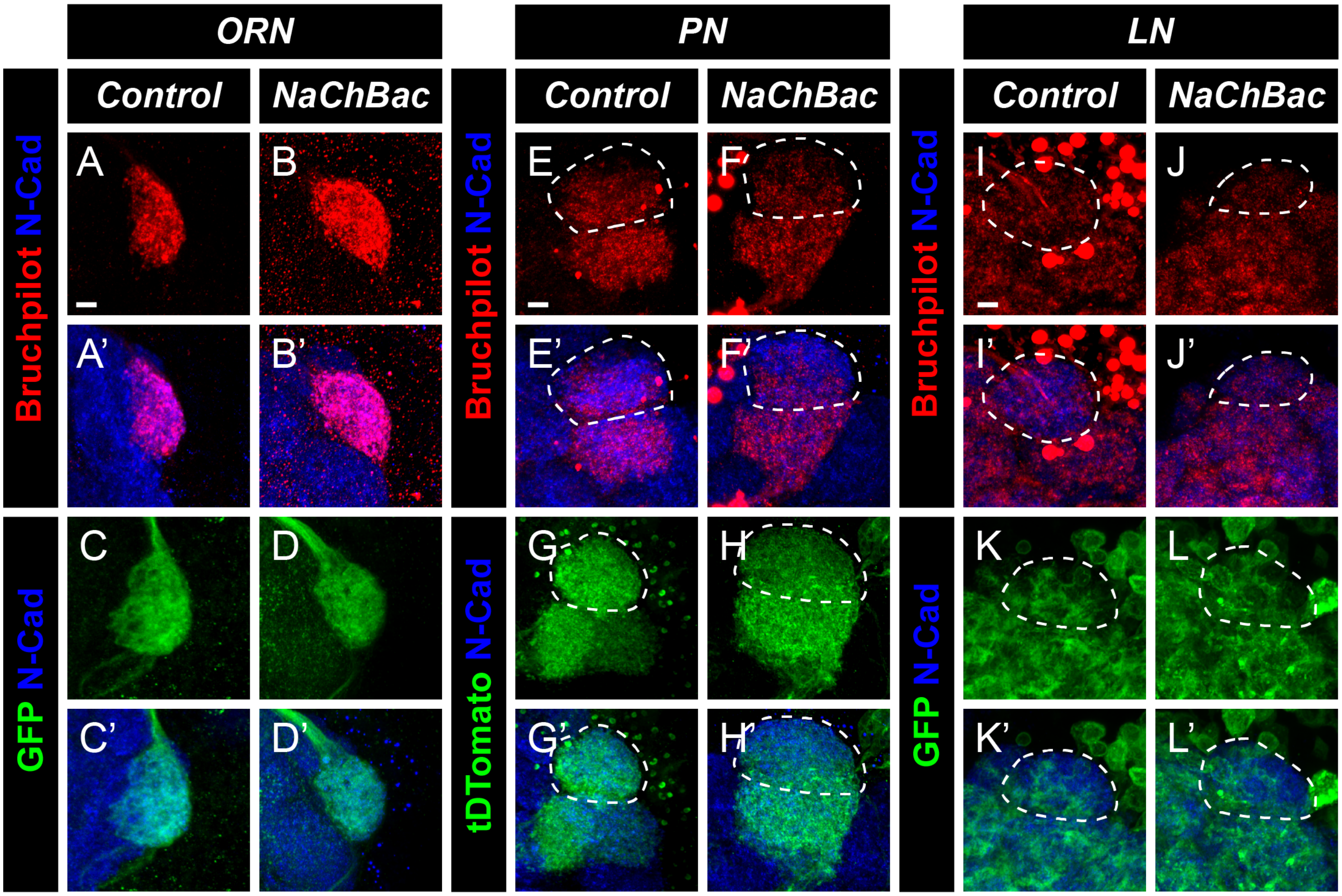


**Figure 7 – figure supplement 1. Increasing neuronal activity in female antennal lobe neurons of the DA1 glomerulus.** A-D', Representative confocal image stacks of 10-day old DA1 ORNs expressing either Brp-Short-mStraw (A-B) or membrane-tagged GFP (C-D) in control flies (A, C) or in flies expressing a NaChBac (B, D) transgene to increase neuronal activity. Brains were stained with antibodies against mStraw (red) or GFP (green) and N-Cadherin (blue). E-H', Representative image stacks of 10-day old DA1 PNs (dashed white lines) expressing Brp-Short-GFP (E-F) or membrane-tagged tdTomato (G-H) in control (E, G) or NaChBac expressing (F, H) flies. Brains were stained with antibodies against tdTomato (green) or GFP (red) as well as N-Cadherin (blue). I-L', Representative image stacks of 10-day old DA1 LNs (dashed white lines) expressing Brp-Short-mStraw (I-J) or membrane-tagged GFP (K-L) in control (I, K) or NaChBac expressing (J, L) flies with antibody staining as in A-D'.

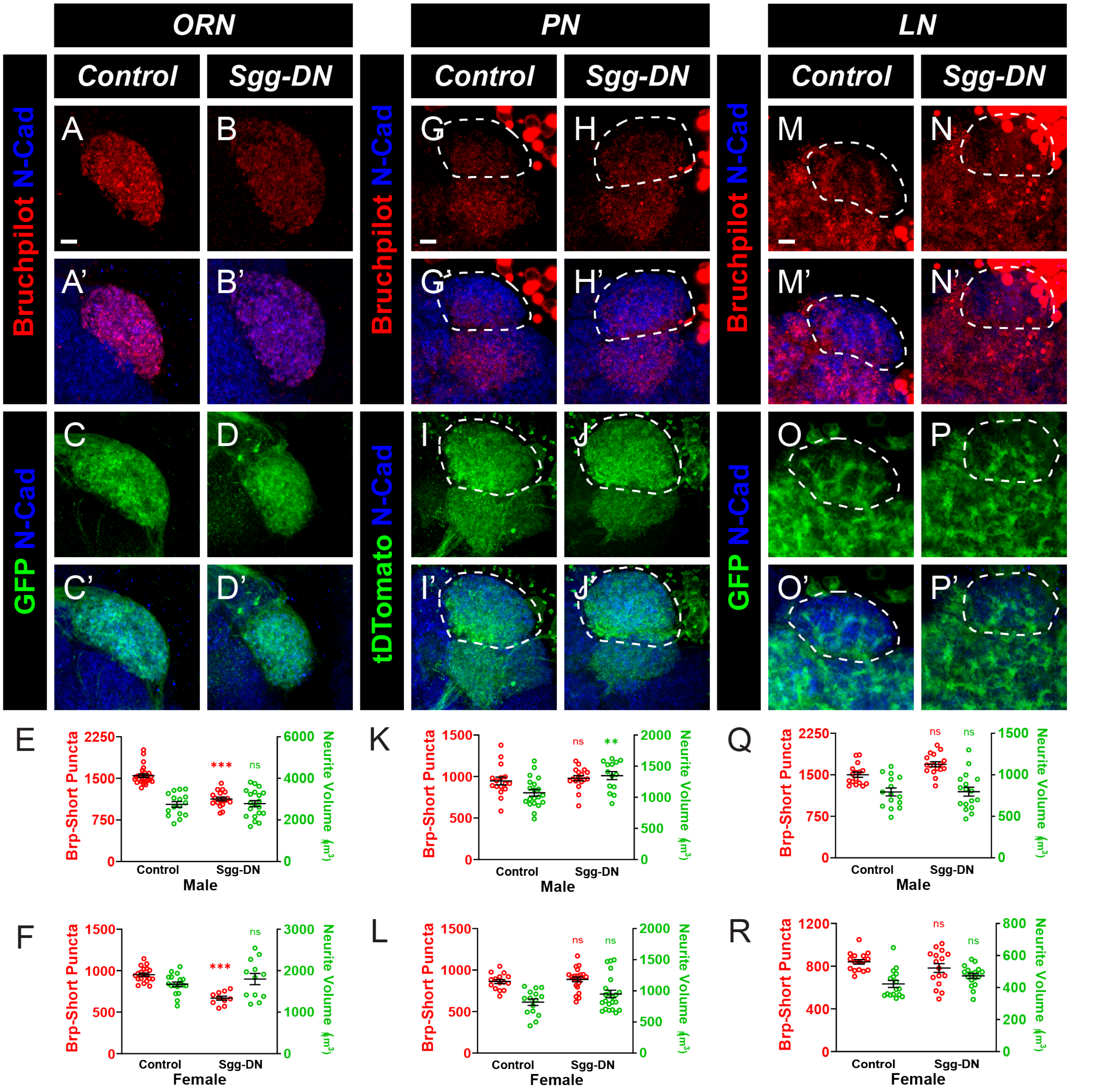


**Figure 8 – figure supplement 1. Decreasing kinase activity in male olfactory neurons decreases synapse number in ORNs, but not PNs or LNs.** A-D', Representative confocal image stacks of 10 day old male ORNs of the DA1 glomerulus expressing Brp-Short-mStraw (A-B') or membrane-tagged GFP (C-D') in control flies (A, C) or in flies expressing a dominant negative (C, D) variant of Shaggy (GSK3β) to decrease overall kinase activity. Brains were stained with antibodies against mStraw (red) or GFP (green) and N-Cadherin (blue). E, Quantification of Brp-Short synaptic puncta and membrane GFP volume in male DA1 ORNs of each group from A-D'. Decreasing kinase activity in ORNs caused a significant decrease in synaptic puncta but did not affect neurite volume. F, Quantification of Brp-Short synaptic puncta and membrane GFP volume in female DA1 ORNs. Decreasing kinase activity in ORNs caused a significant decrease in synaptic puncta but did not affect neurite volume. G-J', Representative image stacks of 10-day old male DA1 PNs (dashed white lines) expressing Brp-Short-GFP (G-H) or membrane-tagged tdTomato (I-J) in control flies (G, I) or in flies expressing dominant negative (H, J) Shaggy. Brains were stained with antibodies against tdTomato (green) or GFP (red) and N-Cadherin (blue). K, Quantification of Brp-Short puncta and neurite volume in male DA1 PNs from G-J'. Decreasing kinase activity did not influence puncta number in PNs but did increase neurite volume. L, Quantification of Brp-Short puncta and neurite volume in female DA1 PNs. Decreasing kinase activity did not alter puncta number or neurite volume in PNs. M-P', Representative confocal image stacks of 10-day old male DA1 LNs (dashed white lines) expressing Brp-Short-mStraw (M-N) or membrane-tagged GFP (O-P) in control flies (M, O) or in flies expressing dominant negative (N, P) Shaggy and stained with antibodies as in A-D'. Q, Quantification of Brp-Short puncta and neurite volume for male DA1 LNs of the groups described in M-P'. Decreased kinase activity did not affect synaptic puncta or neurite volume in LNs. R, Quantification of Brp-Short puncta and neurite volume for female DA1 LNs. Decreased kinase activity did not affect synaptic puncta or neurite volume in LNs. For each experimental group, n ≥ 10 glomeruli from 5 brains. **, $p < 0.01$, ***, $p < 0.001$, n.s. = not significant. Scale bar = 5 µm.

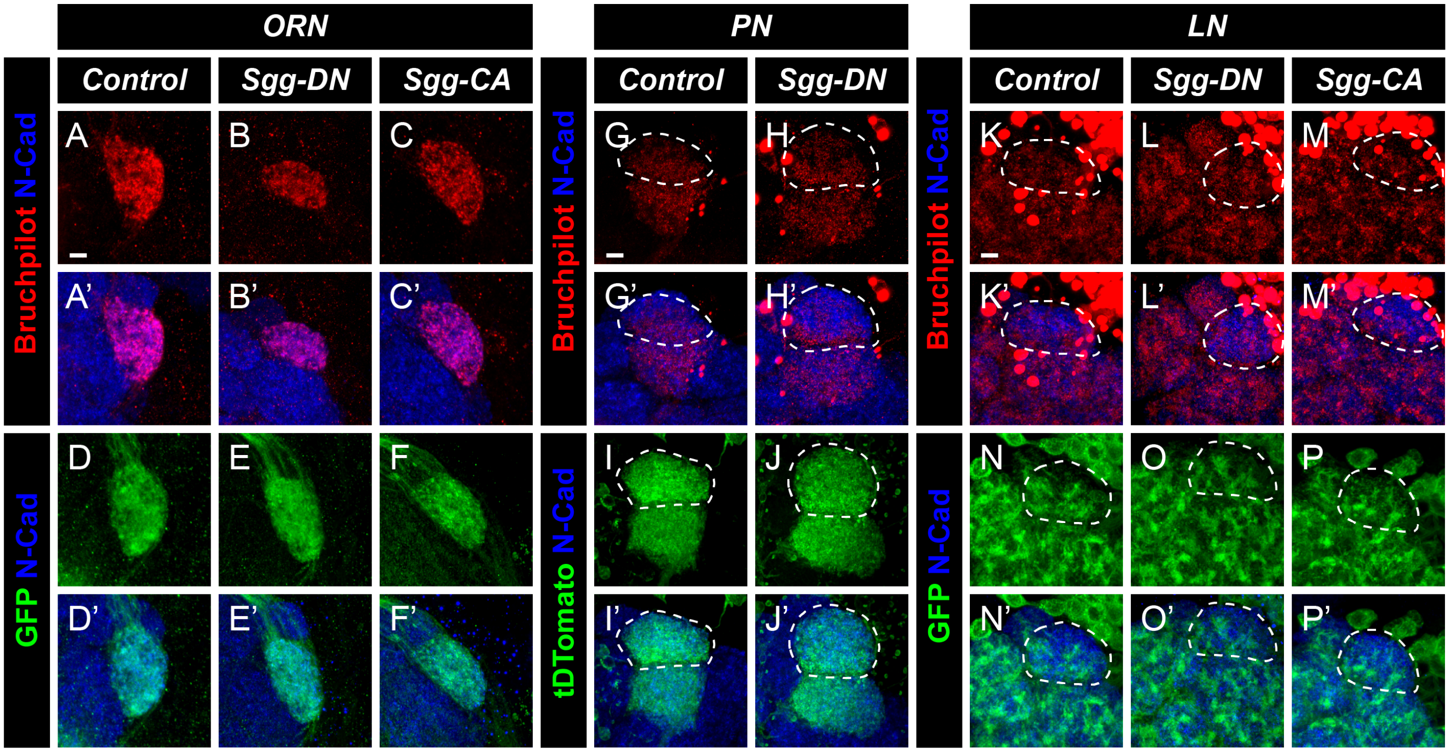


**Figure 8 – figure supplement 2. Altering kinase activity in female olfactory neurons of the DA1 glomerulus.** A-F', Representative confocal image stacks of 10 day old female ORNs of the DA1 glomerulus expressing Brp-Short-mStraw (A-C') or membrane-tagged GFP (D-F') in control flies (A, D) or in flies expressing either a dominant negative (B, E) or a constitutively active (C, F) variant of Shaggy (GSK3β) to decrease or increase overall kinase activity respectively. Brains were stained with antibodies against mStraw (red) or GFP (green) and N-Cadherin (blue). G-J', Representative image stacks of 10-day old DA1 PNs (dashed white lines) expressing Brp-Short-GFP (G-H') or membrane-tagged tdTomato (I-J') in control flies (G, I) or in flies expressing dominant negative (H, J) Shaggy. Brains were stained with antibodies against tdTomato (green) or GFP (red) and N-Cadherin (blue). K-P', Representative confocal image stacks of 10-day old DA1 LNs (dashed white lines) expressing Brp-Short-mStraw (K-M') or membrane-tagged GFP (N-P') in control flies (K, N) or in flies expressing either dominant negative (L, O) or constitutively active (M, P) Shaggy and stained with antibodies as in A-F'.
